# Structural determinants of protocadherin-15 elasticity and function in inner-ear mechanotransduction

**DOI:** 10.1101/695502

**Authors:** Deepanshu Choudhary, Yoshie Narui, Brandon L. Neel, Lahiru N. Wimalasena, Carissa F. Klanseck, Pedro De-la-Torre, Conghui Chen, Raul Araya-Secchi, Elakkiya Tamilselvan, Marcos Sotomayor

## Abstract

Protocadherin-15 (PCDH15), an atypical member of the cadherin superfamily, is essential for vertebrate hearing and its dysfunction has been associated with deafness and progressive blindness. The PCDH15 ectodomain, made of eleven extracellular cadherin (EC1-11) repeats and a membrane adjacent domain (MAD12), assembles as a parallel homodimer that interacts with cadherin-23 (CDH23) to form the tip link, a fine filament necessary for inner-ear mechanotransduction. Here we report X-ray crystal structures of a PCDH15 + CDH23 heterotetrameric complex and ten PCDH15 fragments that were used to build complete high-resolution models of the monomeric PCDH15 ectodomain. Using molecular dynamics (MD) simulations and validated crystal contacts we propose models for complete PCDH15 parallel homodimers and the tip-link bond. Steered MD simulations of these models predict their strength and suggest conditions in which a multimodal PCDH15 ectodomain can act as a stiff or soft gating spring. These results provide a detailed view of the first molecular steps in inner-ear sensory transduction.

## INTRODUCTION

Inner-ear sensory perception begins when mechanosensitive ion channels^1–8^ are gated by “tip links”, fine protein filaments essential for hearing and balance (Fig. 1A)^9–12^. Tip links are 150 to 180 nm long^13–16^, their integrity is calcium (Ca^2+^)-dependent^10, 17^, and they are made of protocadherin-15 (PCDH15) and cadherin-23 (CDH23)^18–21^, two enormous proteins involved in hereditary deafness^22, 23^, progressive blindness^24^, and cancer^25–29^. Mature tip links are thought to be heterotetramers formed by parallel (*cis*) homodimers of PCDH15 interacting in an antiparallel *trans* mode (tip-to-tip) with *cis* homodimers of CDH23^21^, whereas immature tip links are likely formed by *trans* interactions of two PCDH15 molecules^30, 31^.

**Figure 1.**
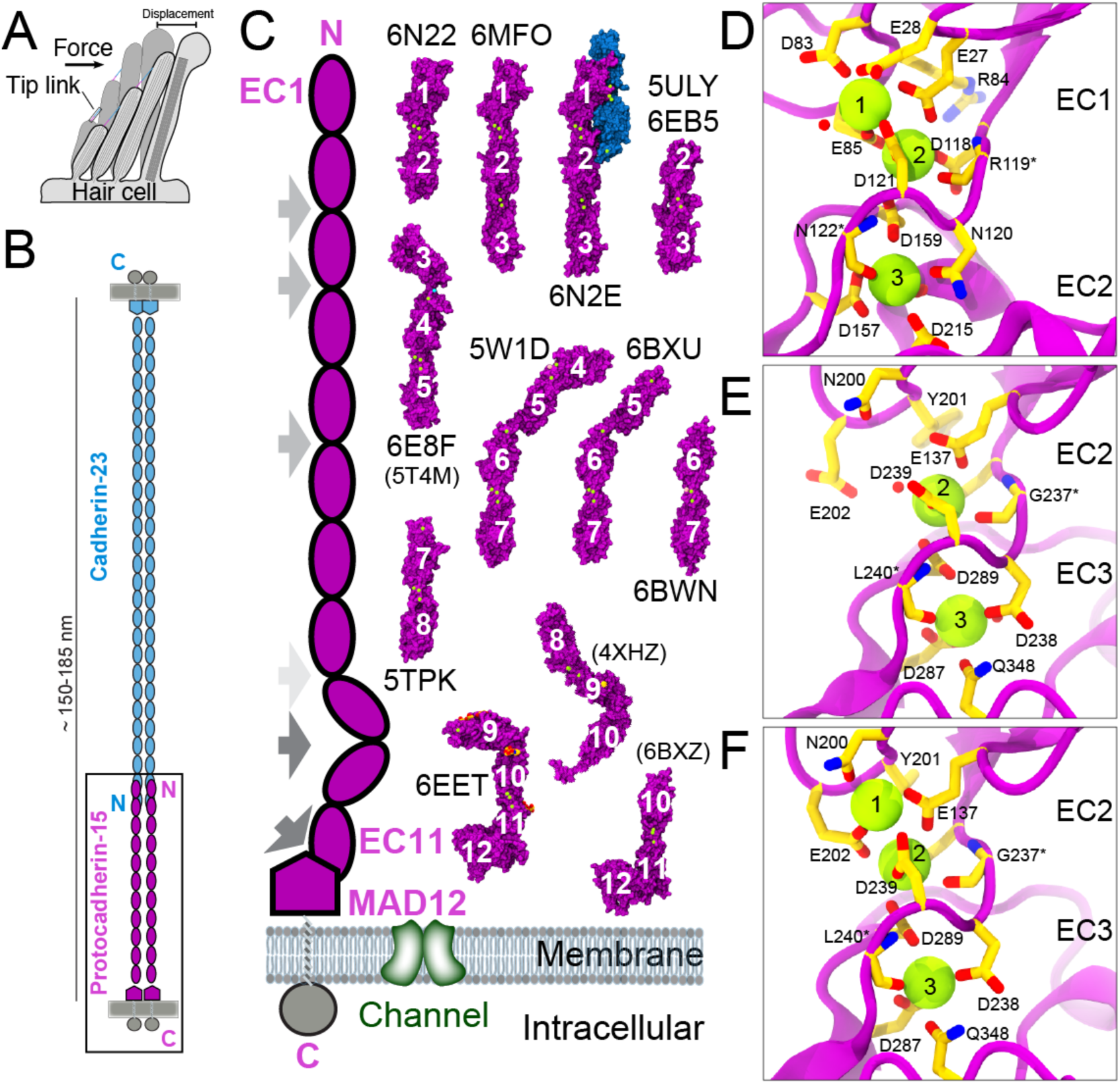
Inner-ear mechanotransduction and PCDH15 structures. (**A**) Hair-cell bundle row showing location of tip links. Force from sound displaces the bundle to activate the transduction apparatus. (**B**) The tip link is made of CDH23 (blue) and PCDH15 (purple). (**C**) Schematic representation of PCDH15 (left panel) and structures of PCDH15 fragments (right) used to build models of the entire PCDH15 ectodomain. PDB codes are indicated for all 11 structures presented here, with codes in parenthesis for three additional structures presented elsewhere^40, 43, 44^. All structures are in surface representation, with PCDH15 fragments in purple and CDH23 EC1-2 in blue. Gray arrows in the schematics indicate EC linkers with atypical canonical-like linker regions (EC8-9 in light gray), with partial Ca^2+^-free linker regions (EC2-3, EC3-4, and EC5-6 in gray), and with Ca^2+^-free linker regions (EC9-10 and EC11-MAD12 in dark gray). (**D**) Detail of a canonical Ca^2+^-binding linker region from *mm* PCDH15 EC1-2BAP. Protein backbone is shown in magenta ribbon. Relevant residues are shown in stick representation and labeled. Asterisk (*) denotes backbone coordination of Ca^2+^. Water molecules are shown as red spheres. Some backbone atoms and some water molecules are omitted for clarity. (**E**) Detail of a partial Ca^2+^-free linker region in *hs* PCDH15 EC2-3 as in D. (**F**) The monomeric mutant *hs* PCDH15 EC2-3 V250N binds all three Ca^2+^ ions. Residue p.E202, previously interacting with p.R254 (not shown), flips in to coordinate Ca^2+^ at sites 1 and 2.

Some details of the PCDH15 and CDH23 interaction are well understood: immunogold electron microscopy (EM) suggests that tip-link lengths are mostly consistent with the predicted length of the combined proteins interacting tip-to-tip^14, 21^; competitive binding of exogenous PCDH15 and CDH23 tip fragments to endogenous proteins blocks regeneration of tip-links and associated transduction currents during hair-cell development and after tip-link rupture with a Ca^2+^ chelator^32^; and structures of the complex formed by the tips of PCDH15 and CDH23 engaged in a heterodimeric molecular “handshake” have been obtained and validated *in vitro* and *in vivo*^33, 34^. Furthermore, mutations that cause deafness in humans and mice have been shown to or are predicted to break the handshake interaction^33–36^. In addition, studies of the CDH23 and PCDH15 heterodimer show that its strength can be tuned by PCDH15 isoforms that have distinct N-terminal tips^37^. The strength of the heterotetrameric bond can be further diversified when considering combinations of PCDH15 isoforms in parallel. However, little was known about the structure of parallel *cis* dimers formed by PCDH15 until recently^38–40^. While negative staining transmission EM showed conformationally heterogeneous parallel dimers for tip-link components^21^, most extracellular fragments of PCDH15 (and CDH23) studied had been monomeric in solution^38, 41–45^. Recent crystals structures of PCDH15 fragments^38–40^ and a low-resolution EM-based model suggest two points of dimerization^38, 39^, yet a detailed atomistic-model of the complete ectodomain is still missing. Also, how PCDH15 engages in *trans* homodimers is unclear, and the structural details and strength of the PCDH15 + CDH23 heterotetrameric bond are not known.

The oligomerization states of PCDH15 and CDH23 and the architecture of their heterotetrameric bond are important determinants of tip-link mechanics and inner-ear mechanotransduction^16, 38^. Disruption of either the complex between PCDH15 and CDH23 or the oligomerization of PCDH15 impairs transduction^33, 34, 38^. The inner-ear transduction channel is gated by a soft element called the “gating spring”, which is either in series with the tip link or the tip link itself^46–48, 6, 8, 49^. Whether cadherin tip links are elastic or rigid has been controversial^11, 16, 42^. The ultrastructure of tip links suggested a stiff elastic element^11^, but the length of tip links vary *in situ*^13, 14^. PCDH15 and CDH23 feature 11 and 27 extracellular cadherin (EC) “repeats” (Fig. 1B), respectively, and initial studies of the structure and simulated dynamics of the CDH23 EC1-2 tip predicted that these canonical repeats and their linker region fully occupied by Ca^2+^ ions at sites 1, 2, and 3 would be stiff^42^ (a similar linker region is shown in Fig. 1D). However, while the EC repeats along PCDH15 and CDH23 are predicted to share a common fold, they vary in sequence, which can result in structural heterogeneity, as seen for other long cadherins^50, 51^. For instance, crystal structures of PCDH15 EC9-10 revealed a bent and L-shaped Ca^2+^-free linker region, with simulations evincing that unbending can provide some elasticity to the tip link^43^. Bending and flexibility at this L-shaped EC9-10 linker region are also observed in low-resolution electron cryo-microscopy (cryo-EM) conformations of a *cis* dimeric PCDH15 fragment encompassing EC8 down to its transmembrane helix^39^. These results highlight the diverse mechanical responses of various tip-link cadherin fragments.

Crystal structures and simulations have also shown that other parts of PCDH15 can be flexible. The PCDH15 EC3-4 linker is flexible and binds two Ca^2+^ ions, instead of three. In addition, this atypical linker binds Ca^2+^ ions with decreased binding affinity (45 μM for site 3 and >100 μM for site 2) as compared to the canonical Ca^2+^-binding linker of CDH23 EC1-2 (5 μM for site 3, 44 μM for site 2, and 71 μM for site 1)^42, 44^. Occupancy of Ca^2+^-binding sites at EC linkers will greatly determine the mechanics of tip links and whether unfolding of EC repeats could occur before unbinding of the handshake bond^33^. Interestingly, bulk endolymphatic Ca^2+^ concentration in the cochlea is tightly regulated and varies along its length from ∼20 μM (base) to ∼40 μM (apex)^52, 53^. However, Ca^2+^ concentration near cochlear tip links could be significantly larger^54, 55^, and vestibular Ca^2+^ concentrations are >100 μM^53, 56^. Yet, the structure and the Ca^2+^-dependent mechanics of entire heterotetrameric tip links remain undetermined. Further issues that need to be investigated include how much Ca^2+^ surrounds tip links *in vivo*^55^, how affinities for Ca^2+^ vary across EC linkers, and how these affinities change when PCDH15 and CDH23 form complexes and when they are under tension.

To better understand the molecular mechanics of tip links and the first steps of inner-ear mechanotransduction, we have determined eleven X-ray crystal structures of PCDH15 fragments (Fig. 1C, Table 1), which show all of PCDH15’s EC repeats, all its EC linkers, and its membrane adjacent domain (MAD12^40^, also referred to as SEA^57^, PICA^38^, or EL^39^). These structures allowed us to assemble a complete model of the monomeric ectodomain of PCDH15, to suggest models of *trans* and *cis* PCDH15 homodimers, and to build a model of the heterotetrameric PCDH15-CDH23 bond. In addition, we used steered molecular dynamics (SMD^58–61^) simulations to predict the Ca^2+^-dependent mechanics of these models, which suggest a multimodal (stiff or soft) elastic response for PCDH15 and indicates some conditions in which PCDH15 can provide both the elasticity and elongation typically associated with the hair-cell gating spring.

**Table 1.**
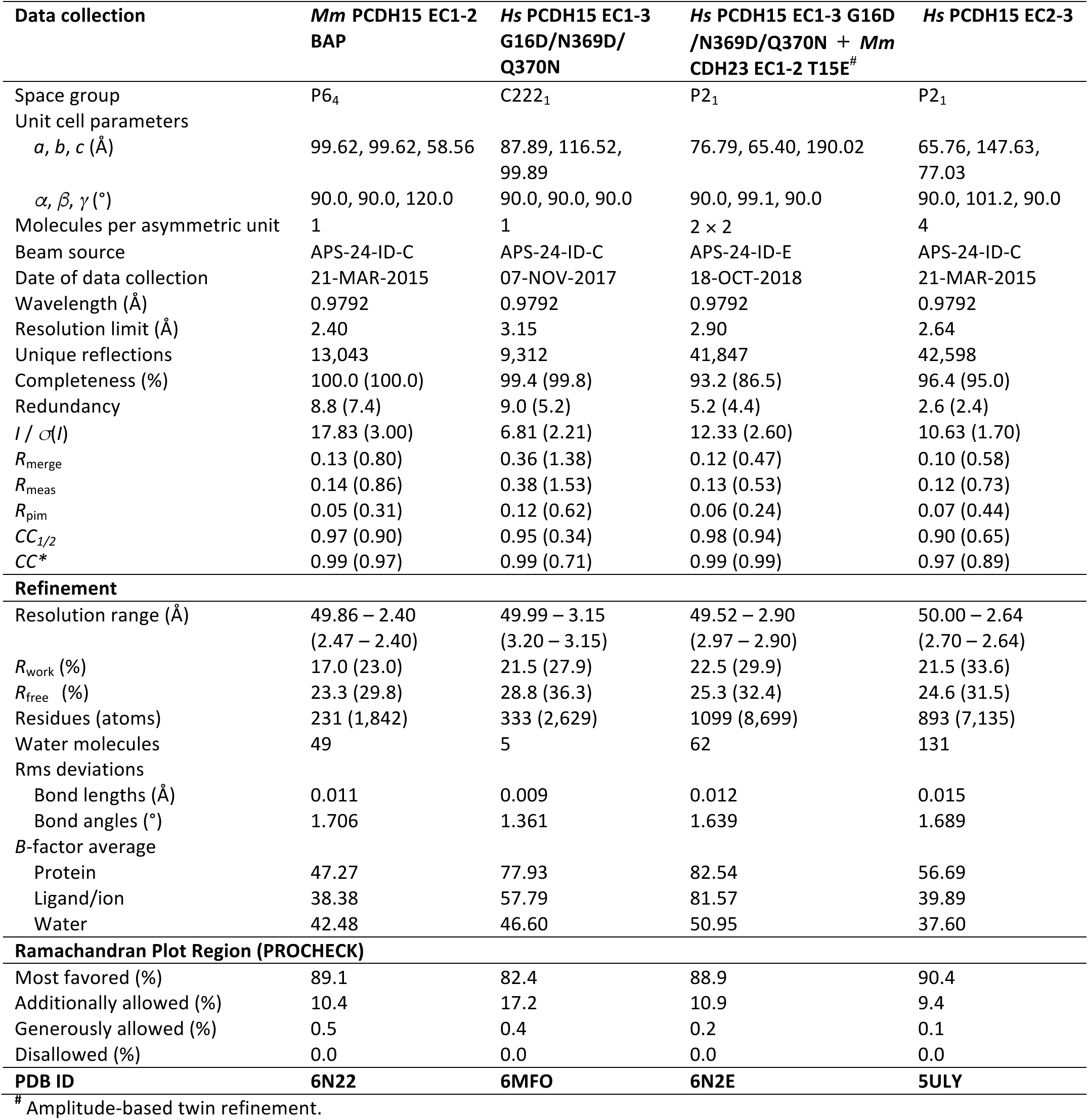
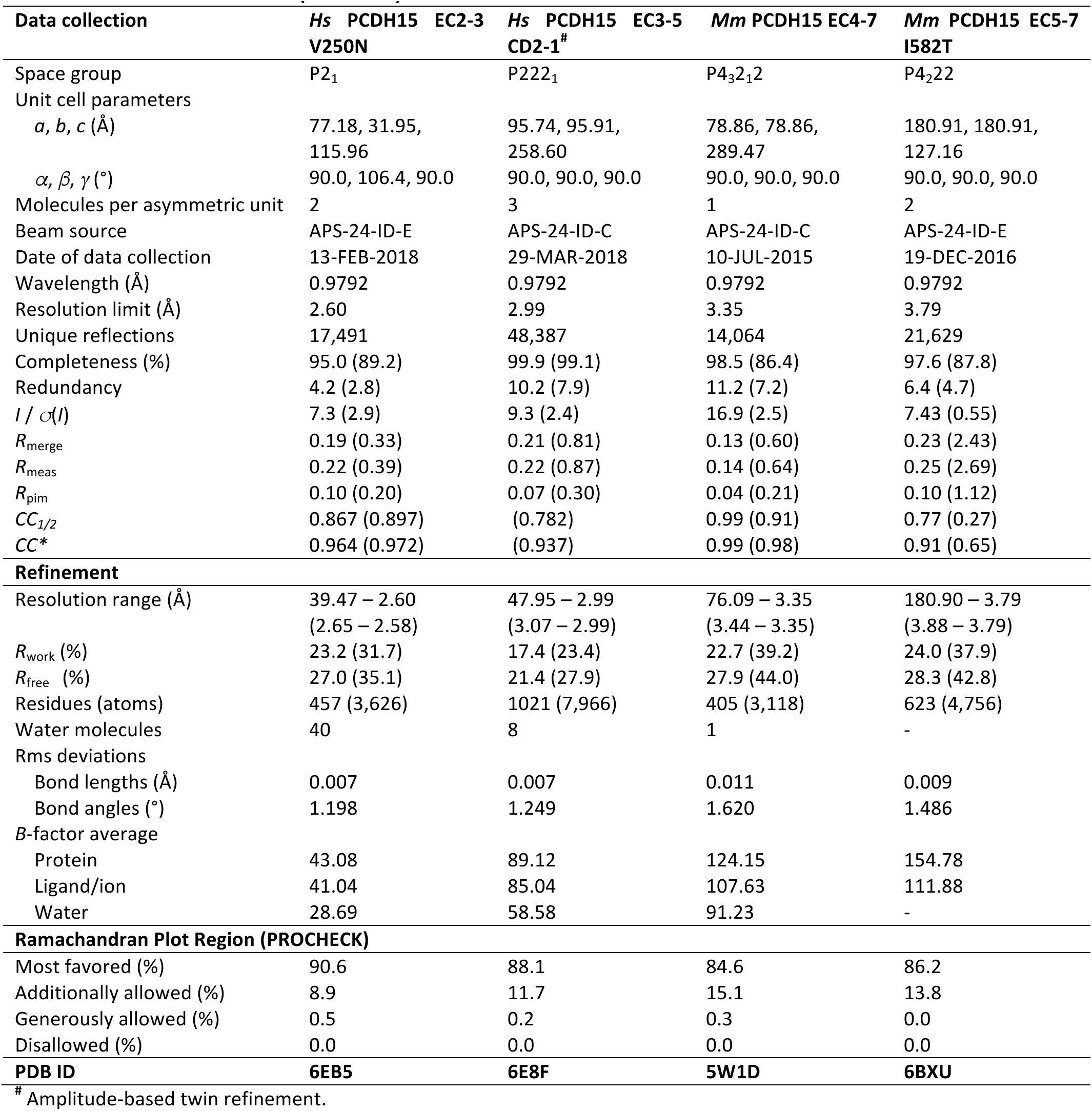
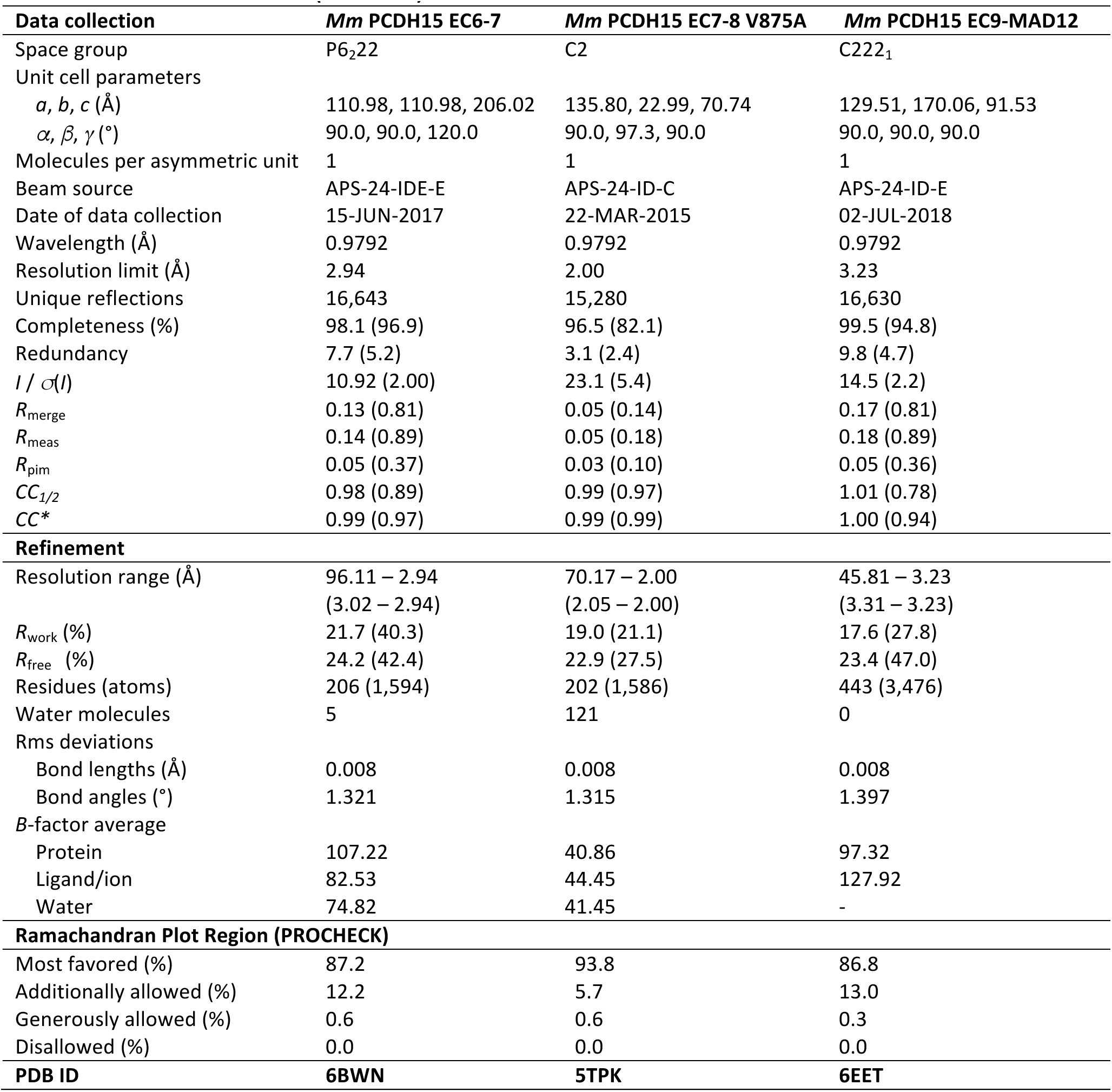
Statistics for structures.

## RESULTS

Sequence analyses of PCDH15 EC repeats and its MAD12 from different species reveal an overall sequence identity of ∼46 ± 9 %, with EC1 and MAD12 being the most conserved (∼59 %) and EC8 being the least (∼32 %). In contrast, comparison among EC repeats (including MAD12) in *Homo sapiens* (*hs*) PCDH15 reveals low overall sequence identity (∼21 %), suggesting structural variability throughout various parts of the protein’s ectodomain within a single species (Figs. S1, S2, and S3; Table S1). Previously published structures of wild-type (WT) *Mus musculus* (*mm*) PCDH15 EC1-2 (bound to *mm* CDH23 EC1-2)^33^, *mm* PCDH15 EC1-3^38^, *hs* PCDH15 EC3-5^44^, *hs* PCDH15 EC8-10^43^, *Sus scrofa* (*ss*) PCDH15 EC10-11+MAD12 (EC10-MAD12)^40^, and *mm* PCDH15 EC11-MAD12^39^ reflect this variability, with unique features seemingly adapted for function in hetero (EC1-2) and homophilic binding (EC2-3 and EC11-MAD12), and in force communication (EC3-5 and EC8-MAD12). Isoform-specific structural variability of PCDH15’s ectodomain also hints at adaptations that might be functionally relevant^37^.

To gain further insights into the function of specific repeats and to build a complete model of PCDH15, we worked with multiple WT and mutated protein fragments (rationale for mutations are explained throughout the text). Successful crystallization and structure determination (Table 1 and Table S2) was possible for *mm* PCDH15 EC1-2 with a biotin acceptor peptide (BAP); *hs* PCDH15 EC1-3 with mutations p.G16D, p.N369D, and p.Q370N (residue numbering in the text and structures corresponds to processed protein, see Methods), alone and in complex with *mm* CDH23 EC1-2 carrying the mutation p.T15E; *hs* PCDH15 EC2-3 (WT and mutant p.V250N); *hs* PCDH15 EC3-5 with exon 12a (isoform CD2-1); *mm* PCDH15 EC4-7; *mm* PCDH15 EC5-7 (p.I582T variant); *mm* PCDH15 EC6-7; and *mm* PCDH15 EC7-8 (p.V875A variant). All these protein fragments were purified from bacterial inclusion bodies under denaturing conditions and refolded (see Methods and Table S3). In addition, we solved the structure of the mammalian-expressed, and thus glycosylated, *mm* PCDH15 EC9-MAD12 fragment. A summary of the protein structures presented is shown in Figure 1A (see also Table 1). Below, we discuss overall properties of EC repeats (from N- to C-terminus) as revealed by these structures, as well as models and simulations of the full-length monomeric and dimeric PCDH15 ectodomain in complex with CDH23 EC1-2 or EC1-3.

### Structures of PCDH15’s ectodomain N-terminal fragments suggest *cis* and *trans* interactions

Individual EC repeat structures and their relative orientation (secondary and tertiary structure features) determine the architecture and shape of entire cadherin ectodomains. While our structures reveal that all PCDH15 repeats (EC1 to EC11) have a typical seven β-strand cadherin Greek-key motif topology^62, 63^ (β strands labeled A to G, Fig. S1), there are some significant variations in loops, secondary structure, linker regions, relative orientation of the EC repeats, and oligomerization states that are likely important for PCDH15’s function in inner-ear mechanotransduction.

The structure of *mm* PCDH15 EC1-2BAP (Fig. 1C) is similar to other structures in which the same protein fragment has been crystallized in complex with *mm* CDH23 EC1-2^33^, or as part of *mm* PCDH15 EC1-3^38^ (core RMSD < 1.5 Å). The *mm* PCDH15 EC1-2BAP fragment is monomeric in solution, and an analysis of the structure’s crystallographic contacts does not reveal possible homophilic interfaces that could mediate PCDH15-PCDH15 *trans* interactions expected for immature tip links^31^. The *mm* PCDH15 EC1-2BAP linker region is canonical (Fig. 1D) and is fully occupied by Ca^2+^ ions at sites 1, 2, and 3, as expected for linker regions with the Ca^2+^-binding motif NTerm-XEX-DXD-D(R/Y)(D/E)-XDX-DXNDN-CTerm^64^. In contrast, the EC2-3 linkers in the structures of the *hs* PCDH15 EC1-3 G16D/N369D/Q370N and *hs* PCDH15 EC2-3 WT fragments (both dimeric in solution) have only two bound Ca^2+^ ions (sites 2 and 3; Fig. 1E and Fig. S4A). A couple of modified Ca^2+^-binding motifs in which DYE is p.200NYE202, and DXNDN is p.236DGDDL240, would suggest that impaired Ca^2+^ binding at site 1 is encoded in the sequence, yet a third structure of a designed mutant *hs* PCDH15 EC2-3 p.V250N that disrupts *cis* homodimerization has three bound Ca^2+^ ions (Fig. 1F and Fig. S4B). While we cannot rule out crystallization conditions, including MgCl_2_ (Table S2), as the source of this discrepancy in ion occupancy, we do observe the p.E202 side chain now coordinating Ca^2+^ at sites 1 and 2 (Fig. S4A,B). These structures suggest that oligomerization can directly or indirectly alter Ca^2+^ binding at the EC2-3 linker, and effect that might be relevant to other cadherin complexes.

Interestingly, the crystal structures of *hs* PCDH15 EC1-3 G16D/N369D/Q370N and *hs* PCDH15 EC2-3 WT reveal crystallographic contacts with arrangements that are compatible with both *trans* and *cis* bonds for PCDH15 (Fig. 2A and Figs. S5A-C and S6A-C). In both cases, the interfaces involve repeats EC2 and EC3. The antiparallel interface observed in *hs* PCDH15 EC1-3 G16D/N369D/Q370N positions EC2 and EC3 from one subunit opposite to EC3 and EC2 from the other (interface area of 919 Å^2^ with an aperture angle of 154.5°; Fig. S5A), thus forming an antiparallel *trans* bond similar to that observed for clustered and δ protocadherins where EC1 and EC4 also play a role in the binding interface^65–69^. The crystallographic contact observed in the *hs* PCDH15 EC2-3 WT structure reveals a potential “X” dimerization interface (∼1,052 Å^2^; Fig. 2A,B) where protomers overlap their linker regions with an aperture angle of 58.75°. In this arrangement the EC2-3 contact forms the hinge of a “molecular scissor” with an estimated *K*_D_ < 1 μM according to analytical ultracentrifugation sedimentation velocity (AUC-SV) experiments (Fig. 2A-C and Fig. S7A). Both interfaces involve the same sides of EC2 and EC3, suggesting that aperture-angle variations due to some relative displacement of the protomers could result in switching between the antiparallel and X-dimer states, with the latter in principle able to mediate both *trans* and *cis* PCDH15-PCDH15 bonds (Fig. S6A-C).

**Figure 2.**
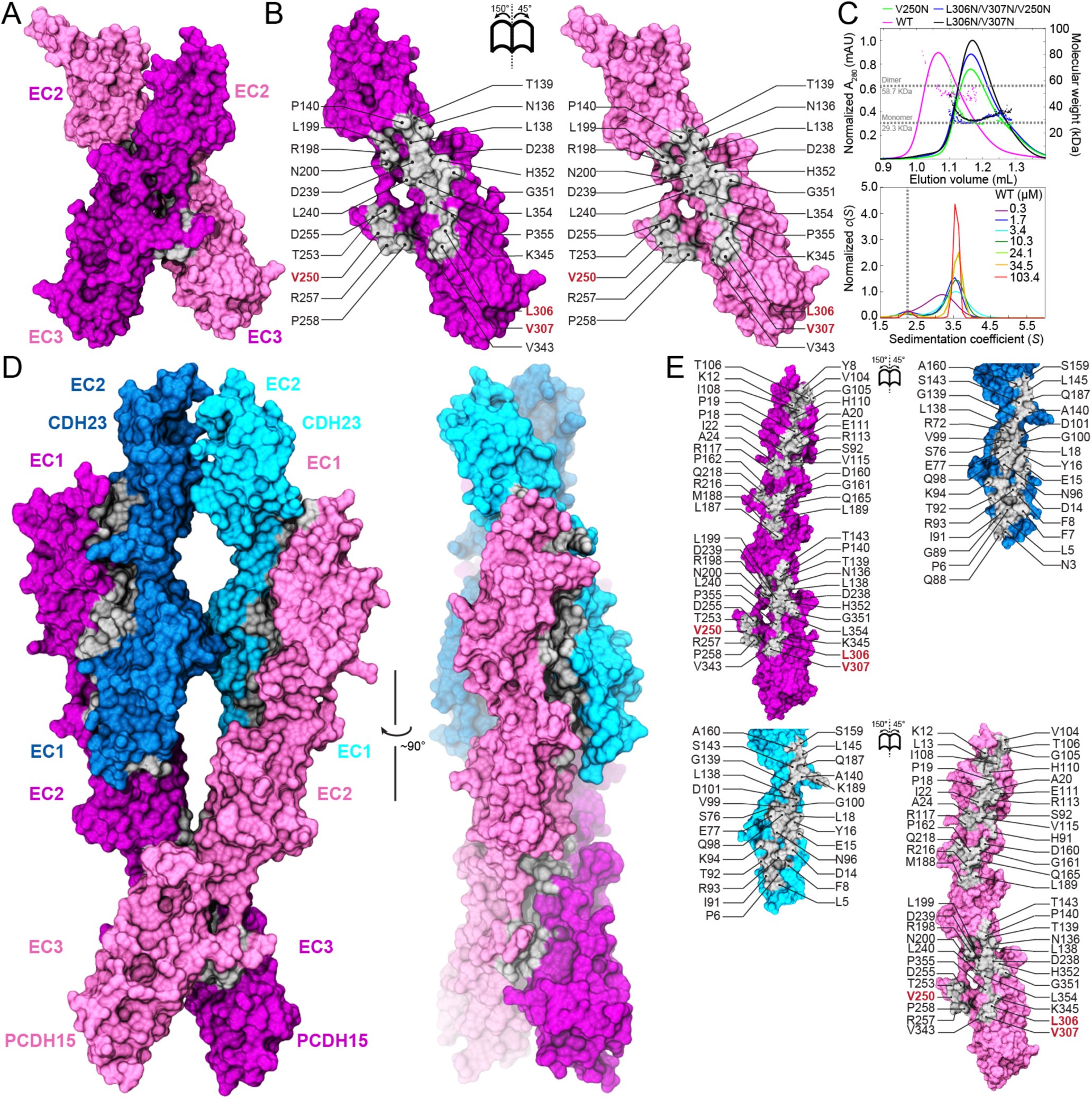
Dimerization of PCDH15 at the EC2-3 linker and the heterotetrameric tip-link bond. (**A**) Front view of the *hs* PCDH15 EC2-3 dimer in surface representation (mauve and purple). Dimer interface is shown in silver. (**B**) Interaction surfaces exposed and colored as in A with interfacing residues labeled. Red labels indicate sites of mutations used to test interface. (**C**) Top panel shows SEC-MALS data for *hs* PCDH15 EC2-3 WT fragment and a few representative mutants. Molecular mass is plotted in colored dots. Horizontal dashed lines indicate theoretical molar mass of 29.3 kDa and 58.7 kDa for monomer and dimer, respectively. Bottom panel shows *c*(*S*) distribution from AUC-SV experiments using *hs* PCDH15 EC2-3 at various concentrations. Peaks at *S* ∼3.5 represent dimers. Vertical dashed line indicates expected position for monomer. Data for mutants is shown in Fig. S7. (**D**) Front and side views of the heterotetrameric *hs* PCDH15 EC1-3 G16D/N369D/Q370N + *mm* CDH23 EC1-2 T15E structure. Molecular surfaces are shown as in A with CDH23 in cyan and blue. (**E**) Interaction surfaces exposed and colored as in D with interfacing residues labeled.

The X-dimer interface we observed in *hs* PCDH15 EC2-3 WT is similar to that reported by Dionne et al.^38^, where a crystal structure of the glycosylated *mm* PCDH15 EC1-3 WT fragment (PDB: 6CV7) shows an X-dimer with an aperture angle of 59.59° (Fig. S8A). Overlapping the glycosylated *mm* PCDH15 EC1-3 WT monomer in the antiparallel configuration of our non-glycosylated *hs* PCDH15 EC1-3 G16D/N369D/Q370N structure reveals potential steric clashes between sugars at glycosylation site p.N180 and the cysteine-stapled loop in EC3 (with poor density in our structure), thus suggesting that a complete antiparallel *trans* state is not feasible unless the involved cysteine loop rearranges (Fig. S5B). It is possible that variations in the angle of the X interface controlled by glycosylation lead to transitions from antiparallel to X-dimer states. Of note, some mutations that disrupt the X-dimer (such as p.V250N and p.V250D^38^ in EC2-3) are also predicted to disrupt the PCDH15-PCDH15 antiparallel dimer, and a monomeric state was observed when the p.V250N mutation was incorporated into our glycosylated *mm* PCDH15 EC1-4 fragment (Table 2 and Fig. S7B). In contrast, the double mutation p.L306N/p.V307N at the EC3-EC3 protomer interface of the X-dimer is not predicted to disrupt the *trans* antiparallel interface (Fig. S5A), but both *hs* PCDH15 EC2-3 p.L306N/p.V307N and *hs* PCDH15 EC1-3 p.L306N/p.V307N are monomers in solution (Table 2). Thus, the antiparallel *trans* PCDH15-PCDH15 interface may use the X-dimer interaction as a required intermediate or might be short lived^70^. While the partially deleterious impact of a similar set of mutations (including p.V250D) on hair-cell mechanotransduction highlights the relevance of EC2-3 contacts for PCDH15 function *in vivo*^38^, an evaluation of how these mutations alter PCDH15-PCDH15 tip links^31^ is still missing.

**Table 2.** Oligomerization state of PCDH15 fragments.

| System | Predicted Mass<br>Monomer / Dimer kDa | SEC-MALS<br>kDa<sup>†</sup> | SEC-SAXS<br>kDa<sup>†</sup> | State |
| --- | --- | --- | --- | --- |
| *Hs* PCDH15 EC2-3 WT (*n* = 3) | 29.3 / 58.7 | 57.2 (2.6%) | -- | Dimer |
| *Hs* PCDH15 EC2-3 V250N (*n* = 3) | 29.3 / 58.7 | 32.4 (10.6%) | -- | Monomer |
| *Hs* PCDH15 EC2-3 L306N/V307N (*n* = 3) | 29.3 / 58.7 | 32.7 (11.6%) | -- | Monomer |
| *Hs* PCDH15 EC2-3 V250/L306N/V307N (*n* = 2) | 29.3 / 58.7 | 30.1 (2.7%) | -- | Monomer |
| *Hs* PCDH15 EC1-3 WT (*n* = 1) | 43.0 / 86.0 | 84.3 (2.0%) | -- | Dimer |
| *Hs* PCDH15 EC1-3 L306N/V307N (*n* = 1) | 43.0 / 86.0 | 46.4 (7.9%) | -- | Monomer |
| *Hs* PCDH15 EC1-4 WT (*n* = 1) | 55.7 / 111.5 | 106.7 (4.3%) | -- | Dimer |
| *Hs* PCDH15 EC1-4 L306N/V307N (*n* = 1) | 55.8 / 111.5 | 57.9 (3.8%) | -- | Monomer |
| *Mm* PCDH15 EC9-MAD12 (*n* = 1) | 51.9 / 103.7 | -- | 98.1 (5.4%) | Dimer |
| *Mm* * PCDH15 EC1-3 WT (*n* = 1) | 43.7 / 87.4 | 105.4 (20.6%) | -- | Dimer |
| *Mm* * PCDH15 EC1-4 WT (*n* = 2) | 56.7 / 113.3 | 139.6 (21.9%) | -- | Dimer |
| *Mm* * PCDH15 EC1-4 V250N (*n* = 2) | 56.7 / 113.3 | 72.6 (26.7%) | -- | Monomer |
| *Mm* * PCDH15 EC9-MAD12 WT (*n* = 1) | 52.9 / 105.7 | 111.9 (5.9%) | -- | Dimer |
| *Mm* * PCDH15 EC1-MAD12 WT (*n* = 1) | 152.4 / 304.8 | 377.1 (23.2%) | -- | Dimer |
| *Mm* * PCDH15 EC1-MAD12 WT EDTA<sup>¶</sup> (*n* = 1) | 152.4 / 304.8 | 355.8 (16.3%) | -- | Dimer |

<sup>†</sup> Values in parenthesis indicate percentage deviation from predicted molecular mass.
\* Mammalian expressed protein. The predicted mass represents that of the protein sequence without glycosylation. The calculated mass from SEC-MALS experiments is higher due to the presence of sugar moieties on the protein.
<sup>¶</sup> EDTA was present in the SEC column but not directly added to the sample.

Using our structure of *hs* PCDH15 EC2-3, we asked if its X-dimeric arrangement would be compatible with the *trans* heterodimeric “handshake” interaction between PCDH15 EC1-2 and CDH23 EC1-2^33^. Alignment of EC2 repeats of PCDH15 from the handshake and X-dimer structures allowed us to build a model of the PCDH15 and CDH23 heterotetramer without steric clashes, suggesting that this is a valid model for the tip-link bond (obtained independently by Dionne et al. using *mm* PCDH15 EC1-3^38^; Fig. S8B). Superposition of the *Danio rerio* (*dr*) CDH23 EC1-3 structure^45^ onto this complex is also compatible with the heterotetrameric bond conformation in which PCDH15 EC2-3 forms an X-dimer interface (Fig. S8C), further suggesting that parallel dimerization in CDH23 involves repeats beyond EC3.

To better understand the structural determinants of the heterotetrameric tip-link configuration and to determine the *cis* configuration of PCDH15 upon complex formation with CDH23, we crystallized higher affinity variants^36^ of tip-link fragments containing repeats EC1 to EC3 of PCDH15 and EC1 to EC2 of CDH23. This structure, comprising two molecules each of *hs* PCDH15 EC1-3 G16D/N369D/Q370N in complex with *mm* CDH23 EC1-2 T15E, provides a complete view of the heterotetrameric tip-link bond (Fig. 2D). The two PCDH15 protomers are seen in an X configuration flanking two molecules of CDH23, thereby forming a “double handshake” consistent with the heterotetrameric model deduced from the *hs* PCDH15 EC2-3 structure and independently proposed by Dionne et al.^38^ (Fig. 2 and Fig. S8), but with novel structural variations. The structure is also compatible with longer fragments of CDH23 (Fig. S9). Interface areas for the two handshakes in our structures are different, with one of them being larger than all the previously published heterodimeric handshakes^33^ (Table S4). The aperture angle for the *hs* PCDH15 dimer is 55.69° in the tetramer, which is smaller than observed for dimeric *mm* PCDH15 EC1-3^38^ and *hs* PCDH15 EC2-3, implying that the “scissor” is less open in the tip-link tetramer structure.

Interestingly, electron density quality is generally good throughout the heterotetrameric *hs* PCDH15 EC1-3 G16D/N369D/Q370N + *mm* CDH23 EC1-2 T15E structure, except for parts of CDH23 EC2 repeats, which were not built (β-strands F and G, Fig. S10A). This is also reflected in larger B factor values across the PCDH15 EC1 and CDH23 EC1-2 repeats, which suggest some inherent flexibility or disorder compared to other regions of the model (Fig. S10). Mutations G16D and T15E had been identified as enhancers of equilibrium binding for the *mm* PCDH15 EC1-2 and *mm* CDH23 EC1-2 complex^36^, and may have helped stabilize a heterotetrameric complex that otherwise requires longer CDH23 fragments to form (mutations p.N369D and p.Q370N complete a canonical DXNDN Ca^2+^-binding motif at the end of PCDH15 EC3 and are not expected to affect structure in any way). Alternatively, the stoichiometry of the complex might not be uniform across the crystal lattice, yet a composite omit 2mF_o_-DF_c_ map generated with simulated annealing suggests that both *mm* CDH23 EC1-2 protomers are present in the crystal (Fig. S10B,C).

Some details of the handshake interactions seen in our heterotetrameric *hs* PCDH15 EC1-3 G16D/N369D/Q370N + *mm* CDH23 EC1-2 T15E structure (Fig. 2D,E) are subtly different when comparing to the single handshake *mm* PCDH15 EC1-2 + CDH23 EC1-2 complex^33^ (PDBs: 4APX and 4AXW in form I; 4AQ8 in form II). In both PCDH15 protomers of the heterotetrameric structure, the side chain of residue p.R113, known to be important for the hetero *trans* tip-link bond and for inner-ear function^21, 32, 33, 71^, interacts with CDH23’s p.Q98 rather than with CDH23’s p.E77 (Fig. S11A). A similar arrangement was seen in the structure of *mm* PCDH15 EC1-2 isoform CD1-2 bound to *mm* CDH23 EC1-2^37^ (PDB: 4XXW). In addition, the CDH23 EC1 repeat of one of the protomers is closer to its PCDH15 EC2 partner (Fig. S11A,B), with PCDH15 EC1-2 and CDH23 EC1-2 inter-repeat angles varying across chains and structures (Fig. S8), although this flexing has been observed across previously published structures of the *mm* PCDH15 EC1-2 + CDH23 EC1-2 complex^33^.

In addition, in our heterotetramer structure, the α helices between β-strands C and D of CDH23 EC1 repeats are significantly closer to each other in the crystal structure (Fig. S11C) as compared to the heterotetramer model generated from our *hs* PCDH15 EC2-3 dimer and to the one generated by Dionne *et al.*^38^ Mismatched species (mouse *versus* human CDH23 EC1-2 differing at p.R35Q, p.P153Q, p.Q168R, and p.V174T) and engineered mutations are unlikely to explain this difference. Instead, electrostatic interactions among highly conserved charged residues from both CDH23 protomers favor a close contact in this region. Residue p.E49 from one CDH23 protomer “caps” the N-terminus of the neighboring CD α helix in the opposite protomer. This conspicuous long-range helix capping^72^ appears to stabilize the overall structure of the heterotetramer. Furthermore, p.E50 from one CDH23 protomer forms a salt bridge with p.R53 of the neighboring CDH23 and vice versa (Fig. S11C). Thus, the less open conformation of the PCDH15 “scissor” squeezes the CDH23 protomers together, creating a CDH23-CDH23 interface with favorable contacts among the two protomers, which in turn may stabilize the overall heterotetrameric structure. Taken together, our structures and data strongly support parallel dimerization of PCDH15 at the N-terminal end mediated by EC2-3 repeats and tip-link bonds forming a heterotetramer with two PCDH15 EC1-2 + CDH23 EC1-2 handshakes (Fig. 2 and Figs. S8-S11).

### Structures of PCDH15’s ectodomain middle region reveal atypical flexible linkers

The middle region of PCDH15, encompassing repeats EC4 to EC8, has been less thoroughly explored than other parts of this protein. A previous structure of *hs* PCDH15 EC3-5 CD1-1 (PDB: 5T4M) revealed two bound Ca^2+^ ions at the EC3-4 linker region and a canonical Ca^2+^-binding site at the EC4-5 linker region^44^. Protomers in the asymmetric unit displayed different orientations of EC3 with respect to EC4, and this flexibility was also evident in simulations of this fragment^44^. However, this *hs* PCDH15 EC3-5 CD1-1 structure lacked exon 12a, which encodes for a seven residue insertion p.V(414+1)PPSGVP(414+7) near the EC3-4 linker region. Given that structural variations due to isoform specific (in-frame) insertions or deletions might be relevant for function, we sought to obtain a structure that carries exon 12a, present in some of the CD2 isoforms thought to be essential for inner-ear mechanosensation^20, 73, 74^. We solved the *hs* PCDH15 EC3-5 CD2-1 structure, which contains three molecules in the asymmetric unit having distinct EC3-4 inter-repeat conformations and with two of the linkers with only one bound Ca^2+^ ion at site 3 (Fig. 3A-E). The insertion enlarges the EC4 BC loop, which projects away from the repeat without altering its folding but may affect Ca^2+^-binding affinity (Fig. 3B and Fig. S4C). Only the most bent conformation of this isoform is compatible with the antiparallel PCDH15 EC1-3 interface (Fig. S5), while all protomer conformations observed in the structure are compatible with the PCDH15 X-dimer mediated by EC2-3 (Fig. S12A). Overall, our *mm* PCDH15 EC3-5 CD2-1 structure is consistent with enhanced flexibility at the EC3-4 linker.

**Figure 3.**
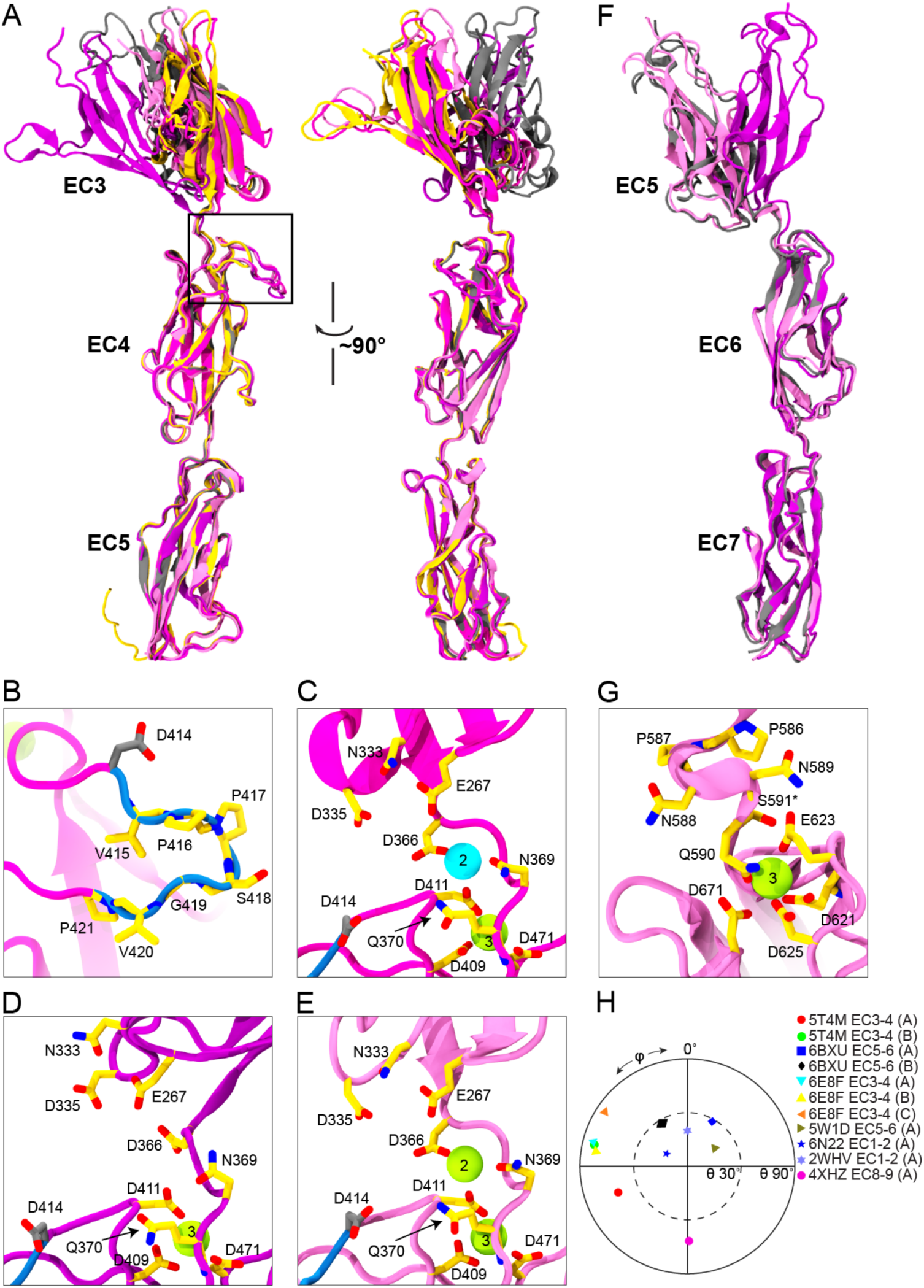
Flexibility of PCDH15 EC3-4 and EC5-6 linker regions. (**A**) Ribbon representation of *hs* PCDH15 EC3-5 CD2-1 with protomers (mauve, purple, and magenta) superposed in two views illustrating flexibility at EC3-4. Also superposed are protomers of *hs* PCDH15 EC3-5 CD1-1 (yellow and gray, PDB 5T4M^44^). Black box highlights location of CD2-1 insertion. (**B**) Detail of the insertion p.V(414+1)PPSGVP(414+7) encoded by exon 12a (blue backbone). Relevant residues are shown in stick representation and labeled. Some backbone atoms are omitted for clarity. Residue p.D414, thought to be under positive selection^134^ and involved in inherited deafness^135^ is shown in gray. (**C**-**E**) Detail of *hs* PCDH15 EC3-4 linker region for the three different protomers observed in the asymmetric unit of the *hs* PCDH15 EC3-5 CD2-1 structure. Shown as in B. Sodium is in cyan. (**F**) Ribbon representation of *mm* PCDH15 EC5-7 with protomers from *mm* PCDH15 EC4-7 (gray) and from *mm* PCDH15 EC5-7 I582T (mauve and purple) superposed. (**G**) Detail of atypical Ca^2+^-binding linker region in *mm* PCDH15 EC5-6 with only one Ca^2+^ at site 3. (**H**) Orientation of tandem EC repeats for listed PCDH15 linker regions (5T4M, 6BXU, 6E8F, 5W1D, 6N22, 4XHZ) and CDH23 EC1-2 (2WHV). The N-terminal EC repeat for labeled structures was used as reference and aligned to the *z* axis. The projection in the *x*–*y* plane of the principal axis of the C-terminal EC repeat was plotted. CDH23 EC1-2 was used to define the azimuthal angle φ as 0°.

Another atypical linker region is expected in PCDH15 EC5-6: Sequence alignments show that EC5 has a modified XEX motif (p.499TDM501) and lacks the DRE motif (p.559ALT561) present in canonical Ca^2+^-binding linker regions (Fig. S1). In addition, the DXNDN linker is p.586PPNNQ590 in EC5-6, but motifs involved in Ca^2+^ binding at site 3 (top of EC6) are mostly unchanged (DXD is p.621DRE623 and XDX is p.670SDG672). Two of our crystal structures cover the EC5-6 linker, which is seen in three different conformations (one in *mm* PCDH15 EC4-7 and two in *mm* PCDH15 EC5-7 I582T; Fig. 3F and Fig. S4D). In all cases, only one bound Ca^2+^ ion is observed at site 3 in EC6, as expected from the modified motifs listed above (Fig. 3G). Relative orientations of EC6 with respect to EC5 are significantly different (Fig. 3F-H and Fig. S12D). In contrast, structures of *mm* PCDH15 EC6-7 and *mm* PCDH15 EC7-8 V875A show canonical, rather straight linkers with three bound Ca^2+^ ions (variants I582T and V875A are not expected to alter structures). While the two PCDH15 EC5-7 protomers in the asymmetric unit form a compelling parallel *cis* dimer in the *mm* PCDH15 EC5-7 I582T crystal structure (interface area of 1049.5 Å^2^; Fig. S12B,C), all our fragments covering PCDH15 EC4-7, EC5-7, and EC6-7 were monomeric in solution, as also reported for similar fragments by Dionne et al^38^.

Overall, our structures of the middle region of PCDH15 suggest flexibility and altered Ca^2+^ binding at EC3-4 and EC5-6 linkers. Furthermore, three alternate conformations for each of the EC3-4 and EC5-6 linker regions assembled in an X-dimer mediated by EC2-3 give rise to 45 unique possible combinations of PCDH15 protomers (Fig. S12D). Some of these conformations display a remarkable separation between EC7 repeats (up to ∼26.8 nm), compatible with a PCDH15-PCDH15 *trans* X-dimer (Fig. S6A). Importantly, given prior simulations of the EC3-4 fragment^44^ and the architecture of EC5-6, flexibility at these linker regions will occur even at high Ca^2+^ concentrations, with a large number of structurally diverse configurations facilitated by the EC2-3 X-dimer.

### Structures of PCDH15’s ectodomain C-terminal fragments reveal kinks and *cis* dimerization

Three previous studies reported high-resolution structures covering parts of the C-terminal end of PCDH15 including EC9 to MAD12 (*hs* PCDH15 EC8-10, *mm* PCDH15 EC9-10, *ss* PCDH15 EC10-MAD12, and *mm* PCDH15 EC11-MAD12)^39, 40, 43^. These structures revealed a semi-canonical EC8-9 linker region, a flexible, Ca^2+^-free and bent EC9-10 linker region, a canonical EC10-11 linker region, and the L-shaped arrangement of a ferredoxin-like MAD12 tucked against EC11, with EC11-MAD12 inducing parallel dimerization^38–40^. Our structure of *mm* PCDH15 EC9-MAD12 (Fig. 4A,B) displays the bent EC9-10 linker region (as observed in *hs* PCDH15 EC8-10 and *mm* PCDH15 EC9-10, albeit with a different azimuthal angle)^43^ and the EC10-MAD12 dimerization domain, both in one symmetric dimer (one molecule per asymmetric unit). The total dimer interface surface area is 1,267.9 Å^2^ (including contacts between MAD12 and EC11 and between EC10 repeats; Fig. 4B), somewhat larger than what is observed in the more asymmetric *ss* PCDH15 E10-MAD12 structure^40^ (∼1,1160 Å^2^, two molecules per asymmetric unit), with the interface area at the EC10-EC10 contact being slightly larger (∼336 Å^2^ compared to ∼185 Å^2^ for 6BXZ). The *mm* PCDH15 EC9-MAD12 structure confirms that dimerization is induced by EC11-MAD12 and that it is compatible with a kinked EC9-10.

**Figure 4.**
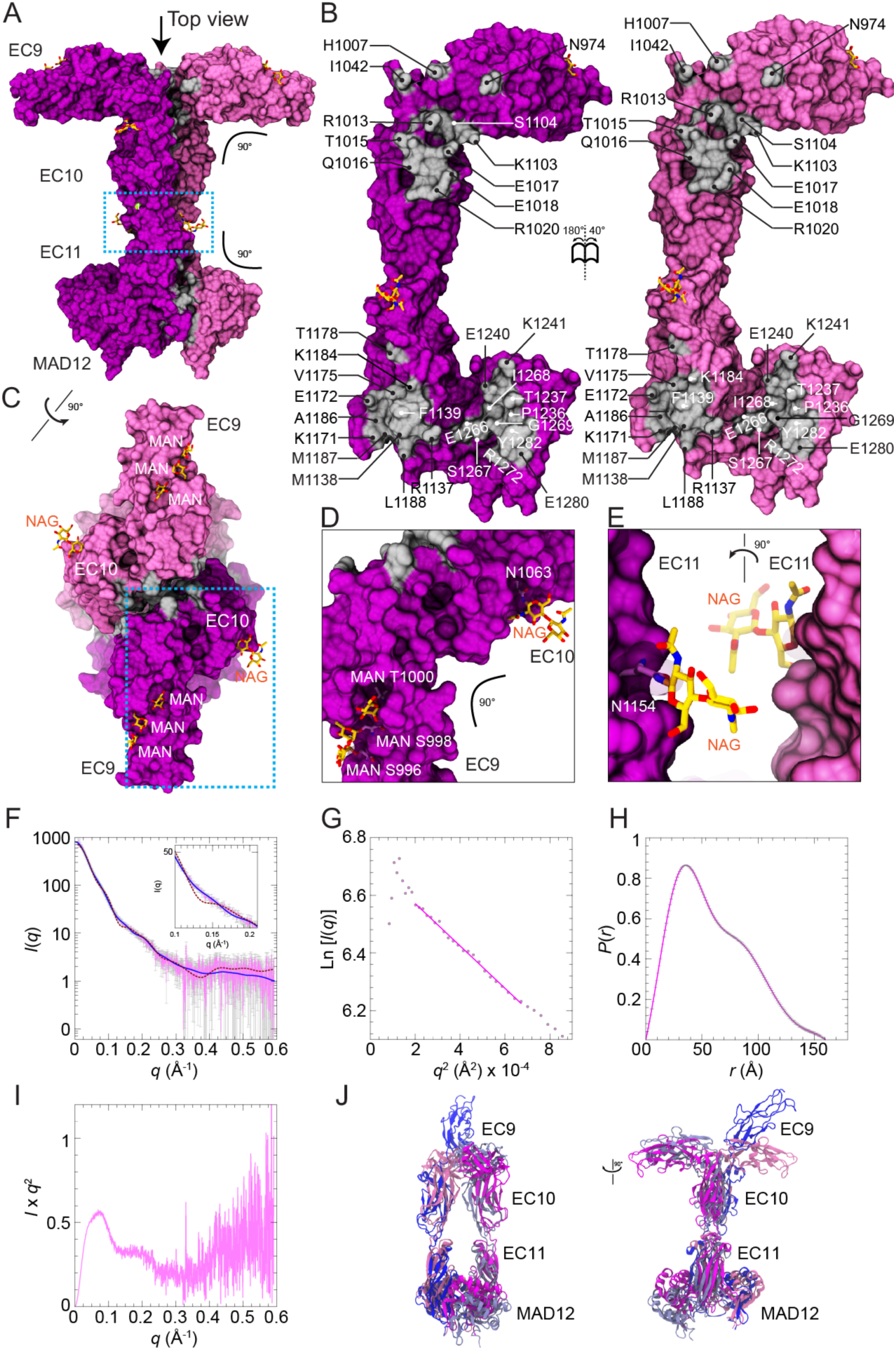
Dimerization of PCDH15 at the EC9-MAD12 region. (**A**) Molecular surface showing the glycosylated *mm* PCDH15 EC9-MAD12 parallel dimer in mauve and purple. Dimer interfaces are shown in gray, and glycosylation sugars are shown as yellow sticks. Blue dashed box denotes region highlighted in E. (**B**) Interaction surfaces exposed and colored as in A with interfacing residues labeled. (**C**) Top view of the *mm* PCDH15 EC9-MAD12 parallel dimer, with EC9 repeats pointing in opposite directions. Blue dashed box denotes region highlighted in D. (**D-E**) Details of glycosylation sites (dashed boxes in A and C). (**F**) X-ray scattering intensity as a function of the scattering vector *q* (SAXS profile) for the bacterially produced *mm* PCDH15 EC9-MAD12. Predicted scattering intensities from the structure obtained with FoXS are shown in maroon-dashed lines (χ^2^ = 6.52), while the theoretical scattering curve obtained from flexible refinement with SREFLEX is shown in blue (χ^2^ = 1.38). (**G**) Guinier plot of the SAXS data in the low *q* region. The magenta solid line shows the linear fit from which the gradient of the slope (-*R*_g_^2^/3) was used to estimate *R*_g_. (**H**) Real-space pair distribution function *P*(*r*). (**I**) Kratky plot indicating the presence of folded and flexible protein. (**J**) Representative model of *mm* PCDH15 EC9-MAD12 from flexible refinement with SREFLEX (dark and light blue) superposed to the crystal structure of *mm* PCDH15 EC9-MAD12, magenta and mauve).

Glycosylation is observed in our *mm* PCDH15 EC9-MAD12 structure at five sites, with O-linked sugars clearly discernible at residues p.S996, p.S998, and p.T1000 in EC9, and N-linked sugars at p.N1063 in EC10 and at p.N1154 in EC11 (Fig. 4C-E and Fig. S13). Interestingly, glycosylation has also been observed in published structures at p.N1063 and p.N1154, but not at the other three sites^39^. Clear electron density at additional glycosylation sites p.N1043 and p.S1141 (as seen in PDBs: 6C10 and 6C13^39^) is not present in our structure. While glycosylation at p.S1141 would sterically hinder the dimeric interface in one of the published PCDH15 structures (PDB: 6C13), none of the other modeled sugars in all available structures that cover this region interfere with bending and dimerization directly.

Intriguingly, bending at the EC9-10 linker region in the context of the dimer positions EC9 repeats pointing in opposite directions, with the projections along their longest principal axis being parallel to a hypothetical membrane plane (Fig. S6D). Data from size exclusion chromatography coupled to multi-angle light scattering (SEC-MALS) confirmed that this protein fragment is dimeric in solution (Fig. S7B and Table 2), and the EC9-10 “kink” is also consistent with our previous structures^43^ and with low-resolution cryo-EM images of a PCDH15 fragment encompassing EC8 down to its transmembrane helix^39^. Small angle X-ray scattering (SAXS) data obtained using bacterially produced *mm* PCDH15 EC9-MAD12 are also consistent with a dimeric arrangement in solution, with molecular weight estimates of 98.12 kDa and a 5.7 % discrepancy with the sequence-derived molecular weight of 51.98 × 2 = 103.06 kDa (Fig. 4F-J, Table S5).

SAXS data can also provide information about the size and shape of a protein in solution. The radius of gyration (*R*_g_) of bacterially produced *mm* PCDH15 EC9-MAD12 from Guinier (*R*_g_ = 46.72 ± 2.94 Å) and SAXS profile analyses (*R*_g_ = 47.28 ± 0.24 Å with maximum dimension *D*_max_ = 160 Å) are in excellent agreement with each other and with the *R*_g_ obtained from the *mm* PCDH15 EC9-MAD12 structure (*R*_g_ = 42 Å). These results further suggest that the overall shape of the dimer observed in the crystallographic structure is maintained in solution. However, comparison of the SAXS data to X-ray intensities modeled from the *mm* PCDH15 EC9-MAD12 crystal structure revealed some discrepancies reflected by a large χ^2^ value obtained for the fitting (χ^2^ = 6.52) and by a clear dip in the *q*-region 0.1 – 0.25 Å^−1^ observed in the modeled intensities but absent in the experimental data (Fig. 4F). X-ray intensities calculated from models obtained using a normal-mode analysis (SREFLEX^75^) fit the experimental data better than the crystal structure (χ^2^ = 1.38) and lack the dip in the *q*-region 0.1 – 0.25 Å^−1^ (Fig. 4F). The best model shows changes in orientation and rotation of EC9 along with loss of symmetry (Fig. 4J), consistent with asymmetry observed in SAXS-derived models of *ss* PCDH15 EC10-MAD12^40^. Analysis of the Kratky plot (Fig. 4I) indicates that while the protein is folded in solution, there is significant flexibility. Overall, SAXS data strongly supports a dimeric conformation of PCDH15 EC9-MAD12 that is asymmetric and flexible in solution.

Conformational transitions that straighten the EC9-10 linker, observed in simulations^43^ and in cryo-EM data^39^, strongly suggest that bending and unbending is relevant for tip-link assembly and function. The variety of angles that EC9-10 may adopt in solution^39^, along with dimerization of PCDH15 at EC2-3 and MAD12, and the asymmetry and flexibility suggested by SAXS data offer an interesting set of arrangements for PCDH15 + CDH23 filaments that might be adopted by tip-links and kinociliary links^14, 76, 77^ (Fig. S6E). In addition, flexing at EC3-4 (Fig. 3), EC5-6 (Fig. S12), and at EC9-10 (Fig. 4) seems to be a basic geometric requirement for establishing a parallel PCDH15 *cis* dimer compatible with dimerization points mediated by both EC2-3 and EC11-MAD12.

The structures and models discussed above depict parts of the entire ectodomain of PCDH15, and only small fractions of the entire tip link. Although each of them provides valuable insight into PCDH15 function, the assembly of PCDH15 filaments and force transduction mediated by tip links, which must withstand forces ranging from 10 to 100 pN^47, 48, 78, 79^, requires an understanding of the structure, dynamics, and elastic response of the entire PCDH15-CDH23 complex. Therefore, we used our structures to build, simulate, and computationally stretch monomeric PCDH15 EC1-MAD12 models, heterotetrameric PCDH15 EC1-5 + CDH23 EC1-2 or PCDH15 EC1-5 + CDH23 EC1-3 bonds, and two independent complete dimeric PCDH15 EC1-MAD12 models bound to CDH23 EC1-2 or CDH23 EC1-3.

### Structural models and predicted elasticity of the complete PCDH15 ectodomain monomer

To gain insights into the function and elastic behavior of PCDH15, and as similarly done for parts of CDH23^45^ and the entire ectodomain of the olfactory cell adhesion molecule^80^, we used overlapping structures of PCDH15 fragments and built two atomistic models of its entire ectodomain. The first model was built to get the *hs* PCDH15 EC1-MAD12 CD1-1 WT protein ectodomain (p.Q1 to p.I1342), which was coupled to a model of *hs* CDH23 EC1-2 (p.Q1 to p.D105) using the original handshake interaction^33^ as a template (PDB: 4APX; Fig. 5A, Fig. S14, and Table S6; ∼172 kDa for the complex). The second model was built to get the *mm* PCDH15 EC1-MAD12 CD2-1 WT protein ectodomain (p.D5 to p.S1341(+7) carrying the p.I582(+7)T and p.V875(+7)A variations) coupled to a model of *mm* CDH23 EC1-3 (p.V2 to p.D317) using the handshake interaction observed in the *hs* PCDH15 EC1-3 G16D/N369D/Q370N + *mm* CDH23 EC1-2 T15E structure (Fig. 5B, Fig. S15, and Table S6; ∼184 kDa for the complex). These PCDH15 models allowed us to visualize three in-frame deletion segments and fifteen sites of missense mutations implicated in inherited deafness (Fig. 5A-J and Table S7), thus providing a structural framework to propose mechanisms underlying dysfunction of PCDH15 (see Supplementary Discussion).

**Figure 5.**
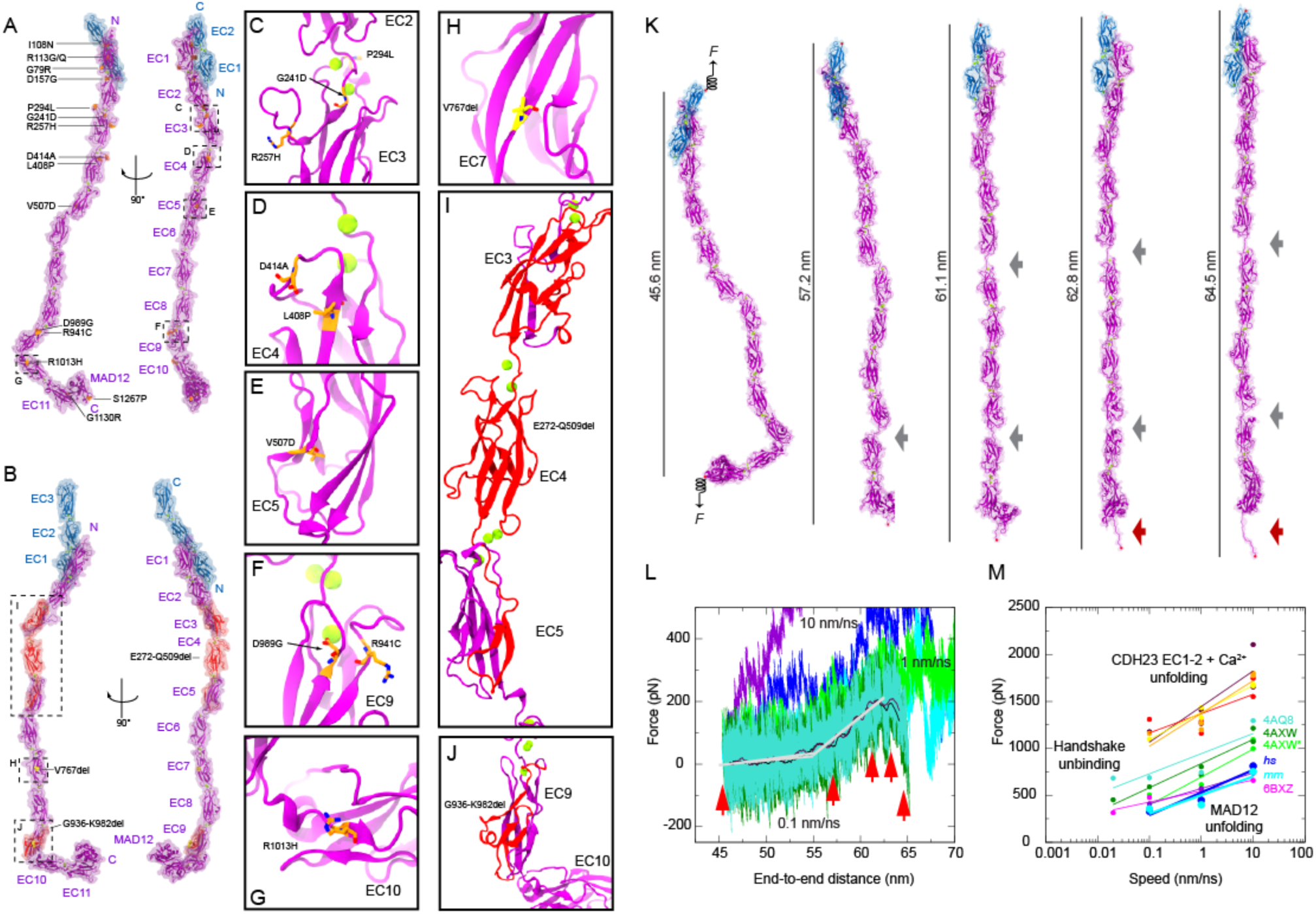
Disease-causing mutations and complete model and elasticity of the monomeric PCDH15 ectodomain. (**A**) Two views of the *hs* PCDH15 EC1-MAD12 CD1-1 + CDH23 EC1-2 model with PCDH15 deafness-causing missense mutations listed. Proteins are shown in purple (PCDH15) and blue (CDH23) ribbon representations with transparent molecular surfaces. Ca^2+^ ions are shown as green spheres and mutations are highlighted in orange. Dashed boxes and labels correspond to select missense mutations shown in C-G panels. (**B**) The *mm* PCDH15 EC1-MAD12 CD2-1 + CDH23 EC1-3 model with PCDH15 in-frame deletion mutations that cause deafness highlighted in red (multi-residue deletions) and yellow (single residue deletions). Dashed boxes and labels correspond to detailed views in H-J panels. (**C**-**J**) Detail of missense mutations and in-frame deletions that cause deafness. (**K**) Snapshots of the monomeric *hs* PCDH15 EC1-MAD12 CD1-1 + CDH23 EC1-2 system during stretching simulation S1d (0.1 nm/ns, Table S8). Stretched C-terminal Cα atoms are shown as red spheres. Springs in first panel indicate position and direction of applied forces. Gray arrows highlight stretching of PCDH15 EC linkers (EC9-10 followed by EC5-6). Dark red arrow indicates unfolding of PCDH15 MAD12’s C-terminal end. (**L**) Force versus end-to-end distance for constant velocity stretching of *hs* PCDH15 EC1-MAD12 + CDH23 EC1-2 at 10 nm/ns (S1b, purple and blue), 1 nm/ns (S1c, bright green and cyan), and 0.1 nm/ns (S1d, dark green and turquoise; 10-ns running averages shown in black and maroon; gray lines are fits used to determine elasticity of the complex). Red arrowheads indicate time-points for S1d illustrated in K. (**M**) *In silico* force peak maxima versus stretching speed for CDH23 EC1-2 unfolding (red, maroon, yellow, and orange)^42^, for CDH23 EC1-2 and PCDH15 EC1-2 handshake unbinding (turquiose-4AQ8, dark green-4AXW, and light green-4AXW* [after a 1 μs-long equilibration])^33, 37^, for PCDH15 EC10-MAD12 unfolding (magenta-6BXZ with unfolding at MAD12)^40^, for *hs* PCDH15 EC1-MAD12 CD1-1 + CDH23 EC1-2 unfolding (S1b-d, blue with unfolding at MAD12), and for *mm* PCDH15 EC1-MAD12 CD2-1 + CDH23 EC1-3 unfolding (S2b-d, cyan, with unfolding at MAD12).

Our models of the PCDH15 ectodomain adopt conformations dictated by the relative orientation (tilt and azimuthal angles) of successive EC repeats observed in various crystallographic structures, including the EC9-10 kink observed in *hs* PCDH15 EC9-MAD12^43^ (used for *hs* PCDH15 EC1-MAD12 CD1-1) and in *mm* PCDH15 EC9-MAD12 (used for *mm* PCDH15 EC1-MAD12 CD2-1). These may represent only some of the possible conformations adopted in solution and might not be representative of states under physiological tension as experienced by hair-cell tip links^78, 79^. Short equilibrations followed by SMD simulations of the entire *hs* and *mm* PCDH15 ectodomain monomers bound to the tips of CDH23 and with all their Ca^2+^-binding sites occupied, allowed us to explore their conformational dynamics and elasticity *in silico*. Constant-velocity stretching of the *hs* PCDH15 EC1-MAD12 CD1-1 + CDH23 EC1-2 heterodimer complex at 0.1 nm/ns revealed straightening of the structure (Fig. 5K, Fig. S16A, Table 3, and Table S8) in a first phase characterized by a soft effective spring constant (*k*_1_ ∼3.3 mN/m; Fig. 5L), followed by stretching of the EC5-6 and EC9-10 linker regions, ensued by stretching of the entire chain (*k*_2_ ∼24.4 mN/m). Subsequent unrolling and then unfolding of MAD12 at a force peak of ∼331 pN happened without unbinding of CDH23 EC1-2 from PCDH15. The elastic response of *mm* PCDH15 EC1-MAD12 CD2-1 + CDH23 EC1-3 at 0.1 nm/ns was very similar to that of the human complex (*k*_1_ ∼1.5 mN/m, *k*_2_ ∼27.5 mN/m, unfolding peak at ∼356 pN; Fig. S16B-C), suggesting that, at the stretching speeds used in our simulations, unfolding of MAD12 occurs before unbinding.

**Table 3.** Overview of MD simulations.

| System | Label | $t_{\text{sim}}$<br>(ns) | Type | Slowest speed<br>(nm ns<sup>-1</sup>) | Size<br>(#atoms) | Size<br>(nm<sup>3</sup>) |
| --- | --- | --- | --- | --- | --- | --- |
| *hs* PCDH15 EC1-MAD12 +<br>CDH23 EC1-2 + 3 Ca<sup>2+</sup>/linker | S1a-d | 215.4 | EQ/SMD | 0.1 | 924,024 | 71.2 × 9.2 × 14.9 |
| *mm* PCDH15 EC1-MAD12 +<br>CDH23 EC1-3 + 3 Ca<sup>2+</sup>/linker | S2a-d | 296.2 | EQ/SMD | 0.1 | 1,016,888 | 91.7 × 11.1 × 10.5 |
| *hs* (PCDH15 EC1-5)<sub>2</sub> +<br>(CDH23 EC1-2)<sub>2</sub> + 3 Ca<sup>2+</sup>/linker | S3a-d | 121.1 | EQ/SMD | 0.1 | 833,064 | 40.5 × 17.3 × 12.4 |
| *hs* (PCDH15 EC1-5)<sub>2</sub> +<br>(CDH23 EC1-2)<sub>2</sub> + 2 Ca<sup>2+</sup>/linker | S4a-d | 125.1 | EQ/SMD | 0.1 | 833,037 | 40.5 × 17.3 × 12.4 |
| *hs* (PCDH15 EC1-5)<sub>2</sub> +<br>(CDH23 EC1-2)<sub>2</sub> + 1 Ca<sup>2+</sup>/linker | S5a-d | 157.2 | EQ/SMD | 0.1 | 832,984 | 40.5 × 17.3 × 12.4 |
| *hs* (PCDH15 EC1-5)<sub>2</sub> +<br>(CDH23 EC1-2)<sub>2</sub> + 0 Ca<sup>2+</sup>/linker | S6a-d | 169.3 | EQ/SMD | 0.1 | 832,924 | 40.5 × 17.3 × 12.4 |
| *mm* (PCDH15 EC1-5)<sub>2</sub> +<br>(CDH23 EC1-3)<sub>2</sub> + 3 Ca<sup>2+</sup>/linker | S7a-d | 114.1 | EQ/SMD | 0.1 | 743,064 | 51.1 × 11.7 × 12.9 |
| *hs* (PCDH15 EC1-MAD12)<sub>2</sub> +<br>(CDH23 EC1-2)<sub>2</sub> + 3 Ca<sup>2+</sup>/linker | S8a-d | 153.2 | EQ/SMD | 0.1 | 1,178,985 | 99.4 × 11.4 × 10.9 |
| *mm* (PCDH15 EC1-MAD12)<sub>2</sub> +<br>(CDH23 EC1-3)<sub>2</sub> + 3 Ca<sup>2+</sup>/linker | S9a-d | 170.6 | EQ/SMD | 0.1 | 1,292,017 | 91.5 × 11.8 × 12.5 |
| *mm* (PCDH15 EC1-MAD12)<sub>2</sub> +<br>(CDH23 EC1-3)<sub>2</sub> + 3 Ca<sup>2+</sup>/linker | S10a | 95.3 | EQ | - | 2,325,962 | 65.5 × 18.9 × 19.5 |
| *mm* (PCDH15 EC9-MAD12)<sub>2</sub> + 3<br>Ca<sup>2+</sup>/linker | S11a-d | 124.8 | EQ/SMD | 0.1 | 456,282 | 28.2 × 13.2 × 13.2 |

Overview of MD simulations. System indicates proteins used. Type denotes the simulation protocol used (equilibrium: EQ, constant-velocity: SMD). Initial size of the systems is indicated in the last column.

The soft elasticity phase observed during stretching simulations of the monomeric PCDH15 EC1-MAD12 CD1/2-1 + CDH23 EC1-2/3 complexes stems mainly from unbending of the Ca^2+^-free EC9-10 linker region, as previously predicted^43^, and from stretching of EC5-6 and various other linkers. The predicted spring constants at 0.1 nm/ns are an upper limit, as this stretching speed is one order of magnitude larger than physiological speeds of the basilar membrane^81^ at loud, high-frequency sound and drag might be reduced at slower speeds. Similarly, unfolding forces of MAD12 at 0.1 nm/ns represent an upper bound as simulations and experiments at slower stretching speeds (loading rates) will naturally report smaller forces^82, 83^. The elastic response of MAD12 in the context of stretching simulations of the monomeric *hs* and *mm* PCDH15 EC1-MAD12 CD1/2-1 + CDH23 EC1-2/3 complexes is consistent with simulations stretching the dimeric *mm* PCDH15 EC9-MAD12 (Fig. S17) and our previous simulations of *ss* PCDH15 EC10-MAD12^40^. Our new simulations including the entire PCDH15 ectodomain predict that MAD12 will unfold before unbinding of the cadherin handshake at fast stretching speeds.

### Structural models and predicted strength of the heterotetrameric PCDH15 – CDH23 bond

To better understand the behavior of the handshake bond in the context of the heterotetrameric arrangement predicted from our *hs* PCDH15 EC2-3 structure and observed in the *hs* PCDH15 EC1-3 G16D/N369D/Q370N + *mm* CDH23 EC1-2 T15E complex, we built two model systems for simulation (Fig. 6A and Fig. S18A,C). The first one includes two molecules of *hs* PCDH15 EC1-5 CD1-1 (p.Q1 to p.P587) forming an X-dimer (as seen in the *hs* PCDH15 EC2-3 structure) coupled to two molecules of *hs* CDH23 EC1-2 (p.Q1 to p.D205; each arranged as seen in the *mm* PCDH15 EC1-2 + CDH23 EC1-2 complex [4APX]^33^ and mutated to match the human sequence). The second system includes two molecules of *mm* PCDH15 EC1-5 CD2-1 (p.Q1 to p.L585(+7)) forming an X-dimer coupled to two molecules of *mm* CDH23 EC1-3 (p.Q1 to p.D317) with the *cis* and *trans* interactions based on the *hs* PCDH15 EC1-3 G16D/N369D/Q370N + *mm* CDH23 EC1-2 T15E tetrameric structure. In both models, referred to here as *hs* and *mm* (PCDH15 EC1-5)_2_ + (CDH23 EC1-2/3)_2_, all linkers and repeats were in conformations obtained from crystal structures and all Ca^2+^-binding sites were occupied. To mimic possible low Ca^2+^ concentration conditions in the cochlea, three additional models of the *hs* (PCDH15 EC1-5)_2_ + (CDH23 EC1-2)_2_ tetramer were built with two, one, or zero Ca^2+^ ions per linker, respectively (Table 3, Supplementary Discussion, and Table S8). All systems, saturated and not saturated with Ca^2+^, were equilibrated and then stretched at various speeds to test their stability and elasticity *in silico*.

**Figure 6.**
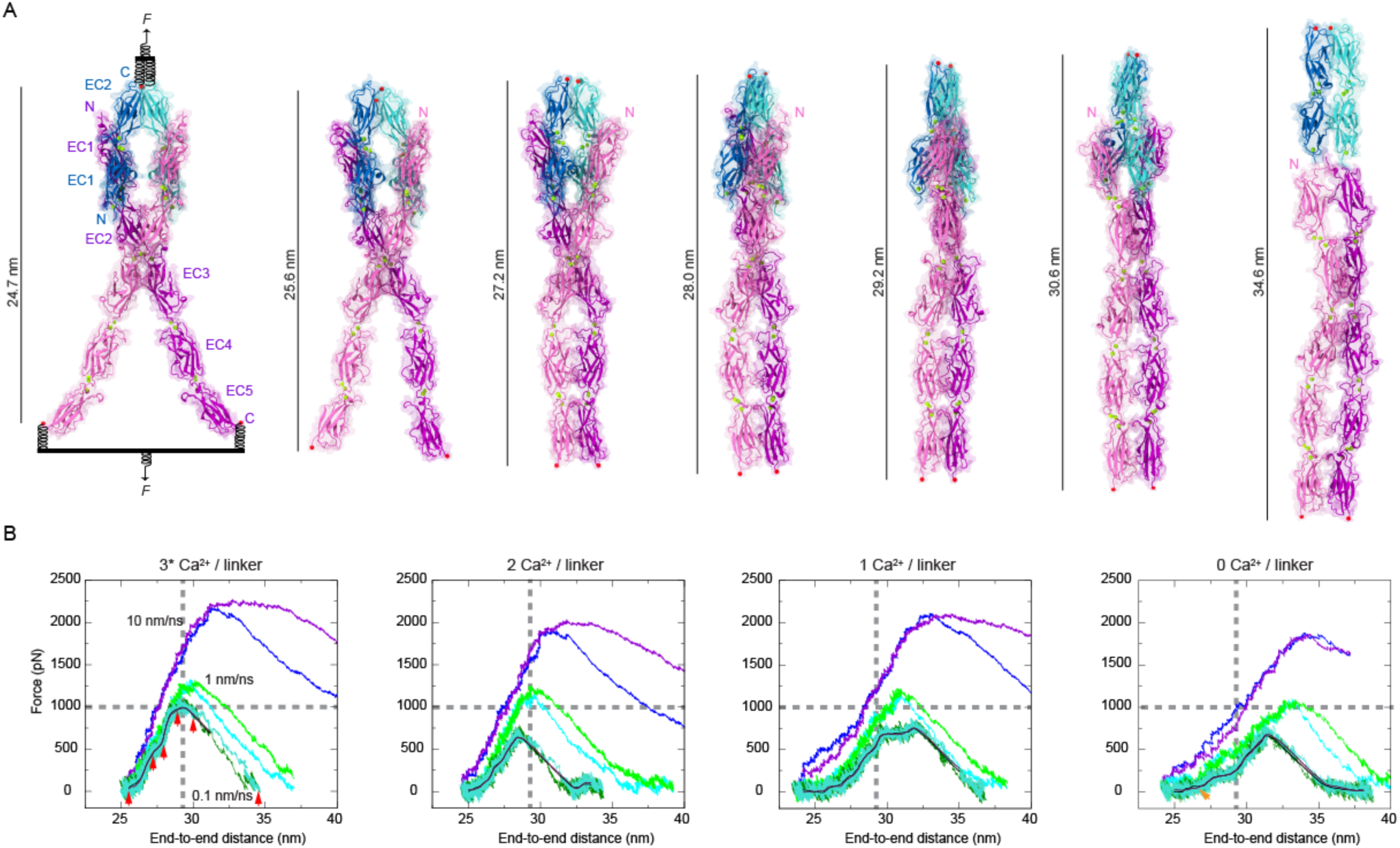
*In silico* strength of *trans* heterotetrameric bonds. (**A**) Snapshots of the heterotetrameric *hs* (PCDH15 EC1-EC5)_2_ + (CDH23 EC1-2)_2_ system during stretching simulation S3d (0.1 nm/ns, Table S8). Stretched C-terminal Cα atoms are shown as red spheres. Stretching was carried out by attaching two slabs to springs that were in turn attached to the terminal ends of each protein protomer. Slabs were moved in opposite directions at constant speed through individual springs. (**B**) Force applied to each of the slabs versus protein separation (see Methods) for constant velocity stretching simulations of *hs* (PCDH15 EC1-EC5)_2_ + (CDH23 EC1-2)_2_ systems with all Ca^2+^-binding sites occupied (S3b-d; 3 Ca^2+^ ions per linker except for PCDH15 EC2-3 and EC3-4 linkers), with 2 Ca^2+^ ions per linker (S4b-d; Ca^2+^-binding sites 2 and 3), with 1 Ca^2+^ ion per linker (S5b-d; Ca^2+^-binding site 3), and without bound Ca^2+^ ions (S6b-d). Traces are for constant velocity simulations at 10 nm/ns (purple and blue), 1 nm/ns (bright green and cyan), and 0.1 nm/ns (dark green and turquoise with 10-ns running averages in black and maroon). Gray dashed lines indicate magnitude and position of force peak at the slowest stretching speed tested with all Ca^2+^-binding sites occupied. Red arrowheads in leftmost panel indicate time-points for simulation S3d illustrated in A. Orange arrowhead in rightmost panel indicates soft elastic response region.

Equilibration trajectories of the *hs* and *mm* (PCDH15 EC1-5)_2_ + (CDH23 EC1-2/3)_2_ tetramers saturated with Ca^2+^ revealed a tendency for the PCDH15 EC5 ends to get closer to each other within 10 ns (from ∼16 nm in *hs* and ∼12 nm in *mm* to ∼10 nm in both). The *hs* CDH23 EC1-2 protomers also get close to each other, mimicking the arrangement seen in the *hs* PCDH15 EC1-3 G16D/N369D/Q370N + *mm* CDH23 EC1-2 T15E structure, where β-strands A and G of EC2 from one protomer get closer to the same strands in EC2 from the adjacent protomer. A similar arrangement was also seen in the equilibration of the *mm* complex model. However, the p.E50:p.R53 salt bridge between adjacent CDH23 protomers (as seen in the crystal structure of the heterotetramer bond) did not form in the *hs* complex, and *mm* CDH23 EC3 protomers drifted away from each other. Moreover, the PCDH15 p.R113 salt bridge with CDH23 p.E77, absent in the *mm* model, was formed throughout its equilibration trajectory. These rearrangements highlight the dynamic nature of the heterotetrameric complex. The double handshake bond, however, along with the PCDH15 EC2-3 interface for parallel dimerization, remained stable during these short equilibrations.

Systems with less Ca^2+^ at all linkers exhibited two distinct behaviors in our equilibrium simulations. The system with 2 Ca^2+^ ions per linker was stable, with the two PCDH15 EC5 protomers coming as close as ∼9 nm in a configuration that is more compatible with a parallel dimer than the one observed in the system with fully saturated linkers (3 Ca^2+^ ions per linker, except for PCDH15 EC2-3 and EC3-4). More dramatic conformational changes, including bending and twisting of linkers, were observed for systems with 1 and 0 Ca^2+^ ions per linker, especially at the PCDH15 EC3-4 linker regions. The conformational changes observed with less Ca^2+^, resulting in overall shrinkage of the end-to-end distance of the complex, are consistent with results from simulations of classical^84–86^ and tip-link cadherins^33, 42, 44^ that in the absence of bound Ca^2+^ ions exhibited enhanced hinge-like flexibility at their linker regions. These results are also consistent with EM images showing curled and collapsed classical^87^ and tip-link^21^ cadherins in the absence of Ca^2+^.

Stretching of the equilibrated *hs* and *mm* (PCDH15 EC1-5)_2_ + (CDH23 EC1-2/3)_2_ tetramers saturated with Ca^2+^ at the constant speeds tested (10, 1, and 0.1 nm/ns) revealed unbinding of the CDH23 EC1-2/3 protomers from PCDH15 without unfolding of EC repeats (Fig. 6A and Fig. S18A-C). At the slowest stretching speed (0.1 nm/ns), the *hs* (PCDH15 EC1-5)_2_ + (CDH23 EC1-2)_2_ complex straightened, with PCDH15 EC4-5 repeats establishing close contacts in a parallel arrangement. The “open scissor” conformation of the PCDH15 EC1-3 dimer closed itself as the stretching proceeded, thereby squeezing the CDH23 EC1-2 protomers together. As a result, an intricate network of hydrogen bonds developed between conserved CDH23 EC1 residues. Further, a rotation of the CDH23 EC1-2 protomers (Fig. 6 and Fig. S18A) preceded unbinding at large force (∼1080 pN, simulation S3d), which was roughly three times as large as the force observed during unfolding of MAD12 in the monomeric *hs* PCDH15 EC1-MAD12 + CDH23 EC1-2 complex (∼330 pN, simulation S1d), and ∼1.5 times larger than the force predicted for the unbinding of two non-interacting parallel handshakes (∼720 pN in Sotomayor *et al.*^33^). Similarly, stretching of the *mm* (PCDH15 EC1-5)_2_ + (CDH23 EC1-3)_2_ complex at 0.1 nm/ns resulted in unbinding at a large force peak (∼800 pN, simulation S7d), with squeezing of the CDH23 EC1-3 protomers (Fig. S19A-B). Thus, force-induced “locking” of CDH23 may make the tip-link bond more resistant against unbinding. The PCDH15 EC2-3 X-dimer interface was distorted but not lost after unbinding. These results suggest that the mechanical strength of the tetrameric tip-link bond is bolstered by the PCDH15 EC2-3 X-dimer.

Stretching of the equilibrated *hs* (PCDH15 EC1-5)_2_ + (CDH23 EC1-2)_2_ tetramers that were not saturated with Ca^2+^ at all linker regions revealed a more complex set of events during unbinding (Fig. 6B and Fig. S19A-B). SMD simulations of the system with 2 Ca^2+^ ions per linker at 0.1 nm/ns revealed unbinding without unfolding (force peak at ∼737 pN, simulation S4d) with complete separation of the PCDH15 EC2-3 X-dimer interface. SMD simulations of the system with 1 Ca^2+^ ion per linker region at 0.1 nm/ns revealed sequential unbinding (main force peak at ∼833 pN, simulation S5d) with severe stretching of linker regions. SMD simulations of the system with 0 Ca^2+^ ions per linker region at 0.1 nm/ns revealed minor partial unfolding, rupture of the EC2-3 X-dimer, and unbinding (force peak at 853 pN, simulation 6d). The starting states for SMD simulations of systems with 1 and 0 Ca^2+^ ions per linker region were shorter due to bending of ECs with respect to each other, resulting in a longer stretching phase (<25 nm to ∼27.5 nm in Fig. 6B and Fig. S19A) at low force with rather soft elasticity (*k*_1Ca_ ∼20.5 mN/m and *k*_0Ca_ ∼21.7 mN/m) dominated by unbending and straightening of the repeats. Extension of the linkers between EC repeats at higher force resulted in a displacement of the force peak to higher end-to-end distances. Interestingly, the elasticity stemming from the EC repeats also progressively softened with lower number of Ca^2+^ ions at the linker (Fig. S19A).

Overall, our simulations of the heterotetrameric PCDH15 – CDH23 bond suggest that parallel dimerization and Ca^2+^ ions modulate the strength of the tip link connection, which might in turn determine whether unbinding happens before unfolding.

### Structural models and predicted elasticity of the complete PCDH15 ectodomain dimer

Our structures and biochemical data obtained using various PCDH15 fragments indicate that there are two possible points of parallel dimerization (EC2-3 and EC11-MAD12, as also reported by Ge et al.^39^ and Dionne et al.^38^). Additional MALS experiments with the full-length ectodomain of *mm* PCDH15 EC1-MAD12 confirm that this fragment is a dimer that can withstand mild Ca^2+^-chelation with EDTA (Fig. S7C). However, atomistic models of the entire PCDH15 ectodomain seem to be incompatible with an arrangement in which both points of dimerization exist simultaneously (Fig. S6D), even when the PCDH15 EC9-10 linker unbends (Fig. S20A-E). This might reflect the constraints imposed by limited crystallographic conformations sampled by fragments containing only a few EC repeats, rather than the entire PCDH15 EC1-MAD12 protein. To model a parallel dimeric PCDH15 complex as observed in tip links^11, 21^, we did a systematic search of conformations obtained from equilibrium and SMD simulations that could be assembled into structures compatible with both points of dimerization (Fig. 7A,B and Fig. S20E-G).

**Figure 7.**
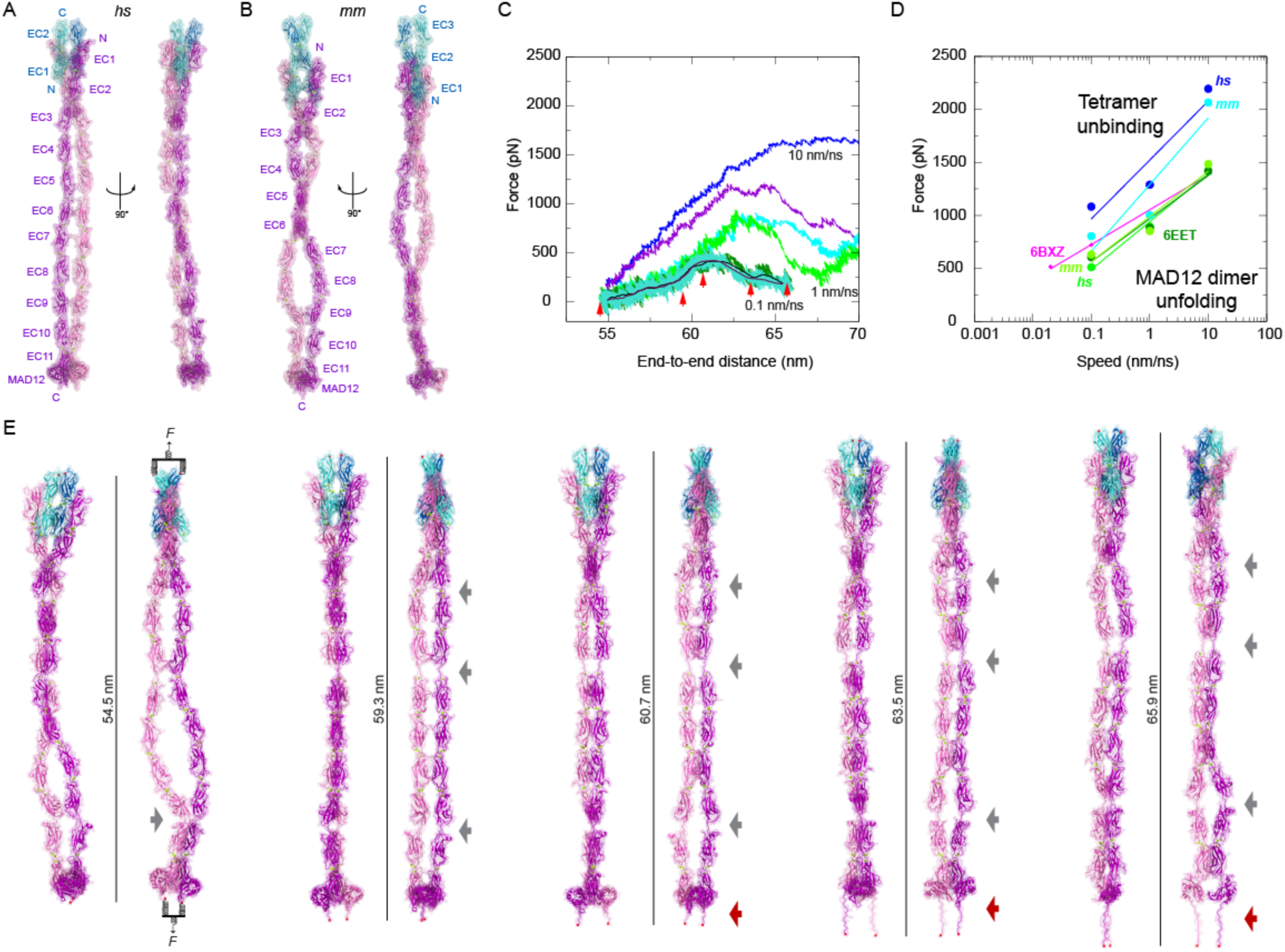
Models and mechanics of the PCDH15 ectodomain as part of an heterotetrameric complex with CDH23. (**A**-**B**) Two views of the (A) *hs* (PCDH15 EC1-MAD12)_2_ + (CDH23 EC1-2)_2_ and (B) *mm* (PCDH15 EC1-MAD12)_2_ + (CDH23 EC1-3)_2_ heterotetrameric systems before simulation. (**C**) Force applied to slabs versus protein separation (see Methods and Table S8) for constant velocity stretching simulations of the *hs* (PCDH15 EC1-MAD12)_2_ + (CDH23 EC1-2)_2_ system at 10 nm/ns (S8b, purple and blue), 1 nm/ns (S8c, bright green and cyan), and 0.1 nm/ns (S8d, dark green and turquoise with 10-ns running averages in black and maroon). Red arrowheads indicate time-points for simulation S8d illustrated in E. (**D**) *In silico* force peak maxima versus stretching speed for *hs* (PCDH15 EC1-EC5)_2_ + (CDH23 EC1-2)_2_ (S3b-d, blue), for PCDH15 EC10-MAD12 unfolding (magenta-6BXZ with unfolding at MAD12)^40^, for *mm* PCDH15 EC9-MAD12 dimer unfolding (S11b-d, dark green with unfolding at MAD12), and for *hs* (PCDH15 EC1-MAD12)_2_ + (CDH23 EC1-2)_2_ unfolding (S8b-d, bright green with unfolding at MAD12). (**E**) Snapshots of the heterotetrameric *hs* (PCDH15 EC1-MAD12)_2_ + (CDH23 EC1-2)_2_ system during stretching simulation S8d (0.1 nm/ns). Stretched C-terminal Cα atoms are shown as red spheres. Stretching was carried out by attaching two slabs to springs that were in turn attached to the terminal ends of each protein protomer. Slabs were moved in opposite directions at constant speed through individual springs. Gray arrows highlight stretching of PCDH15 EC linkers (EC9-10 followed by EC5-6) and re-bending at EC9-10. Dark red arrow indicates unfolding of PCDH15 MAD12’s C-terminal end.

The first tetramer model built contains the *hs* (PCDH15 EC1-MAD12 CD1-1)_2_ + (CDH23 EC1-2)_2_ complex (p.Q1 to p.I1342 and p.Q1 to p.D205 per monomer; ∼343 kDa), while the second contains the *mm* (PCDH15 EC1-MAD12 CD2-1)_2_ + (CDH23 EC1-3)_2_ complex (p.Q1 to p.S1341(+7) carrying the p.I582(+7)T and p.V875(+7)A variations for PCDH15 and p.Q1 to p.D317 for CDH23; ∼369 kDa; Table S6). All EC repeats and linker regions in these *hs* and *mm* (PCDH15 EC1-MAD12)_2_ + (CDH23 EC1-2/3)_2_ tetrameric models were obtained from crystallographic conformations or coordinates from simulations of crystal structures (Fig. S20F,G). In equilibrium simulations of the *hs* heterotetrameric model, in which the PCDH15 EC9-10 segment is initially straight (Fig. 7A and Fig. S20F), we monitored re-bending at this linker region, with one of the PCDH15 protomers more kinked than the other (Fig. 7E). A similar asymmetric bent conformation is maintained in an equilibration of the *mm* model (Fig. 7B and Fig. S21A). These results are consistent with simulations of the isolated PCDH15 EC8-10 fragment^43^ and suggest that the EC9-10 kink can exist within the context of the entire PCDH15 homodimer.

Equilibrated states of both *hs* and *mm* (PCDH15 EC1-MAD12)_2_ + (CDH23 EC1-2/3)_2_ tetrameric complexes were further used for SMD simulations at various stretching speeds (Fig. 7C-E and Fig. S21). In the slowest simulations (0.1 nm/ns) of the *hs* complex, straightening by unbending of various linker regions (most notably the EC9-10 kink) was followed by elongation of the EC3-4, EC5-6, and EC9-10 linkers and subsequent partial unrolling and unfolding of both MAD12s at the C-terminal end. Unbinding of CDH23 from PCDH15 was not observed at the slowest stretching speeds used. The initial straightening phase had an effective spring constant of *k* ∼45 mN/m, while unrolling and unfolding occurred at ∼510 pN, with a re-coil reflected in re-bending at the EC9-10 linker region (Fig. 7E, rightmost panel). A similar series of events was observed for the *mm* complex when stretched at 0.1 nm/ns, with an initial straightening phase that had an effective spring constant of *k* ∼37 mN/m and that was followed by MAD12 unrolling and unfolding at ∼630 pN. Simulations thus predict that the complete PCDH15 ectodomain dimer is significantly stiffer than the monomeric PCDH15 ectodomain.

## DISCUSSION

Our structures, biochemical data, and simulations provide an integrated atomistic view of the entire PCDH15 ectodomain and its connection to the CDH23 tip. We found two points of parallel dimerization for PCDH15 (at EC2-3 and EC11-MAD12) and provide a detailed and unique view of the two *trans* PCDH15-CDH23 handshakes in the heterotetrameric bond facilitated by the PCDH15 EC2-3 X-dimer. Simulations suggest that the two handshakes in the heterotetramer are squeezed together by closing of the X-dimer scissor under tension that mimics physiological stimuli. This squeezing strengthens the heterotetrameric bond with unbinding forces predicted to be on average ∼1.3 times larger than the sum of unbinding forces for independent handshakes, reminiscent of a catch-bond behavior^88^. Our structures also revealed three points of Ca^2+^-independent flexibility for PCDH15 at linker regions EC3-4, EC5-6, and EC9-10. Simulations of the entire monomeric PCDH15 ectodomain bound to CDH23 EC1-2/3 revealed a soft elastic response dominated by these flexible points, while the heterotetrameric complex (dimeric PCDH15 bound to two CDH23 monomers) was stiffer yet still presented kinks at the PCDH15 EC9-10 linker region. Intriguingly, our simulations predict that PCDH15’s MAD12 unrolls and then would unfold before CDH23 unbinding at the stretching speeds tested. These results provide a structural framework to compare and interpret complementary experimental results and to understand the function of PCDH15 as a key component of the tip links that open inner-ear transduction channels.

Two recent studies^38, 39^ have provided important insights into PCDH15’s structure using alternate and complementary approaches to those used here. Medium resolution (∼11 Å) cryo-EM images of PCDH15 EC8-MAD12 and its transmembrane domain in complex with TMHS^39^ eloquently revealed a highly dynamic EC8-10 ectodomain with either one or both protomers displaying kinked EC9-10 conformations that are consistent with our EC9-MAD12 structure and simulations. In the images reported by Ge *et al.*^39^, about 20% of the recorded conformations showed a bent conformation at this linker, with the refined straight model (∼14% of conformations) suggesting that bending-unbending transitions may play a role in tip-link mechanics^43^. However, it was unclear whether kinks and spontaneous bending-unbending dynamics were compatible with the complete dimeric ectodomain, as repeats EC1-7 were missing. Elegant structural models from negative staining EM images of longer PCDH15 ectodomain fragments (EC1-EC11 and EC1-MAD12) revealed flexural flexibility^38^ and were nicely consistent with parallel dimerization at both the EC2-3 contacts (confirmed by a high resolution structure of PCDH15 EC1-3^38^) and the EC11-MAD12 segments. However, reconstructions and averages may have neglected significant bending and flexibility observed in raw images (Fig. S8 in^38^), and resolution of the envelope (∼20 Å) was not enough to visualize details of the middle and C-terminal regions of PCDH15. Our structures, models, and simulations, obtained independently from the two studies mentioned above, not only fill in the missing pieces by providing a high-resolution and detailed view of the entire dimeric PCDH15 ectodomain, but also test its elasticity and dynamics *in silico*. Altogether, these data strongly support a model in which PCDH15 has two points of *cis* dimerization (EC2-3 and EC11-MAD12) and flexible linker regions at EC3-4, EC5-6, and EC9-10 that, along with unrolling of MAD12, may contribute to the overall elasticity of PCDH15.

Exquisite single-molecule force spectroscopy experiments^89^ have also started to elucidate the elastic response of the PCDH15 monomer. In these experiments, the monomeric PCDH15 EC1-EC11 p.V250D protein fragment stretched in the presence of different Ca^2+^ concentrations (3 mM, 20 μM, and no Ca^2+^) behaved as a soft entropic spring, with an effective spring constant that varied from < 0.5 mN/m to ∼6 mN/m before rupture. Three types of unfolding events (among many) were robustly identified, with extensions of ∼4 nm (A_U_), ∼15 nm (B_U_), and ∼35 nm (C_U_). The last type of unfolding event is consistent with rupture of an entire EC repeat. Comparison of these results with predictions from our simulations is difficult, as these experiments did not include CDH23 or MAD12 and were performed at slow stretching speeds equivalent to 1 Hz, with an artificial constant force of 2-4 pN maintained between stretching cycles. However, our predictions of reduced unfolding force peaks at low Ca^2+^ concentrations for various cadherins^33, 42, 85, 90^, including PCDH15 as presented here, are consistent with an increase in the number of PCDH15 unfolding events as observed in experiments upon Ca^2+^ depletion^89^. In addition, the predicted effective spring constants for the PCDH15 EC9-10 fragment at 0.02 nm/ns, *k* ∼8.4 mN/m, and for the heterodimeric PCDH15 EC1-MAD12 + CDH23 EC1-2 complex at 0.1 nm/ns, *k* ∼3 mN/m, are within the range of those measured experimentally^89^. Our simulations suggest that this soft elasticity stems from unbending of kinked linker regions (EC9-10) and elongation of flexible linkers (EC3-4 and EC5-6) in the presence of saturating Ca^2+^ concentrations, rather than from each EC repeat behaving as an individual part of a freely jointed chain^89^. Interestingly, lengthening of the heterodimeric PCDH15 EC1-MAD12 + CDH23 EC1-2 complex (∼15 nm, Fig. 5 and Fig. S16) matches lengthening in B_U_ events, but it is unclear why these continuous transitions in simulations would appear as discrete length jumps in experiments. Importantly, the elastic response of the monomeric PCDH15 ectodomain will be influenced by unrolling and unfolding of MAD12^40^, not present in the experiments. Since PCDH15 most likely functions as a *cis* dimer in the tip link^21^, the elastic response of the entire dimeric PCDH15 ectodomain, including the mechanically weak MAD12, needs to be taken into account.

Our simulations of the *hs* and *mm* (PCDH15 EC1-MAD12)_2_ + (CDH23 EC1-2/3)_2_ tetrameric complexes predict a stiffer elastic response (*k* ∼40 mN/m at 0.1 nm/ns with saturating Ca^2+^ concentrations) than that monitored for the monomer. Yet this estimate represents an upper bound, given the fast stretching speeds used in simulations and the possibility that further equilibration of our models results in additional bending of EC9-10 and the other flexible linker regions of PCDH15 (EC3-4 and EC5-6). A completely relaxed state with bent linkers will be softer than a pre-stretched initial conformation. Thus, our data suggests that the degree of bending of various linker regions in the initial state of the dimeric PCDH15 molecules, controlled in part by the Ca^2+^ concentration around it, will determine whether it behaves as a soft or stiff spring. Our data also suggests that dimerization of PCDH15 as well as binding to CDH23 will have a significant impact on the conformational space available to the PCDH15 monomers and their response to force upon tetramer formation.

Whether tip links made of PCDH15 and CDH23 act as soft gating springs or stiff cables conveying tension *in vivo* has been an open question for decades^47, 91, 92^. SMD simulations of *mm* CDH23 EC1-2 saturated with Ca^2+^ ions bound at its four sites (three at the EC1-2 linker and one at the EC1 tip) revealed a stiff fragment and large unfolding force peaks that were Ca^2+^-dependent. In absence of structural information from any other tip-link fragment at the time, the tip link was suggested to be stiff under the assumption that all other EC repeats would behave similarly^42^, yet further structural and computational work revealed flexible^44^ and elastic elements^43^ within PCDH15, and some atypical linkers in CDH23^45^. These data suggested that a kinked PCDH15 EC9-10 could provide Ca^2+^-independent elasticity and at the same time withstand the tip-link resting tension^78^, thus supporting a model in which the lower end of the tip link formed by PCDH15 could be soft, while the upper part of the tip link formed by CDH23 could be stiff^43^. Our data partially supports this model: the structure of *mm* PCDH15 EC9-MAD12 reveals that the EC9-10 kink is still present in the context of a longer construct, there are further points of Ca^2+^-independent elasticity in PCDH15 (particularly stemming from EC5-6), and the monomeric PCDH15 EC1-MAD12 is soft (*k* ∼3 mN/m at 0.1 nm/ns). However, while PCDH15 EC9-10 is still kinked in the context of the PCDH15 *cis* dimer, our heterotetrameric models are significantly stiffer than the heterodimeric models (*k* ∼40 mN/m at 0.1 nm/ns). Unrolling and unfolding of MAD12 along with the Ca^2+^-dependent response of various linker regions add another layer of complexity to the elastic response of PCDH15. The elastic response of MAD12 might be modified by its interactions to TMHS^39^ and other components of the transduction apparatus that may help prevent unrolling and unfolding. At very low Ca^2+^ concentrations PCDH15 might behave as a set of independent, freely jointed repeats^85, 89^, with resting tension keeping these chains pre-stretched. Thus, depending on binding partners and the conditions in which PCDH15 is found, it might behave as an elastic or stiff element (Fig. 8 and Fig. S22).

**Figure 8.**
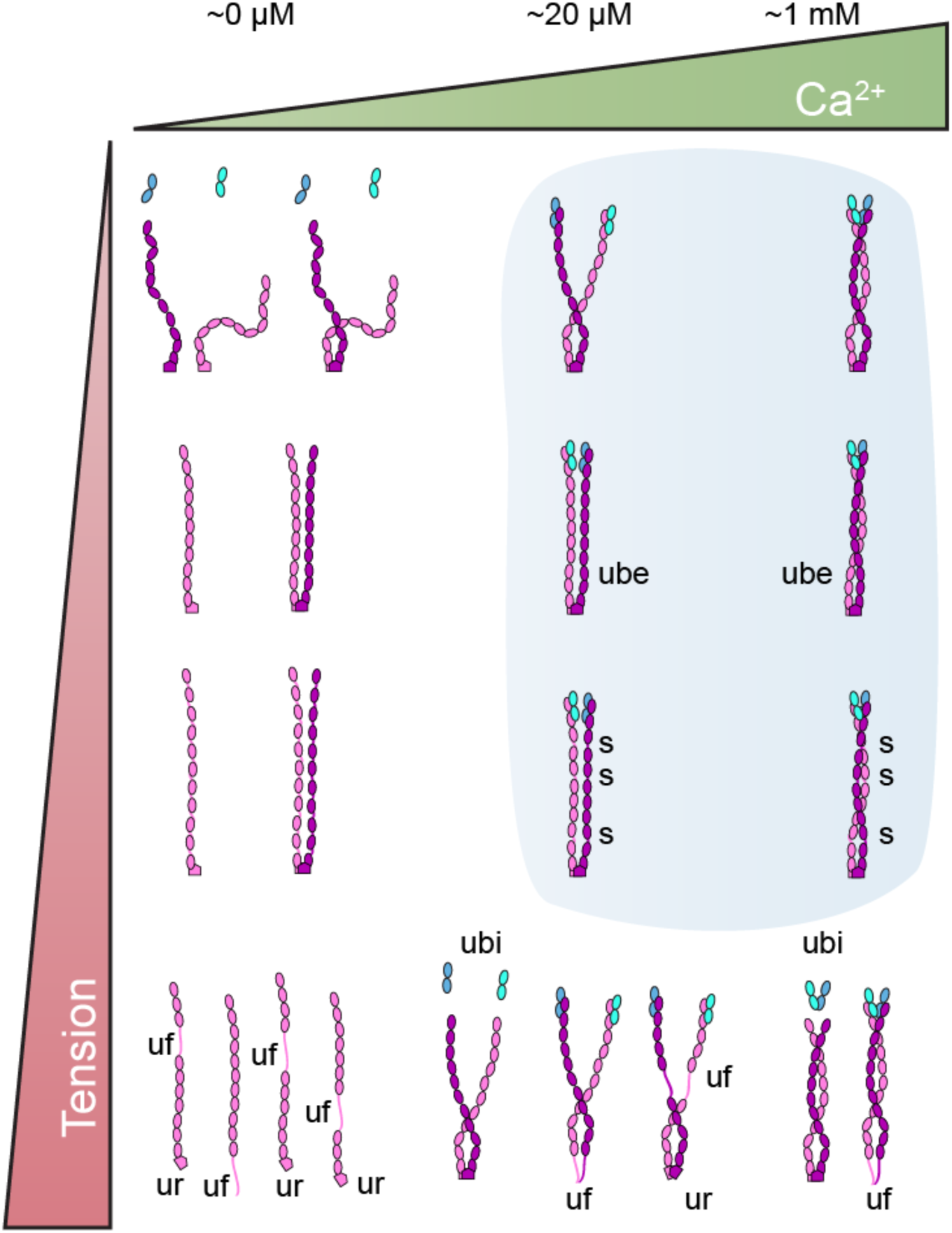
Tuning the elastic response of PCDH15. Structures and simulations indicate that the PCDH15 ectodomain can have distinct elastic responses that are tuned by tension and Ca^2+^. Diagram shows possible hypothetical states for PCDH15’s ectodomain. At very low Ca^2+^ concentrations (top left) PCDH15 will not interact with CDH23 and its linkers will be flexible and easily stretchable under tension. At high tension and very low Ca^2+^ concentrations (lower left), the Ca^2+^-free linkers will be weak and unfolding (**uf**) at any of the EC repeats will ensue. Under normal physiological conditions (blue shade), PCDH15 will form a *cis* dimer that interacts with CDH23 and that can exhibit soft elasticity mediated by bending and unbending transitions of Ca^2+^-free linker regions EC5-6 and EC9-10. As tension increases, unbending (**ube**) results in stiffening of PCDH15. Stretching of linkers (**s**) and unrolling of MAD12 (**ur**) at higher tension may soften the elastic response of PCDH15 for the subsequent stimulus. Under more extreme tension (bottom right) unbinding of PCDH15 from CDH23 (**ubi**) or unfolding of PCDH15 (**uf**) may occur. Glycosylation and specifics of PCDH15 splice isoforms may also influence the tip-link state and elastic response. The effect of resting tension is further dissected in Fig. S22.

Biophysical experiments with bullfrog saccular hair cells stimulated *ex vivo* with a flexible fiber first identified the gating spring, which was estimated to have a stiffness of *k* ∼0.5 mN/m in (1 mM or 4 mM Ca^2+^ saline)^47^. Similar experiments using an optical trap estimated the gating spring stiffness to be *k* ∼0.5 mN/m (4 mM Ca^2+^ saline)^48^, and a recent study using calibrated fluid-jet stimulation determined a gradient of gating spring stiffness along the mammalian cochlea ranging from ∼0.5 mN/m to ∼1.7 mN/m in inner hair cells and from ∼1.3 mN/m to 3.7 mN/m in outer hair cells (stiffness measurements done with 1.5 mM Ca^2+^ saline). Interestingly, the resting tension per tip link was also found to vary along the cochlea, from ∼5.9 pN to ∼16 pN for inner hair cells, and from ∼4.7 pN to ∼34 pN for outer hair cells. Maximal tip-link resting tension was estimated to be ∼50 pN. In all cases, the gating-spring stiffness and resting tension per tip link were indirectly estimated; we lack direct information on what forces each tip link is experiencing, whether resting tension is applied as a constant or fluctuating force, and whether force stimuli from sound is faithfully mimicked by constant loading-rate stretching of tip link molecules *in vivo*.

Laser velocimetry experiments from chinchilla cochlear hair cells *in vivo* indicate that the basilar membrane can move as fast as 0.01 m/s for loud sound (> 80 dB SPL)^81, 93^. Volumetric optical coherence tomography vibrometry data from mouse^94^ is consistent with the chinchilla measurements and indicate that peak stereocilia displacements in mice cochlear hair cells range between ∼10 nm to ∼150 nm in response to sounds that go from 10 dB SPL (∼6 kHz) to 80 dB SPL (∼10 kHz) *in vivo*. These data are in agreement with *in situ* estimates of mouse and guinea pig stereocilia displacements^95, 96^, and suggest that stereocilia can move as fast as 0.0015 m/s in normal hearing, and perhaps even faster for very loud sounds. Thus, stretching speeds used in our simulations are about one order of magnitude faster than those experienced by tip links *in vivo*, yet there is abundant theoretical^97–100^, computational^61, 82, 101^, and experimental work^83^ that can be used to interpret results at these high force loading rates. As stretching speed and loading rates increase, unbinding and unfolding forces are naturally larger, a phenomenon that is well documented in experiments^83^ and simulations^82^ and that is well understood theoretically^99, 100^. While we cannot rule out that the sequence and type of events (unfolding *vs*. unbinding) observed at fast stretching speeds in simulations will be the same at slower stretching speeds, our simulations were carried out at stretching speeds that are slower than those that may favor unfolding of terminal ends^102^ and provide specific predictions that can be tested and challenged experimentally *in vitro*.

*In vivo* conditions for tip links vary significantly across organs and species. For instance, Ca^2+^ concentration near tip links will greatly depend on the organ in which hair cells are located^103^. The vestibular endolymph Ca^2+^ concentration is > 100 μM, while the bulk cochlear endolymph Ca^2+^ concentration varies from 20 to 40 μM^52, 53^. Intriguingly, Ca^2+^ concentrations near hair bundles in the sub-tectorial space might be significantly higher than previously thought (> 300 μM), both because of the action of stereocilia Ca^2+^ pumps functioning in a restricted space and because of the buffering effect of the tectorial membrane^54, 55^. Similarly, resting tension and physiological forces will vary within and across organs, where glycosylation and differential expression of various tip-link PCDH15 isoforms may also be diverse. The complexity of PCDH15’s ectodomain revealed by our structure and simulations and the various set of environments in which it functions in mechanotransduction indicate that PCDH15 might be a versatile and multimodal protein that can be tuned to display distinct elastic responses (Fig. 8 and Fig. S22). While under some conditions PCDH15 can display the soft elasticity and extensibility attributed to gating springs, directly determining the exact *in vivo* conditions in which PCDH15 functions remains a necessary and challenging step required to fully comprehend the role played by tip links in inner-ear mechanotransduction.

PCDH15 is also found in auditory cortex interneurons^104^, eye photoreceptros^71, 105^, and various other tissues^23, 106^ where it may play a role in cell-cell adhesion and tissue development and maintenance. PCDH15’s interaction with CDH23 seems to be essential for auditory cortex wiring^104^, while its specific role in photoreceptor function is less clear. Our PCDH15 structures and biochemical assays probing mutations that impair heterophilic binding to CDH23, *cis* dimerization, and Ca^2+^ binding provide data that can inform the exploration of PCDH15’s function beyond inner-ear mechanotransduction.

## METHODS

### Cloning, expression, and purification of bacterially expressed PCDH15 fragments

DNA sequences encoding for protein fragments *mm* PCDH15 EC1-2, *hs* PCDH15 EC1-3, *hs* PCDH15 EC1-4, *mm* CDH23 EC1-2, *hs* PCDH15 EC2-3, *hs* PCDH15 EC3-5, *mm* PCDH15 EC7-8, and *mm* PCDH15 EC9-MAD12 were subcloned into NdeI and XhoI sites of the pET21a vector. Similarly, NheI and XhoI sites were used for fragments *mm* PCDH15 EC4-7, EC5-7, and EC6-7. All engineered missense mutations were generated using the QuikChange Lightning mutagenesis kit (Agilent). Insertions of a BAP sequence (p.GLNDIFEAQKIEWHE) at the end of *mm* PCDH15 EC1-2BAP and of exon 12a (p.VPPSGVP) in *hs* PCDH15 EC3-5 CD2-1 were carried out using standard protocols. All DNA constructs were sequence verified. Protein fragments were expressed in *Escherichia coli* BL21 (Agilent), BL21 CodonPlus(DE3)-RIPL (Agilent), or BL21 Rosetta(DE3) (Novagen) cells, which were cultured in LB or TB media, induced at OD_600_ ∼ 0.6 with 200 μM or 1 mM IPTG and grown at 30**°**C or 37**°**C for ∼16 hr (Table S3). Cells were lysed by sonication in denaturing buffer (20 mM TrisHCl [pH7.5], 6 M guanidine hydrochloride, 10 mM CaCl_2_, and 20 mM imidazole). The cleared lysates were loaded onto Ni-Sepharose (GE Healthcare), eluted with denaturing buffer supplemented with 500 mM imidazole and refolded as indicated in Table S3 using MWCO 2000 membranes (Spectra/Por). Refolded protein was concentrated (Vivaspin or Amicon 10 kDa) and further purified on Superdex S75 or S200 columns (GE Healthcare) in SEC buffer as indicated in Table S3. Pure fractions of *hs* PCDH15 EC1-3 G16D/N369D/Q370N and *mm* CDH23 EC1-2 T15E were mixed and purified in 20 mM HEPES pH 7.5, 150 mM KCl, and 2 mM CaCl_2_. All protein samples were concentrated by ultrafiltration to > 1 mg/ml for crystallization or biochemical assays.

### Cloning, expression, and purification of mammalian expressed PCDH15 fragments

Mouse PCDH15 EC1-MAD12 CD1-1 (p.Q1 to p.G1327), EC1-3 (p.Q1 to p.N369), EC1-4 (p.Q1 to p.N491), and EC9-MAD12 (p.M905 to p.E1360) were subcloned into a pHis-N1 vector (a modified version of the pEGFP-N1 vector from Clontech where the EGFP has been substituted for a hexahistidine tag) using XhoI and KpnI sites. The native signal sequence was included before the start of EC1 and EC9, mutation p.V250N was introduced in EC1-4 using the QuikChange Lightning mutagenesis kit (Agilent), and all constructs were sequence verified. All protein fragments were expressed by transient transfection of Expi293 cells using ExpiFectamine. After 4-5 days of expression, the conditioned media (CM) was collected and dialyzed overnight against 20 mM TrisHCl, pH 7.5, 150 mM KCl, 50 mM NaCl, and 10 mM CaCl_2_ to remove EDTA. The CM was concentrated using Amicon 10 kD or 30 kD concentrators and incubated with Ni-Sepharose beads for 1 h. The beads were washed 3 times with 20 mM TrisHCl, pH 8.0, 300 mM NaCl (200 mM NaCl for *mm* PCDH15 EC1-MAD12), 10 mM CaCl_2_, and 20 mM imidazole, and the target protein was eluted with the same buffer containing 500 mM imidazole. The *mm* PCDH15 EC9-MAD12 protein was further purified on a Superdex S200 S16/600 column in 20 mM TrisHCl, pH 7.5, 150 mM KCl, 50 mM NaCl, and 2 mM CaCl_2_ and concentrated to 8 mg/mL for crystallization. The *mm* PCDH15 EC1-3 and *mm* PCDH15 EC1-4 (WT and p.V250N) protein fragments were purified using the same procedure and concentrated for SEC-MALS analyses. Last, the *mm* PCDH15 EC1-MAD12 protein was further purified on a Superose 6 10/300 column in 20 mM TrisHCl, pH 8.0, 150 mM KCl, and 5 mM CaCl_2_ and concentrated for SEC-MALS analyses.

### Crystallization, data collection, and structure determination

Crystals were grown by vapor diffusion at 4**°**C by mixing protein and reservoir solutions as indicated (Table S2). Cryoprotection buffers were prepared as indicated in Table S2. All crystals were cryo-cooled in liquid N_2_. X-ray diffraction data was collected as indicated in Table 1 and processed with HKL2000^107^. Structures were determined by molecular replacement using PHASER^108^. The structure of *mm* PCDH15 EC1-2BAP was solved using repeats EC1-2 from *mm* PCDH15 in the handshake complex with *mm* CDH23 EC1-2 (4APX)^33^. The *mm* PCDH15 EC1-2 + CDH23 EC1-2 complex (4APX^33^) and the *hs* PCDH15 EC2-3 V250N fragment (see below) were used to solve the *hs* PCDH15 EC1-3 G16D/N369D/Q370N structure, which was used along with the *mm* PCDH15 EC1-2 + CDH23 EC1-2 complex (4AQ8)^33^ to solve the structure of *hs* PCDH15 EC1-3 G16D/N369D/Q370N in complex with *mm* CDH23 EC1-2 T15E. Refinement of this heterotetrameric structure used the amplitude-based “Twin Refinement” option in REFMAC5 after achieving an *R*_free_ value of ∼32%. The *hs* PCDH15 EC2-3 WT structure was solved using repeat EC2 from *mm* PCDH15 in the handshake complex with *mm* CDH23 EC1-2 (4APX) and EC3 from *hs* PCDH15 EC3-5 (5T4M), and subsequently used to solve for *hs* PCDH15 EC2-3 V250N. The *hs* PCDH15 EC3-5 CD2-1 structure was solved using *hs* PCDH15 EC3-5 (5T4M^44^). Refinement of this structure used the amplitude-based “Twin Refinement” option in REFMAC5 after achieving an *R*_free_ value of ∼27%. Initial search models for *mm* PCDH15 EC4-7 were individual EC4 and EC5 repeats from *hs* PCDH15 EC3-5 (5T4M^44^), and EC7 from *mm* PCDH15 EC7-8 V875A. The *mm* PCDH15 EC5-7 structure was solved using individual EC repeats from *mm* PCDH15 EC4-7. Mouse PCDH15 EC6-7 was solved using EC7 from *mm* PCDH15 EC7-8 V875A as an initial search. The *mm* PCDH15 EC9-MAD12 structure was solved using *ss* PCDH15 EC10-MAD12 (6BXZ)^40^ and *hs* PCDH15 EC8-10 (4XHZ)^43^. Model building was done with COOT^109^ and restrained refinement was performed with REFMAC5^110^ as indicated in the deposited structures. Data collection and refinement statistics are provided in Table 1. Structures were further analyzed using Procheck^111^, Whatcheck^112^, and Checkmymetal^113^ prior to deposition.

### SEC-MALS

SEC-MALS experiments were done using an ÄKTAmicro system connected in series with a Wyatt miniDAWN TREOS system. Protein samples of *hs* PCDH15 EC2-3 (> 1 mg/mL) were separated on a Superdex S75 3.2/30 column in 20 mM TrisHCl, pH 8.0, 150 mM KCl, 50 mM NaCl and 2 mM CaCl_2_. Bacterially produced protein samples of *hs* PCDH15 EC1-3 and EC1-4 (WT and p.L306N/V307N mutants), as well as mammalian expressed protein fragments *mm* PCDH15 EC1-3 and EC1-4 (WT and p.V250N), all at concentrations > 1 mg/mL, were separated on a Superdex S200 3.2/3.0 column in 20 mM TrisHCl, pH 8.0, 150 mM KCl, 50 mM NaCl and 2 mM CaCl_2_. Mammalian expressed *mm* PCDH15 EC1-MAD12 (∼1.3 mg/mL) was separated on a Superose 6 3.2/30 column in 20 mM TrisHCl, pH 8.0, 150 mM KCl, with either 5 mM CaCl_2_ or 5 mM EDTA. Absorbance at 280 nm and light scattering were monitored. The scattering information was subsequently converted into molecular weight using a rod-like model (Table 2). The SEC-MALS curves were adjusted for connecting tubing length before plotting. Measurements listed in Table 2 were taken from distinct samples.

### SEC-SAXS

SEC-SAXS data was collected at the SIBYLS beamline 12.3.1 in the Advanced Light Source facility (Berkeley, CA) as described^114, 115^ (Table S5). A bacterially produced and purified sample of *mm* PCDH15 EC9-MAD12 was used along with an Agilent 1260 series HPLC and a Shodex KW-803 analytical column for data collection at a flow rate of 0.5 ml/min at 20**°**C (20 mM TrisHCl pH 8.0, 150 mM KCl, 5 mM CaCl_2_, and 1 mM TCEP). X-ray exposures lasting 3 s were collected continuously during a ∼40 min elution. SAXS frames recorded prior to the protein elution peak were used to subtract all other frames. Buffer subtraction and data reduction was performed at the beamline with SCÅTTER^116^.

Further data analysis of the merged SAXS data was carried out with PRIMUS^117^ and the ATSAS program suite^118^. Estimates of the radius of gyration (*R*_g_) from the Guinier region were measured with PRIMUS. Maximum dimension (*D*_max_) of particles was estimated from an indirect Fourier transform of the SAXS profiles using GNOM^119^. Values of *D*_max_ between 140 and 160 Å provided the best solutions. The oligomeric state of the sample was assessed by estimating its molecular weight using the method implemented in the SAXSMoW2 server^120^. Search of conformational changes of the dimeric structure in solution was performed by normal modes analysis with SRFLEX^75^ using the crystal structure of *mm* PCDH15 EC9-MAD12 (PDB: 6EET) as starting model. Model scattering intensities were computed from *mm* PCDH15 EC9-MAD12 (PDB: 6EET) and fitted to the experimental SAXS data using FoXS^121^.

### AUC

Sedimentation velocity AUC experiments were performed in a ProteomeLab XL-I analytical ultracentrifuge (Beckman Coulter) following standard procedures^122–124^. Briefly, SEC purified protein samples were loaded into AUC cell assemblies with Epon centerpieces and 12 mm path length. To achieve chemical and thermal equilibrium, the An-50 TI rotor with loaded samples was allowed to equilibrate for ∼2 h at 20°C in the centrifuge. The rotor was spun at 50,000 rpm and data was collected using absorption optics. Data analysis was performed with the software SEDFIT (http://sedfitsedphat.nibib.nih.gov), using a continuous sedimentation coefficient distribution model *c*(*S*). Standard values for buffer viscosity (0.01002 poise), density (1 g/ml) and partial specific volume (0.73 ml/g) were used, and confidence level was set to 0.68 as routinely done. The obtained *c*(*S*) distribution was loaded in GUSSI^125^. All experiments were done in duplicates from distinct samples.

### Simulated systems

The psfgen, solvate, and autoionize VMD^126^ plugins were used to build eleven molecular systems for simulations (Table 3 and Tables S6 and S8). The *hs* PCDH15 EC1-MAD12 + CDH23 EC1-2 model (1,547 residues, ∼171.8 kDa) was built from fragments *mm* PCDH15 EC1-2 (4APX^33^), *hs* PCDH15 EC2-3 (chain D), *hs* PCDH15 EC3-5 (5T4M^44^, chain A), *mm* PCDH15 EC4-7, *mm* PCDH15 EC7-8 V875A, *hs* PCDH15 EC8-10 (4XHZ^43^), and *ss* PCDH15 EC10-MAD12 (6BXZ^40^, chain C) as indicated in Table S6 and Fig. S14. The *mm* PCDH15 EC1-MAD12 CD2-1 + CDH23 EC1-3 model (1,665 residues, ∼184.4 kDa) was built from fragments *hs* PCDH15 EC1-3 G16D/N369D/Q370N + *mm* CDH23 EC1-2 (chains B & C, respectively), *hs* PCDH15 EC3-5 CD2-1 (chain B), *mm* PCDH15 EC4-7, *mm* PCDH15 EC7-8 V875A, hs PCDH15 EC8-10 (4XHZ^43^), *mm* PCDH15 EC9-MAD12, and *dr* CDH23 EC1-3 (5W4T^45^, chain A) as indicated in Table S6 and Fig. S15. The *hs* (PCDH15 EC1-5)_2_ + (CDH23 EC1-2)_2_ model (1,587 residues, ∼176.6 kDa) was built from crystal structures of *mm* PCDH15 EC1-2 (4APX^33^), *hs* PCDH15 EC2-3 (chain D), and *hs* PCDH15 EC3-5 (5T4M^44^, chain A). The *mm* (PCDH15 EC1-5)_2_ + (CDH23 EC1-3)_2_ model (1,818 residues, ∼201.7 kDa) was assembled using structures *hs* PCDH15 EC1-3 G16D/N369D/Q370N + *mm* CDH23 EC1-2, *hs* PCDH15 EC3-5 CD2-1 (chain B), and *dr* CDH23 EC1-3 (5W4T^45^, chain A). All fragments were mutated so as to obtain a model of the species-specific WT proteins (Table S6). The *hs* (PCDH15 EC1-5)_2_ + *hs* (CDH23 EC1-2)_2_ systems with less Ca^2+^ ions at the linker regions were built from the model with Ca^2+^-saturated linkers by removing ions sequentially while maintaining charge neutrality.

The *hs* (PCDH15 EC1-MAD12)_2_ + (CDH23 EC1-2)_2_ tetramer system was assembled from the coordinates obtained from simulation trajectories as follows (Fig. S20F). Chain A of the *hs* PCDH15 used the coordinates from simulation S3c for residues 1-238 (chain A at 6.17 ns), whereas residues 239-485 come from simulation S3c (chain B at 6.17 ns), residues 486-1228 come from simulation S1d (118.342 ns), and residues 1229-1342 come from simulation S11c (chain A at 2.88 ns). Chain B of the *hs* PCDH15 used the coordinates from simulation S3c for residues 1-367 (chain B at 6.17 ns), whereas residues 368-901 come from simulation S1d (7.057 ns), residues 902-1003 come from simulation S1d (aa8.332 ns), residues 1004-1126 come from simulation S1d (117.069 ns), and residues 1127-1342 come from simulation S11c (chain B at 2.88 ns). The *hs* (CDH23 EC1-2)_2_ fragment comes from simulation S3c (chains C & D at 6.17 ns).

The *mm* (PCDH15 EC1-MAD12)_2_ + (CDH23 EC1-3)_2_ tetramer system was assembled from the coordinates obtained from simulation trajectories and crystal structures as follows (Fig. S20G). Residues 3-116, 125-164, and 182-199 of CDH23 EC1-3 come from the crystal structure of *hs* PCDH15 EC1-3 G16D/N369D/Q370N + *mm* CDH23 EC1-2 (chain C), residues 114-121(+3), 162-178(+3), and 197-315(+3) come from the crystal structure of *dr* CDH23 EC1-3 (5W4T^45^; chain A). Residues 1-359 of chain A in the *mm* PCDH15 EC1-MAD12 model come from the crystal structure of *hs* PCDH15 EC1-3 G16D/N369D/Q370N + *mm* CDH23 EC1-2 (chain A), whereas residues 360-493 come from simulation S7b (chain A at 0.19 ns). Residues 494-600 in chain A of PCDH15 come from the equilibration simulation S2a at 2.502 ns, whereas residues 601-913 come from the constant velocity SMD simulation S2c at 1.1 ns, and residues 914-1348 come from the constant velocity SMD simulation S11d at 61.694 ns. Residues 360-369 in chain B of PCDH15 come from the constant velocity SMD simulation S2b at 0.19 ns, whereas residues 370-495 come from the constant velocity SMD simulation S2b at 2.755 ns, residues 496-597 come from the equilibration simulation S2a at 4.182 ns, residues 598-800 come from constant velocity SMD simulation S2c at 8.1 ns, residues 801-911 come from constant velocity SMD simulation S2c at 9.24 ns, residues 912-1021 come from constant velocity SMD simulation S11d at 34.384 ns, and residues 1022-1348 come from constant velocity SMD simulation S11d at 61.694 ns. Coordinates for tetrameric complexes are available upon request.

All protein structures and models described above, along with crystallographic water molecules, had hydrogen atoms automatically added by psfgen. Residues D, E, K and R were assumed charged. Histidine residues were assumed neutral, and their protonation state was chosen to favor the formation of evident hydrogen bonds. Systems were solvated in explicit water, neutralized, and ionized with randomly placed ions so as to mimic a physiological endolymph environment with 150 mM KCl. The *mm* (PCDH15 EC1-MAD12)_2_ + *mm* (CDH23 EC1-3)_2_ tetramer was solvated in two different water boxes, a smaller system for SMD and a larger system for long free-dynamics equilibrations.

### MD simulations using NAMD

MD simulations were performed using NAMD 2.11, 2.12, and 2.13^127^, the CHARMM36 force field for proteins with the CMAP correction and the TIP3P model for water^128^. A cutoff of 12 Å (with a switching function starting at 10 Å) was used for van der Waals interactions. Periodic boundary conditions were used along with the Particle Mesh Ewald method to compute long-range electrostatic forces without cutoff and with a grid point density of >1 Å^−3^. A uniform 2 fs integration time step was used together with SHAKE. Langevin dynamics was utilized to enforce constant temperature *T* = 300 K with a damping coefficient of 0.1 ps^−1^, unless otherwise stated. Constant pressure simulations (*NpT*) at 1 atm were conducted using the hybrid Nosé-Hoover Langevin piston method with a 200 fs decay period and a 100 fs damping time constant.

### Simulations and analysis tools

Each system was energy-minimized and equilibrated in the *NpT* ensemble, and the resulting state was used to perform subsequent equilibrium and SMD simulations (Table 3 and Table S8). Constant-velocity stretching simulations used the SMD method and the NAMD Tcl forces interface^59–61, 129^, whereby Cα atoms of specific terminal residues were attached to independent virtual springs of stiffness *k_s_*= 1 kcal mol^−1^ Å^−2^. For SMD simulations of dimeric (simulations S11b-d) and heterotetrameric systems (simulations S3b-d, S4b-d, S5b-d, S6b-d, S7b-d, S8b-d, S9b-d), these springs were connected to virtual slabs attached to a third stretching spring (*k_s_* = 1 kcal mol^−1^ Å^−2^). The free ends of the stretching springs were moved away from the protein in opposite directions at a constant velocity. The stretching direction was set along the *x*-axis matching the vector connecting terminal regions of the proteins in the corresponding simulated complexes. Applied forces were computed using the extension of the virtual springs. Stiffness was computed through linear regression fits of force-distance plots. Maximum force peaks and their averages were computed from 50 ps running averages used to eliminate local fluctuations. The end-to-end distance in constant-velocity SMD simulations of heterotetrameric systems (S3 to S10 in Table 3 and Table S8) was computed as the separation between the center-of-mass of the Cα atoms stretched on one end and the center-of-mass of the Cα atoms stretched in the opposite direction on the other end of the system. Principal axes of EC repeats were computed using the Orient VMD plugin. Sequence alignments were performed with MUSCLE. Plots and curve fits were prepared with Xmgrace. Molecular images were created with the molecular graphics program VMD^126^. PCDH15 ectodomain sequences were compared among various species using data obtained from the NCBI protein database (Table S1). Ectodomain sequences equivalent to *hs* PCDH15 CD1-1 (without exon 12a, unless indicated) were split into their respective EC repeats prior to alignment, which was done using the ClustalW algorithm^130^ within Geneious^131^ to determine the percent sequence identity for each EC repeat (Fig. S1 and S3A). Aligned sequences were loaded into JalView^132^ and colored based on sequence conservation with a 45% conservation threshold. Comparison of EC repeats within the human PCDH15 protein was carried out by aligning individual EC repeats to each other using ClustalW on Geneious. The alignment (Fig. S2) was imported into the Sequence Identity and Similarity (SIAS) server^133^ to obtain the sequence identity matrix in Fig. S3B. Structural comparisons and computation of core RMSD were obtained using COOT^109^.

## DATA AVAILABILITY

Coordinates for all structures presented here have been deposited in the Protein Data Bank with entry codes 6N22 (*mm* PCDH15 EC1-2BAP), 6MFO (*hs* PCDH15 EC1-3 G16D/N369D/Q370N), 6N2E (*hs* PCDH15 EC1-3 G16D/N369D/Q370N + *mm* CDH23 EC1-2 T15E), 5ULY (*hs* PCDH15 EC2-3), 6EB5 (*hs* PCDH15 EC2-3 V250N), 6E8F (*hs* PCDH15 EC3-5 CD2-1), 5W1D (*mm* PCDH15 EC4-7), 6BXU (*mm* PCDH15 EC5-7 I582T), 6BWN (*mm* PCDH15 EC6-7), 5TPK (*mm* PCDH15 EC7-8 V875A), and 6EET (*mm* PCDH15 EC9-MAD12). Other raw data and data sets generated and/or analyzed during the current study are available from the corresponding author on reasonable request.

## ACKNOWLEDGEMENTS

We thank Michael Hammel and Daniel Rosenberg for data collection and initial SEC-SAXS data analysis, Marina Bakhtina for assistance with AUC experiments, Benjamin Donovan for assistance with cloning, Michelle E. Gray for maintenance of the Expi293 cell line, as well as members of the Sotomayor research group for their feedback, training, and discussions. This work was supported by the Ohio State University, by the National Institutes of Health – National Institute on Deafness and Other Communication Disorders (NIH/NIDCD R01 DC015271), and by the National Science Foundation through XSEDE (XRAC MCB140226). Simulations were performed at the TACC-Stampede, PSC-Bridges, and OSC-Owens (PAS1037) supercomputers. Use of APS NE-CAT beamlines was supported by NIH (P30 GM1241653 and S10 RR029205) and the Department of Energy (DE-AC02-06CH11357) through grants GUP 40277, 49774, and 59521. SAXS data collection at the Advanced Light Source SIBYLS beamline was funded through the DOE BER Integrated Diffraction Analysis Technologies program and NIH grants NIGMS P30 GM124169 ALS-ENABLE and S10OD018483. B.L.N. was supported by a 2017 Cellular, Molecular, and Biochemical Sciences Program training grant fellowship. P.D. and R.A.-S. were Pelotonia fellows. L.N.W. received summer support from The Ohio State University Mayer scholarship.

## AUTHOR CONTRIBUTIONS

D.C. did cloning, protein expression, purification, crystallization, and X-ray crystal structure solution and refinement of *hs* PCDH15 EC1-3 G16D/N369D/Q370N, *hs* PCDH15 EC1-3 G16D/N369D/Q370N + CDH23 EC1-2 T15E (assisted by P.D.), *hs* PCDH15 EC2-3 V250N, and *hs* PCDH15 EC3-5 CD2-1 (assisted by E.T.). D.C. designed and did initial tests with all mutations targeting the PCDH15 X-dimer, and performed all AUC experiments and analyses. D.C. and Y.N. did MALS experiments (assisted by P.D. and E.T.). Y.N. did cloning, protein expression, purification, crystallization and X-ray crystal structure solution and refinement of *mm* PCDH15 EC1-2BAP and *mm* PCDH15 EC9-MAD12 (mammalian expression). Y.N. did cloning and initial protein expression of PCDH15’s full-length ectodomain, which was further produced by D.C. and E.T. for MALS experiments. B.L.N. performed cloning, protein expression, purification, crystallization, and X-ray crystal structure solution and refinement for *mm* PCDH15 EC5-7 (I582T). B.L.N., C.K., and M.S. performed cloning, protein expression, purification, and crystallization of *mm* PCDH15 EC6-7. B.L.N. solved and refined the *mm* PCDH15 EC6-7 X-ray crystal structure. L.N.W. and M.S. did cloning, protein expression, purification, crystallization, and X-ray crystal structure solution and refinement of *hs* PCDH15 EC2-3. C.K. and M.S. did cloning, protein expression, purification, crystallization, and X-ray crystal structure solution for *mm* PCDH15 EC4-7. B.L.N., C.K., and M.S. refined the *mm* PCDH15 EC4-7 X-ray crystal structure. C.C. and M.S. did cloning, protein expression, purification, crystallization, and X-ray crystal structure solution and refinement of *mm* PCDH15 EC7-8 V875A. Cloning of *mm* PCDH15 EC9-MAD12 for bacterial expression was done by R.A.-S. P.D. and D.C. prepared samples for SAXS experiments, which were analyzed by R.A.-S. Models of multi-structure PCDH15 ectodomains were assembled by B.L.N. and M.S. Simulations were designed, carried out, and analyzed by D.C., Y.N., B.L.N., and M.S. Manuscript and figures were prepared by D.C., Y.N., B.L.N., P.D., R.A-S., and M.S. with feedback from all co-authors. B.L.N. prepared all movies. M.S. trained, supervised, and assisted co-authors in experimental design, cloning, crystal fishing/cryo-cooling, structure refinement and deposition, and data analysis.

## COMPETING FINANCIAL INTERESTS

M.S. is co-inventor on a patent application that describes the use of modified PCDH15 proteins for gene therapy. The authors declare no other competing financial interests.

