## Supplement for "Structural determinants of protocadherin-15 elasticity and function in inner-ear mechanotransduction"

July 2019

**Supplementary Discussion**

*Assignment of Crystallographic Ions*

Inspection of 2Fo-Fc electron density maps during initial stages of refinement depicted positive electron densities at site 1, 2, and 3 of the linker regions in all structures with the exception of *hs* PCDH15 EC3-5 CD2-1 (missing density at site 1 of all chains and site 2 of chain C), *mm* PCDH15 EC5-7 I582T (missing density at sites 1 and 2 of EC5-6 linkers), *mm* PCDH15 EC4-7 (missing density at sites 1 and 2 of EC5-6 linkers), and *mm* PCDH15 EC9-12 (missing density at EC9-10 and 11-12 linker). Unassigned densities at sites 1, 2, and 3 at the EC repeat linker regions were all initially modeled as $Ca^{2+}$ ions and validated in all structures based on compatibility with the final 2Fo-Fc map, an analysis of distances to coordinating atoms, and an evaluation of B factor values of the ion and surrounding residue atoms. However, there were a few cases in which assignment was difficult:

1) As opposed to the WT, site 1 at the *hs* PCDH15 EC2-3 V250N linker region has positive Fo-Fc density. We tried to assign a $Mg^{2+}$ ion given that the crystallization condition had 150 mM $Mg^{2+}$, but it resulted in a positive Fo-Fc density near the ion after subsequent refinement by REFMAC. Thus, this site was assigned to a $Ca^{2+}$ ion, which was more compatible with the 2Fo-Fc and Fo-Fc electron density maps.

2) In the *hs* PCDH15 EC3-5 CD2-1 structure, initial assignment of $Ca^{2+}$ at site 2 of the EC3-4 linker region of chain A resulted in a positive value of the Fo-Fc density at the location of the ion. The protein solution buffer had 5 mM $Ca^{2+}$ and 50 mM $Na^{+}$ while the crystallization buffer had 200 mM LiCl. Thus, we tried placing a $Li^{+}$ ion, which resulted in a large positive value of Fo-Fc density at the site of the ion. In contrast, $Na^{+}$ at the location was compatible with the 2Fo-Fc and Fo-Fc electron density maps, its B factor was similar to that of surrounding atoms, and it is also the cation with the closest ionic radius to $Ca^{2+}$. Thus, a $Na^{+}$ ion was placed at this site in the final model for the structure (Fig. 3C).

3) In the *mm* PCDH15 EC4-7 structure, the EC4-5 linker initially had a $Ca^{2+}$ assigned at site 2, but the B factor was large compared to surrounding atoms. The protein purification buffer contained 20 mM Tris HCl pH 8.0, 150 mM NaCl, 50 mM KCl, and 2 mM $CaCl_2$, while the crystallization buffer had 2.0 M Magnesium Acetate. The $Ca^{2+}$ ion was first replaced by an $Mg^{2+}$ ion but the B factor was still significantly larger than surrounding atoms. The $Mg^{2+}$ ion was replaced by a $K^{+}$ ion, which was in good agreement with the 2Fo-Fc and Fo-Fc electron density maps and had a B factor value more similar to local atoms. The $K^{+}$ ion was kept at site 2 in the EC4-5 linker in the final model.

4) In the *mm* PCDH15 EC6-7 structure, initial assignment of $Ca^{2+}$ at site 1 of the EC6-7 linker resulted in a positive value of the Fo-Fc density at the location of the ion. Since the purification buffer contained 150 mM KCl, we tried placing a $K^{+}$ ion, which was compatible with the 2Fo-Fc and Fo-Fc electron density maps and its B factor was similar to that of surrounding atoms. Thus, the $K^{+}$ ion was kept at this site in the final model.

*Structural modeling of protein residues with poor electron density*

Protein chains were generally modeled using a cutoff of 1.5rmsd for the contour level of the 2Fo-Fc map in COOT. In some loops, weak electron density was observed at a contour level of 1rmsd, which allowed us to fit the residues in the density, but resulted in higher B-factor values for those regions. In some of our structures we observed no density for the N- and C- terminal loops even at a contour level of 0.5rmsd. Such regions of the protein were not built. In addition, some regions of our structures did not have clear density and were not built either. Missing residues in our models include: (1) β-strands F and G of the CDH23 EC2 repeats and the BC loop of one of the PCDH15 protomers in the *hs* PCDH15 EC1-3 G16D/N369D/Q370N + *mm* CDH23 EC1-2 T15E structure; (2) the cysteine loop, parts of the AB loop containing a $3_{10}$ helix and the EF loop in the *hs* PCDH15 EC1-3 G16D/N369D/Q370N structure; (3) the BC loop in all four protomers and the FG loop in two of the protomers in the *hs* PCDH15 EC2-3 structure; (4) the cysteine loop, the BC loop, and the FG loop in the *hs* PCDH15 EC2-3 V250N structure; (5) the BC loop in all protomers and β-strand A along with the connecting loop in chain C of the *hs* PCDH15 EC3-5 CD2-1 structure; (6) the BC loop in the *mm* PCDH15 EC4-7 structure; (7) the EF loop in the *mm* PCDH15 EC7-8 V875A structure.

*Molecular mechanisms of inherited deafness*

Structural models of PCDH15 show the location of three segments that are deleted and associated with inherited deafness and Usher syndrome. The first in-frame deletion (p.E272 to p.Q509; Fig. 5I) causes Usher syndrome (moderate to profound deafness, vestibular dysfunction, and impaired vision) and takes out a large part of EC3 (β-strands B to G), all EC4, and a short piece of EC5 (β-strand A). Such large protein deletion may cause misfolding and poor localization, although it is possible that the resulting protein product folds with a hybrid repeat formed by β-strand A of EC3 and β-strands B to G of EC5, thus shortening the ectodomain by two EC repeats. This shortened PCDH15 might be mechanically compromised and might be unable to form the X-dimer mediated by EC2-EC3 linker region. A second in-frame deletion (p.G936-p.K982; Fig. 5J) causes deafness and vestibular dysfunction in mice (*Av*-6J) and takes out β-strands C, D, E, and part of β-strand F in EC9[136,137]. However, functional tip-links are observed in hair cells from these mice, suggesting that the protein product is properly folded and trafficked, but perhaps unstable[137]. The last in-frame deletion involves elimination of a single residue (p.V767)[138], which results in non-syndromic deafness and might affect the register and stability of β-strand F in EC7, thus compromising the mechanical stability of PCDH15[89], but not enough to alter vestibular function.

Missense mutations implicated in inherited deafness and located throughout the PCDH15 ectodomain (15 sites) can be segregated in at least four groups (Table S7). In the first one we included two mutations (p.I108N and p.R113G) that are known to interfere with the PCDH15-CDH23 handshake bond[21,33,34,71]. Three mutations that presumably alter $Ca^{2+}$ binding at site 3 directly (p.D157G[139] and p.D989G[140] at the EC1-2 and EC8-9 linker regions, respectively) or indirectly (p.G241D[71] at the EC2-3 linker region) are in the second group. Four more mutations (p.L408P[141], p.V507D[142], p.G1130R[143], p.S1267P[141]) are likely to disrupt proper folding, either because side chains involved in hydrogen bonding at β strands are mutated to proline residues, or because side chains pointing towards the hydrophobic core of an EC repeat are replaced by large hydrophilic residues. Intriguingly, our last group includes six mutations for which an evaluation of their structural impact is difficult. Four of them (p.G79R, p.R257H, p.P294L, and p.R941C) might not be causative of inherited deafness[143]. The p.R1013H mutation, which has been shown to cause profound non-syndromic hearing impairment[144], is located near the PCDH15 EC9-10 kink, yet how it could perturb PCDH15 structure and dynamics is unclear. The last mutation, p.D414A (Fig. 3B), has been suggested to be positively selected in East-Asian populations[134], but it has also been associated to inherited deafness[135]. Prior structural analyses suggest that this polymorphism, common in many South Asian exomes, is unlikely to cause a severe loss-of-function phenotype[44]. Our *hs* PCDH15 EC3-5 CD2-1 structure provides a view of the p.D414 site in the context of the CD2-1 isoform that includes a segment encoded by exon 12a: p.V(414+1)PPSGVP(414+7). In this structure, the side-chain of p.D414 is pushed closer to the protomer and to p.K410 when compared to the *hs* PCDH15 EC3-5 structure without exon 12a (PDB: 5T4M)[44]. It is possible that a p.D414:p.K410 salt bridge rigidifies the EC3-4 linker region, or changes its affinity for $Ca^{2+}$ in the context of the CD2-1 isoform only, but how losing the p.D414 side chain could cause an evolutionary advantage or a loss-of-function phenotype remains unclear.

*Computational modeling of systems with low $Ca^{2+}$*

Tip links are found in fluid environments with low $Ca^{2+}$. In the cochlear endolymph, the bulk $Ca^{2+}$ concentration ranges from 20 to 40 μM[52,53] and the sub-tectorial space $Ca^{2+}$ concentration possibly goes up to 300 μM[55]. Vestibular endolymph $Ca^{2+}$ concentration might be ~200 μM. Experimental measurements of *mm* CDH23 EC1-2 $Ca^{2+}$-binding affinities[42] suggest that sites 1, 2, and 3 at this canonical linker region have dissociation constants $K_{D1}$ ~71 μM, $K_{D2}$ ~44 μM, and $K_{D3}$ ~5 μM, respectively. Dissociation constants for non-canonical linker regions in PCDH15 might be larger[44]. Therefore, it is possible that some, or even all sites are not occupied by $Ca^{2+}$ ions under some physiological conditions. In all cases, however, we expect that dissociation constants will follow the same trend with $K_{D1} > K_{D2} > K_{D3}$. Hence, to mimic low-$Ca^{2+}$ conditions in simulations, we removed ions from binding sites sequentially. We removed first all the $Ca^{2+}$ ions at site 1 to build the 2 $Ca^{2+}$ system (simulations

S4a-d). To build the 1 $Ca^{2+}$ system (simulations S5a-d), the $Ca^{2+}$ ions at site 1 and 2 were removed. Lastly, to build the 0 $Ca^{2+}$ system, all $Ca^{2+}$ ions were removed (simulations S6a-d; Table S8).

**Movies**

*Supplementary Movie 1: Forced unbending, unrolling, and unfolding in simulations of the hs PCDH15 EC1-MAD12 CD1-1 + CDH23 EC1-2 model.* Stretching of the complex at 0.1 nm/ns (simulation S1d in Table S8, 0 – 191.9 ns) results in straightening of the PCDH15 EC9-10 linker region with a twisting of the PCDH15 protomer. As the trajectory continues the PCDH15 EC5-6 and EC9-10 linker regions stretch without unfolding of the repeats. In the final moments of the simulation PCDH15 MAD12 peels away (unrolls) from EC11 and begins to unfold from the C-terminus without PCDH15 and CDH23 unbinding. Protein is depicted in cartoon representation with a rough surface envelope (PCDH15 – purple; CDH23 – blue). $Ca^{2+}$ ions are shown as green spheres. Water molecules and other atoms are not shown for clarity.

*Supplementary Movie 2: Forced unbending, unrolling, and unfolding in simulations of the mm PCDH15 EC1-MAD12 CD2-1 + mm CDH23 EC1-3 model.* Stretching of the complex at 0.1 nm/ns (simulation S2d in Table S8, 0 – 251.6 ns) results in straightening of the PCDH15 EC9-10 linker with a twisting of the PCDH15 protomer. Later in the trajectory the PCDH15 EC5-6 and EC9-10 linkers stretch without unfolding of the repeats. The PCDH15 MAD12 peels away from EC11 and begins to unfold from the C-terminus without PCDH15 and CDH23 unbinding as the simulation ends. System shown as in supplementary movie 1.

*Supplementary Movie 3: Forced unbending, unrolling, and unfolding in simulations of the mm PCDH15 EC9-MAD12 dimer.* Stretching of the complex at 0.1 nm/ns (simulation S11d – 95.8 ns) results in straightening of the PCDH15 EC9-10 linker regions, followed by unrolling of MAD12 and unfolding at the C-terminus end of PCDH15. System shown as in supplementary movie 1.

*Supplementary Movie 4: Forced unbinding in simulations of the hs $(PCDH15\ EC1\text{-}5)_2$ + $(CDH23\ EC1\text{-}2)_2$ model.* Stretching of the complex at 0.1 nm/ns (simulation S3d in Table S8, 0 – 95.6 ns) results in straightening of PCDH15 EC3-5 protomers coming into contact, squeezing of CDH23 protomers, and in unbinding of both CDH23 protomers from PCDH15 without unfolding of any EC repeats. The X-dimer interface mediated by PCDH15 EC2-3 was partially disrupted towards the end of the simulation trajectory. System shown as in supplementary movie 1.

*Supplementary Movie 5: Forced unbinding in simulations of the mm $(PCDH15\ EC1\text{-}5)_2$ + $(CDH23\ EC1\text{-}3)_2$ model.* Stretching of the complex at 0.1 nm/ns (simulation S7d in Table S8, 0 – 88.3 ns) results in straightening of PCDH15 EC3-5 protomers coming into contact, squeezing of CDH23 protomers, and in unbinding of both CDH23 protomers from PCDH15 without unfolding of any EC repeats. The X-dimer interface mediated by PCDH15 EC2-3 was partially disrupted towards the end of the simulation trajectory. System shown as in supplementary movie 1.

*Supplementary Movie 6: Forced unbinding in simulations of the hs $(PCDH15\ EC1\text{-}5)_2$ + $(CDH23\ EC1\text{-}2)_2$ model in the absence of $Ca^{2+}$.* Stretching of the complex at 0.1 nm/ns (simulation S6d in Table S8, 0 – 142.9 ns) first resulted in lengthening of all EC linker regions without unfolding of domains. The X-dimer interface mediated by PCDH15 EC2-3 broke before unbinding of CDH23 from PCDH15. The PCDH15 and CDH23 interaction unbound without unfolding of EC repeats at the end of the trajectory. System shown as in supplementary movie 1.

*Supplementary Movie 7: Tour of the hs PCDH15 EC1-MAD12 + CDH23 EC1-2 model.* System is shown first within the solvation box (PCDH15 – purples; CDH23 – blues; $Ca^{2+}$ ions – green; SMD atoms - red). Zoom in details focus on: The PCDH15 + CDH23 interaction with emphasis on a salt bridge important in the maintenance of the complex (p.R113:E77); the PCDH15 EC2-3 X-dimer; the non-canonical PCDH15 EC5-6 linker region; the straightened $Ca^{2+}$-free PCDH15 EC9-10 linker region; the PCDH15 EC11-MAD12 *cis* dimer; and the MAD12 fold.

*Supplementary Movie 8: Tour of the mm PCDH15 EC1-MAD12 + CDH23 EC1-3 model.* System is shown first within the solvation box (PCDH15 – purples; CDH23 – blues; $Ca^{2+}$ ions – green; SMD atoms - red). Zoom in details focus on: The interaction between CDH23 EC1 $3_{10}$ helices (between β-strands C and D) with emphasis on the "interlocking" salt bridges (p.R53:E50) and residues p.E49 which cap the helix dipoles; and the PCDH15 CD2-1 structure highlighting the enlarged BC loop in EC4 (p.V(414+1)PPGGVP(414+7)).

*Supplementary Movie 9: Forced unbending, unrolling, and unfolding in simulations of the hs (PCDH15 EC1-MAD12)$_2$ + hs (CDH23 EC1-2)$_2$ model.* Stretching of the complex at 0.1 nm/ns (simulation S8d in Table S8, 0 – 122.3 ns) results in straightening of the PCDH15 ectodomains with lengthening of the EC5-6 and EC9-10 linker regions. As the simulation progressed the PCDH15 MAD12s began to unfold from their C-terminal ends. The PCDH15 MAD12s eventually unrolled away from EC11 while unfolding continued. The CDH23 protomers did not unbind from PCDH15. System shown as in supplementary movie 1.

*Supplementary Movie 10: Forced unbending, unrolling, and unfolding in simulations of the mm (PCDH15 EC1-MAD12)$_2$ + mm (CDH23 EC1-3)$_2$ model.* Stretching of the complex at 0.1 nm/ns (simulation S9d in Table S8, 0 – 135.5 ns) results in straightening of the PCDH15 ectodomains with lengthening of the EC5-6 and EC9-10 linker regions. As the simulation progressed the PCDH15 MAD12s began to unfold from their C-terminal ends. The PCDH15 MAD12s eventually unrolled away from EC11 while unfolding continued. The CDH23 protomers did not unbind from PCDH15. System shown as in supplementary movie 1.

1
2

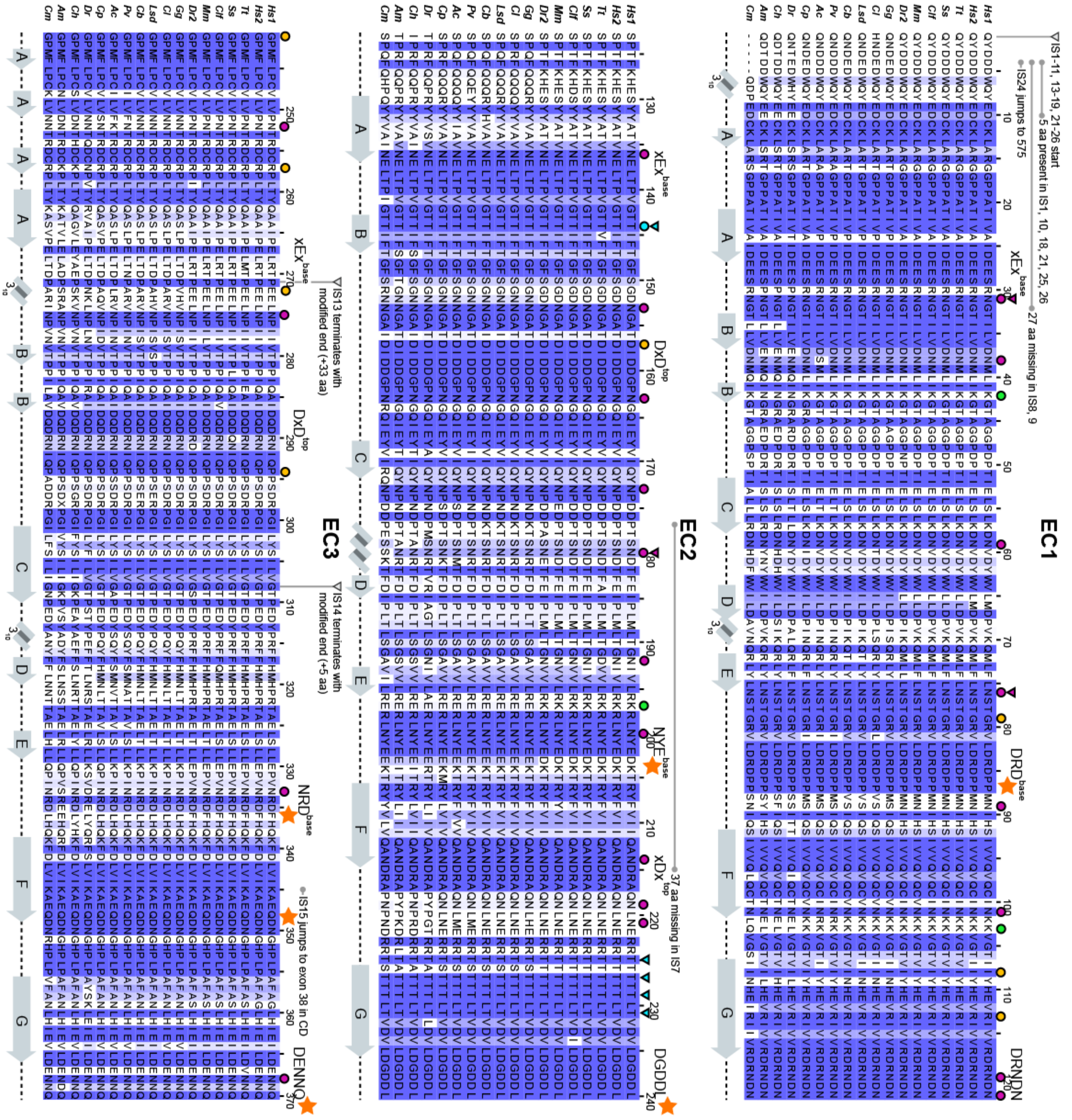

EC1
EC2
EC3
IS1-11, 13-19, 21-26 start
5 aa present in IS1, 10, 18, 21, 25, 26
IS24 jumps to 575
27 aa missing in IS8, 9
37 aa missing in IS7
IS13 terminates with modified end (+33 aa)
IS14 terminates with modified end (+5 aa)
IS15 jumps to exon 38 in CD
Hs1
Hs2
Tt
Ss
Clf
Mm
Dr2
Gg
Cl
Lsd
Cb
Pv
Ac
Gp
Dr
Ch
Am
Cm

1

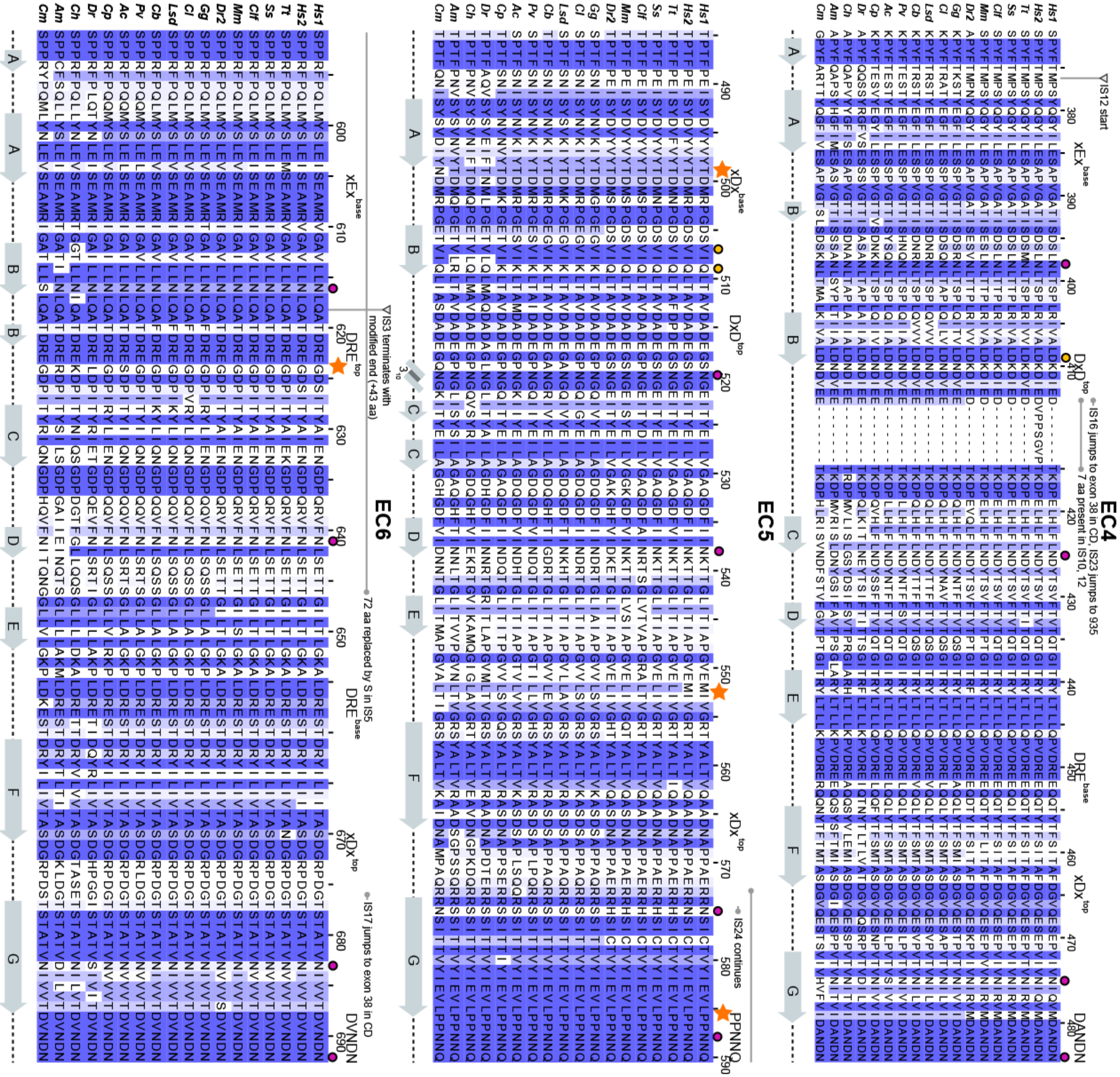
EC4
EC5
EC6
IS12 start
IS16 jumps to exon 38 in CD, IS23 jumps to 935
7 aa present in IS10, 12
IS24 continues
IS3 terminates with modified end (+43 aa)
72 aa replaced by S in IS5
IS17 jumps to exon 38 in CD

1

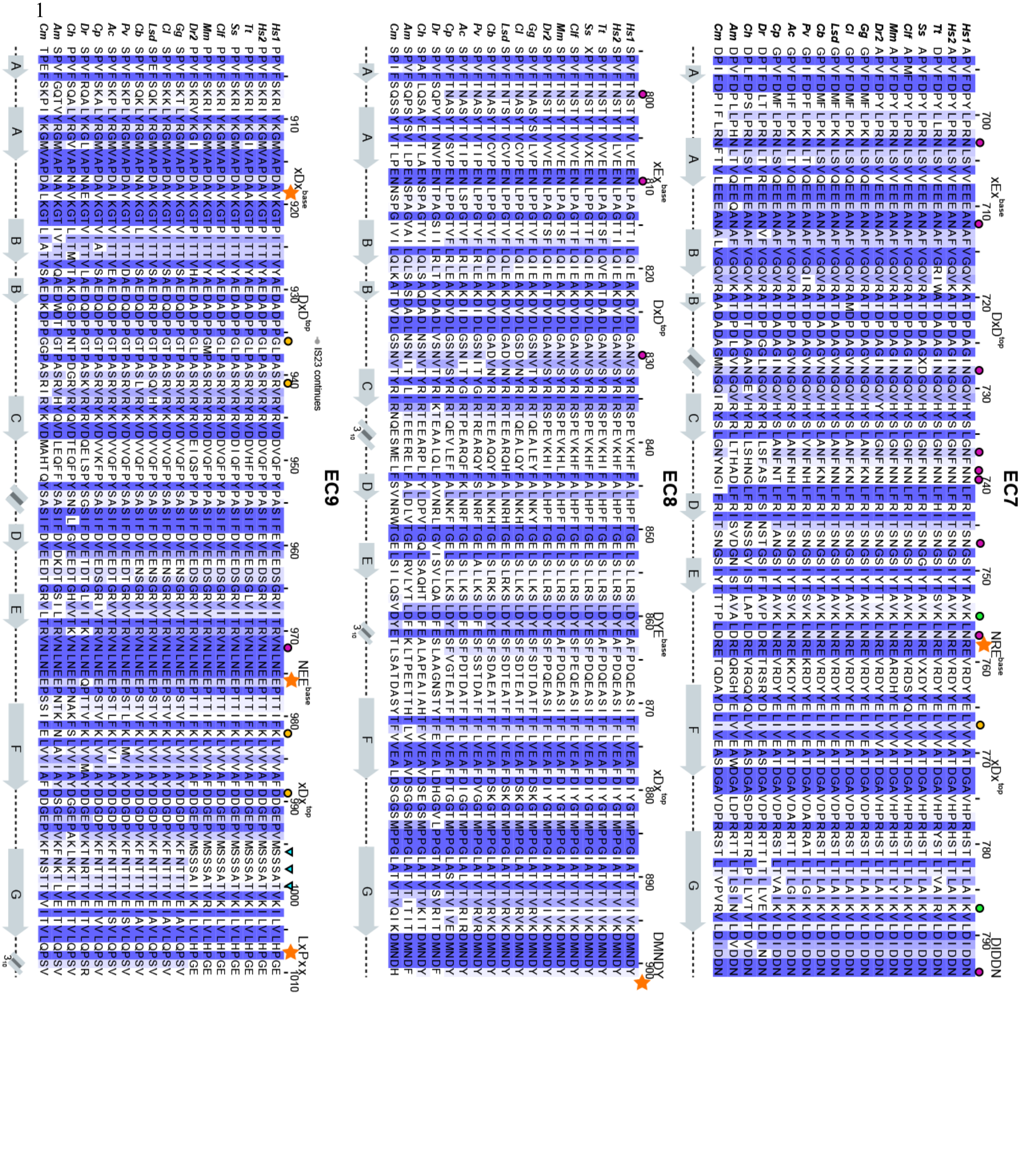

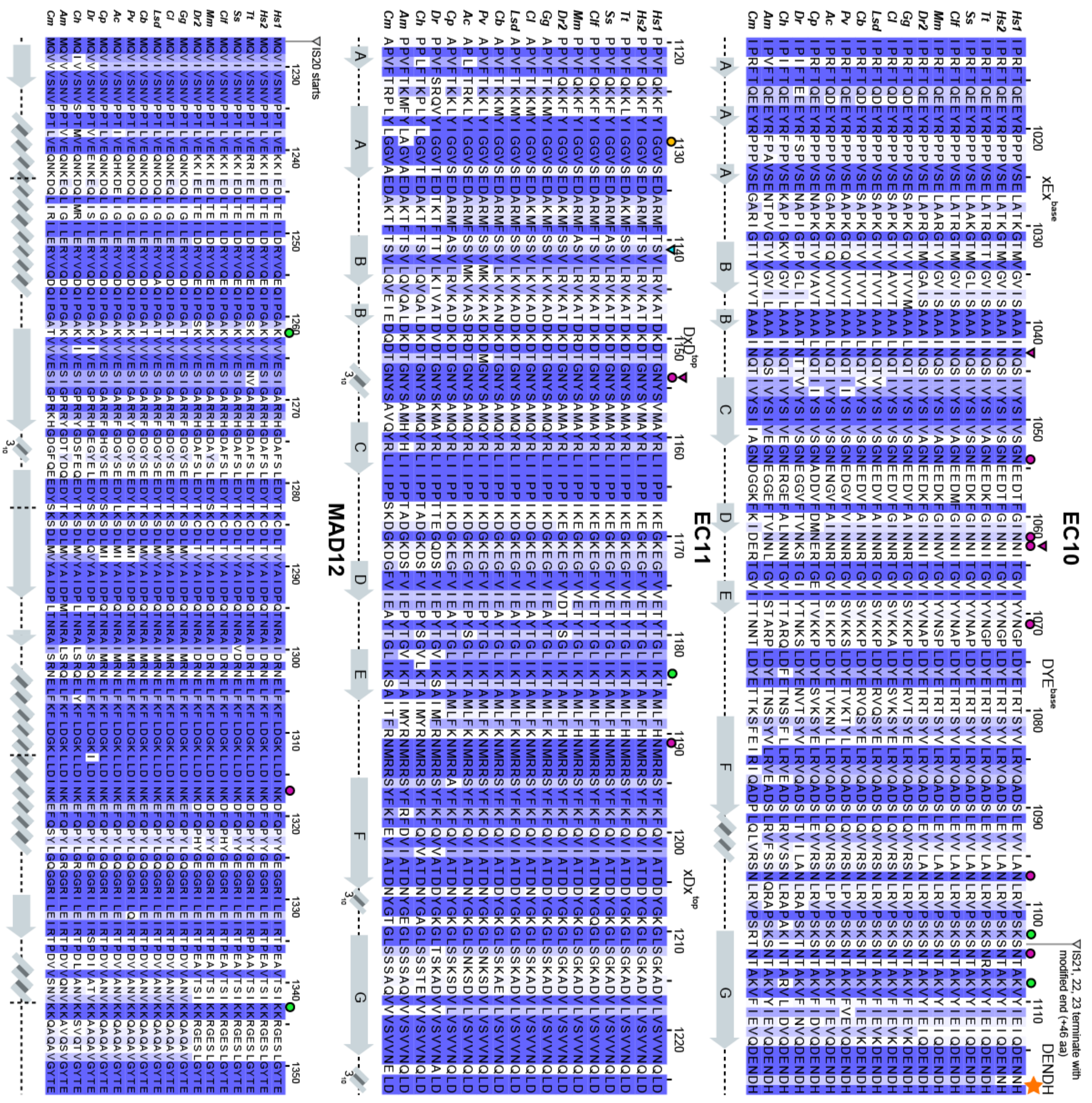
EC10
EC11
MAD12
IS20 starts
IS21, 22, 23 terminate with modified end (+46 aa)

**Figure S1. Sequence alignment of PCDH15 ectodomains from different species.** Multiple sequence alignments comparing sequences of each PCDH15 EC repeat from 17 different species and two different human isoforms (Table S1). Each alignment is colored by sequence similarity with white being the lowest similarity and blue being the highest (see methods). Some columns may not be colored if deletions or changes to residue type are present (e.g. polar amino acid to hydrophobic). $Ca^{2+}$-binding motif sequences are labeled on top of each alignment, with orange stars indicating atypical motifs. Disease mutation sites are highlighted with yellow circles. Predicted sites of N- and O-linked glycosylation are marked with red and cyan circles, respectively. Experimentally observed glycosylation sites are marked with red and cyan triangles. Annotations related to various *mm* PCDH15 isoforms (labeled IS-1 to IS-26) are marked with grey triangles above each alignment. UniProt accession numbers for IS1-26 are Q99PJ1-1 to Q99PJ1-26, respectively. Secondary structure elements are displayed underneath each alignment. Species were chosen based on sequence availability and taxonomical diversity.

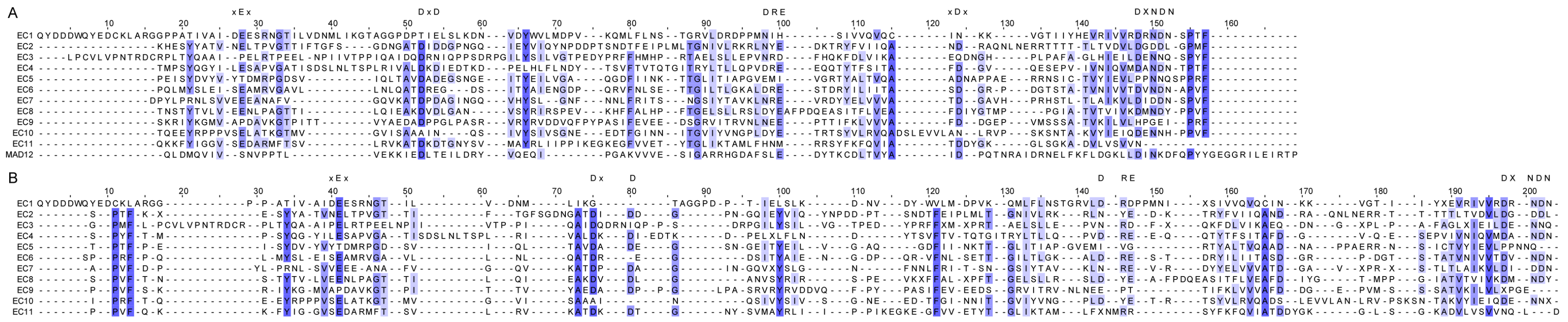

**Figure S2. Sequence alignments for PCDH15 EC repeats and MAD12.** (**A**) Pairwise multiple sequence alignment of *hs* PCDH15 CD1-1 (NP_001136235.1) EC1 to MAD12. (**B**) Structure-based sequence alignment of PCDH15 CD1-1 EC1-EC11 repeats. $Ca^{2+}$-binding motif sequences are labeled on top of the alignments.

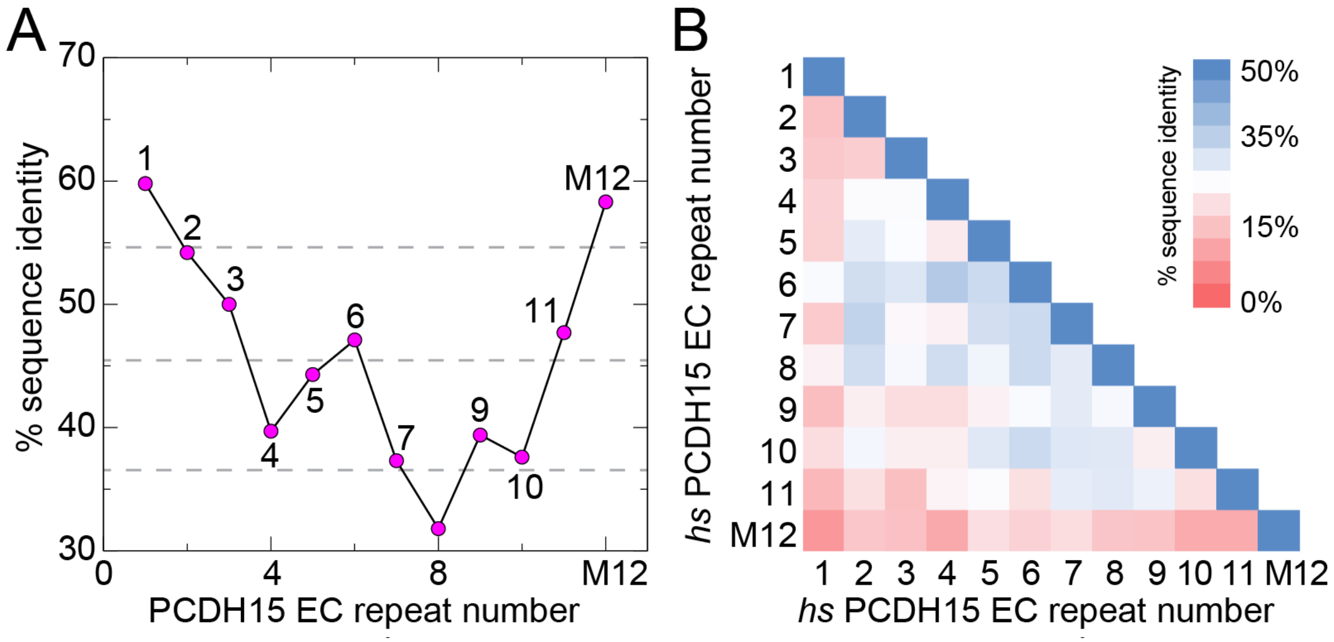


**Figure S3. PCDH15 Sequence identity across species and among repeats within a single species.** (**A**) Plot of percent sequence identity of each PCDH15 EC repeat and MAD12 (M12) among 17 species against repeat number (Table S1). The most conserved regions of the protein include EC1 and MAD12, whereas EC8 is the least conserved. The average sequence identity is 45.6 ± 8.9% (gray dashed lines). (**B**) Heat map of a sequence identity matrix among EC-repeats for *hs* PCDH15 CD1-1 (NP_001136235.1). Sequence alignment from Figure S2A was used as input to compute % sequence identity (see Methods).

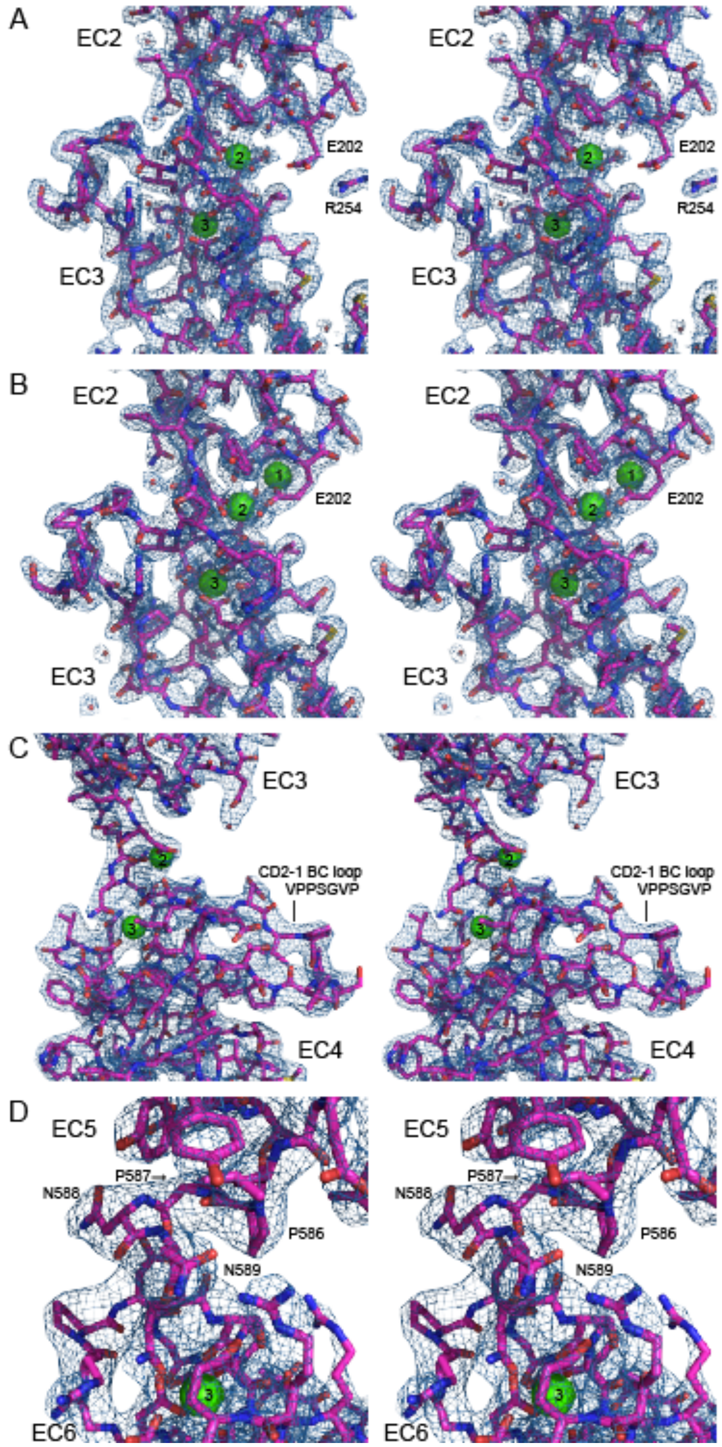


**Figure S4. Electron density maps of distinct structural features in PCDH15.** (**A**) Stereo view of the 2Fo-Fc electron density map (blue mesh) of *hs* PCDH15 EC2-3 WT (PDB: 5ULY; chain D in purple) showing a detail of its linker region with two $Ca^{2+}$ ions (green spheres). Water molecules are shown as red spheres. Residue p.E202 interacts with p.R254 and hence is not involved in $Ca^{2+}$ coordination. (**B**) Similar stereo view for the *hs* PCDH15 EC2-3 V250N structure (PDB: 6EB5; chain A). The linker region has three $Ca^{2+}$ ions with residue p.E202 coordinating the ion at site 1. Notably, the cysteine loop carrying p.R254 is too flexible in the absence of the X-dimer interface and is not seen in this structure. (**C**) Stereo view of the 2Fo-Fc electron density map (shown as in A) for the *hs* PCDH15 EC3-5 CD2-1 structure highlighting the enlarged BC loop in EC4 (p.V(414+1)PPSGVP(414+7)). (**D**) Stereo view of the 2Fo-Fc electron density map (shown as in A) for the EC5-6 linker region in the *mm* PCDH15 EC4-7 structure (PDB: 5W1D). All maps are contoured at 2.0 σ.

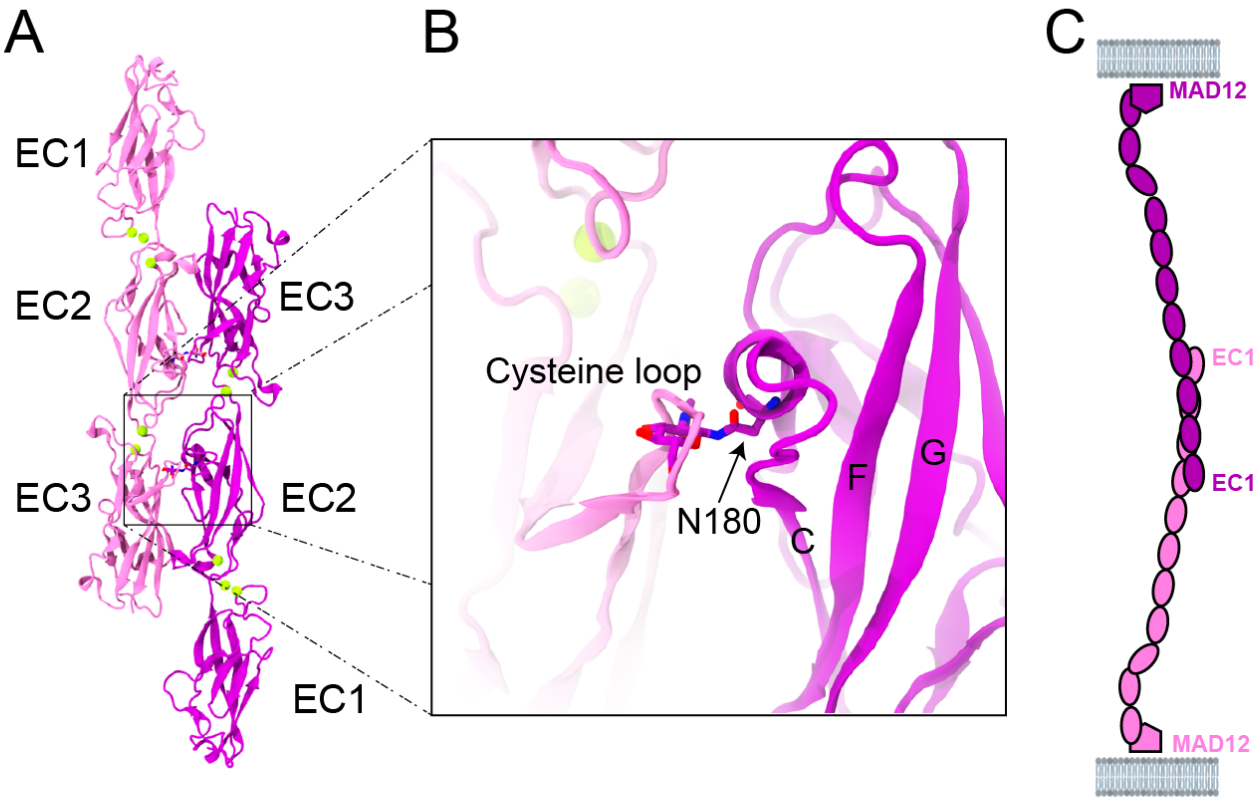


**Figure S5. Glycosylation site at an antiparallel PCDH15 dimer interface.** (**A**) To probe the location and effects of potential glycosylation sites on the antiparallel dimer of *hs* PCDH15 EC1-3 G16D/N369D/Q370N (PDB: 6MFO; protein expressed in bacteria without glycosylation), the glycosylated *mm* PCDH15 EC1-3 structure (PDB: 6CV7; protein expressed in mammalian cells)[38] was superposed on the structure of the antiparallel dimer. Ribbon representation of protomers (mauve and purple) forming an antiparallel *trans* dimer interface using the *mm* PCDH15 EC1-3 structure highlights a steric clash (black box). (**B**) Detail of the cysteine loop clashing with p.N180 carrying a sugar moiety. The same cysteine loop is not visible in the *hs* PCDH15 EC1-3 G16D/N369D/Q370N structure (PDB: 6MFO). (**C**) Schematic representation of a potential antiparallel *trans* PCDH15-PCDH15 tip link if the cysteine loop rearranges and the aperture angle between EC2-3 repeats decreases to avoid steric clashes.

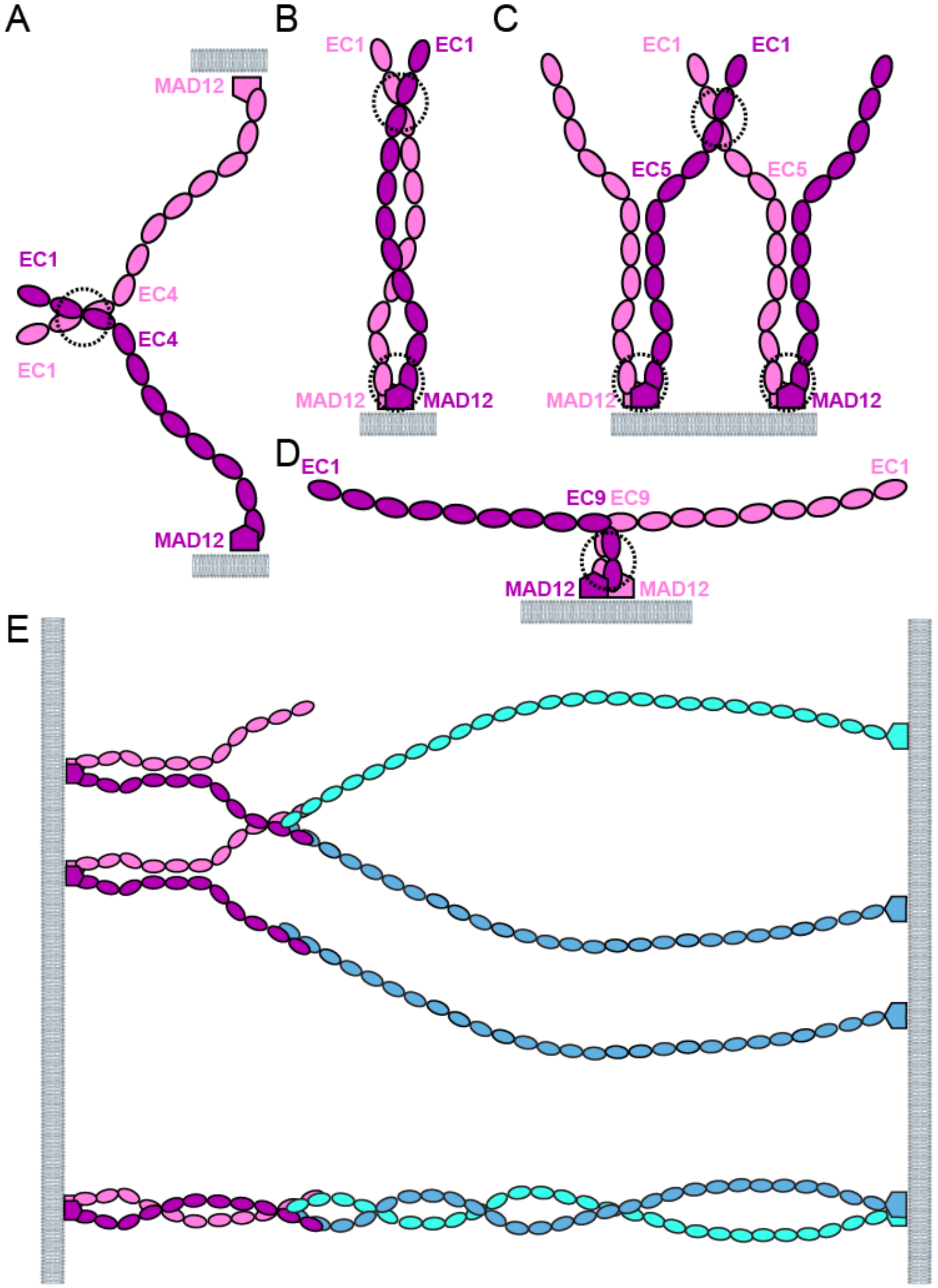


**Figure S6. Schematics of various potential modes of PCDH15 dimerization and interactions with CDH23.** (**A-C**) Modes of *trans* (A) and *cis* (B-C) dimerization that can be mediated by the PCDH15 EC2-3 X-dimer interface presented here and in Dionne et al.[38] Immature transient PCDH15 tip links[30,31] might adopt the *trans* configuration shown in A, similar to the rightmost configuration in Fig. S12D. Dashed circles indicate sites of dimerization. (**D**) Parallel dimerization mediated by MAD12[39,40] along with a fully bent EC9-10 linker[43] results in protomers pointing in opposite directions and parallel to the membrane plane. (**E**) Potential *trans* heterophilic interactions of PCDH15 (mauve and purple) and CDH23 (cyan and blue). Top arrangement could be adopted by kinociliary links[76], while the lower arrangement is compatible with the heterotetrameric tip link[21,33].

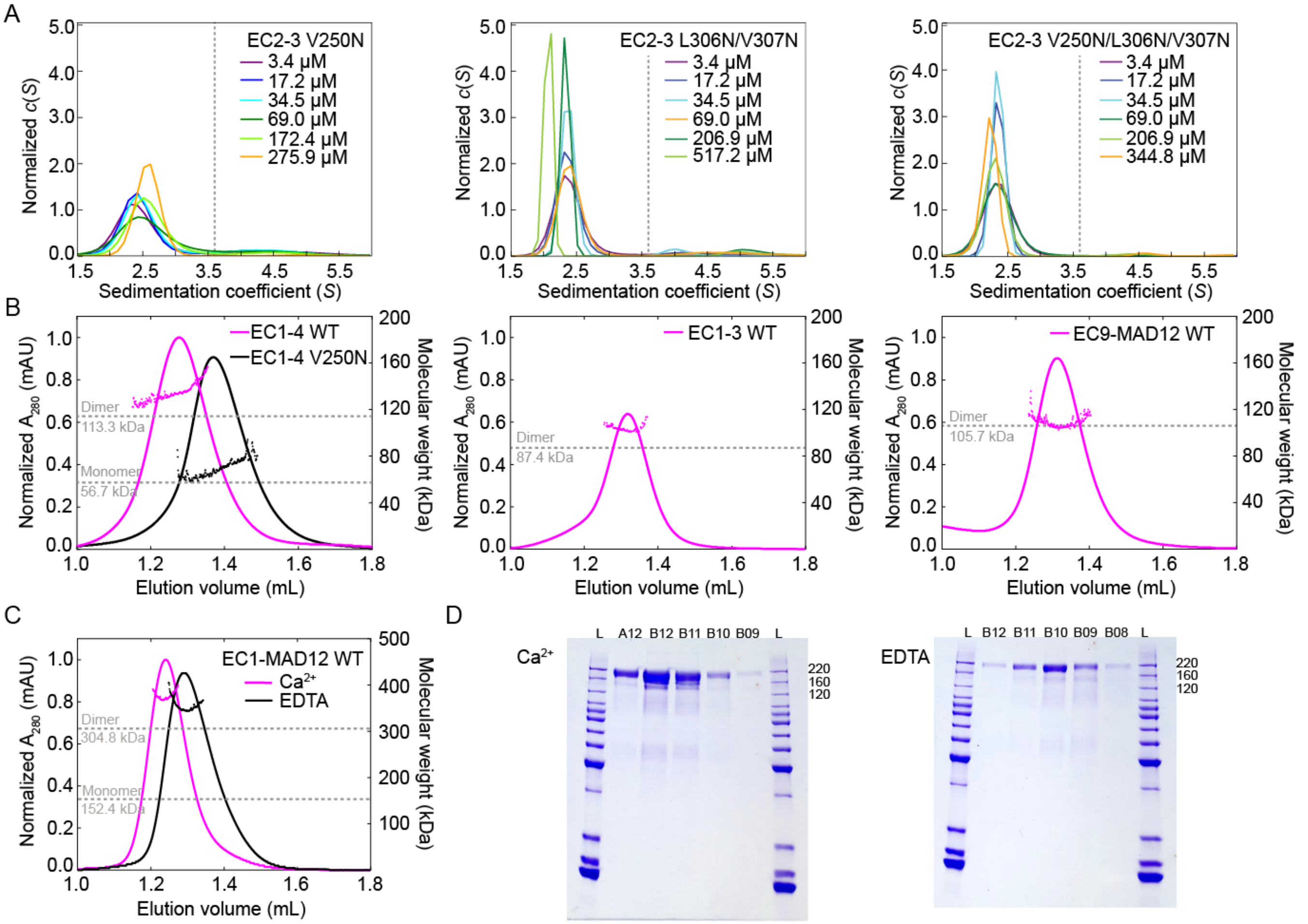


**Figure S7. Dimerization of PCDH15 in solution**. (**A**) Results from AUC experiments for point mutants in the *hs* PCDH15 EC2-3 dimer interface. Plots show normalized sedimentation coefficient distributions *c*(*S*) at varying concentrations of bacterially expressed *hs* PCDH15 EC2-3 p.V250N (left panel), p.L306N/p.V307N (middle), and p.V250N/p.L306N/p.V307N (right). Dotted gray line represents the sedimentation coefficient (*S*) value of WT dimer (Fig. 2C). All mutants are monomeric in solution. (**B**) SEC-MALS results for mammalian expressed *mm* PCDH15 EC1-4 WT and p.V250N mutant (left panel), *mm* PCDH15 EC1-3 WT (middle), and *mm* PCDH15 EC9-MAD12 (right) protein fragments using a Superdex S200 3.2/30 column. Horizontal dashed lines indicate theoretical molar mass for monomer and dimer. (**C**) SEC-MALS results for mammalian expressed *mm* PCDH15 EC1-MAD12 in the presence of $Ca^{2+}$ (magenta) and EDTA (black) using a Superose 6 3.2/30 column. (**D**) Coomassie stained SDS-PAGE of eluted fractions from experiments in C.

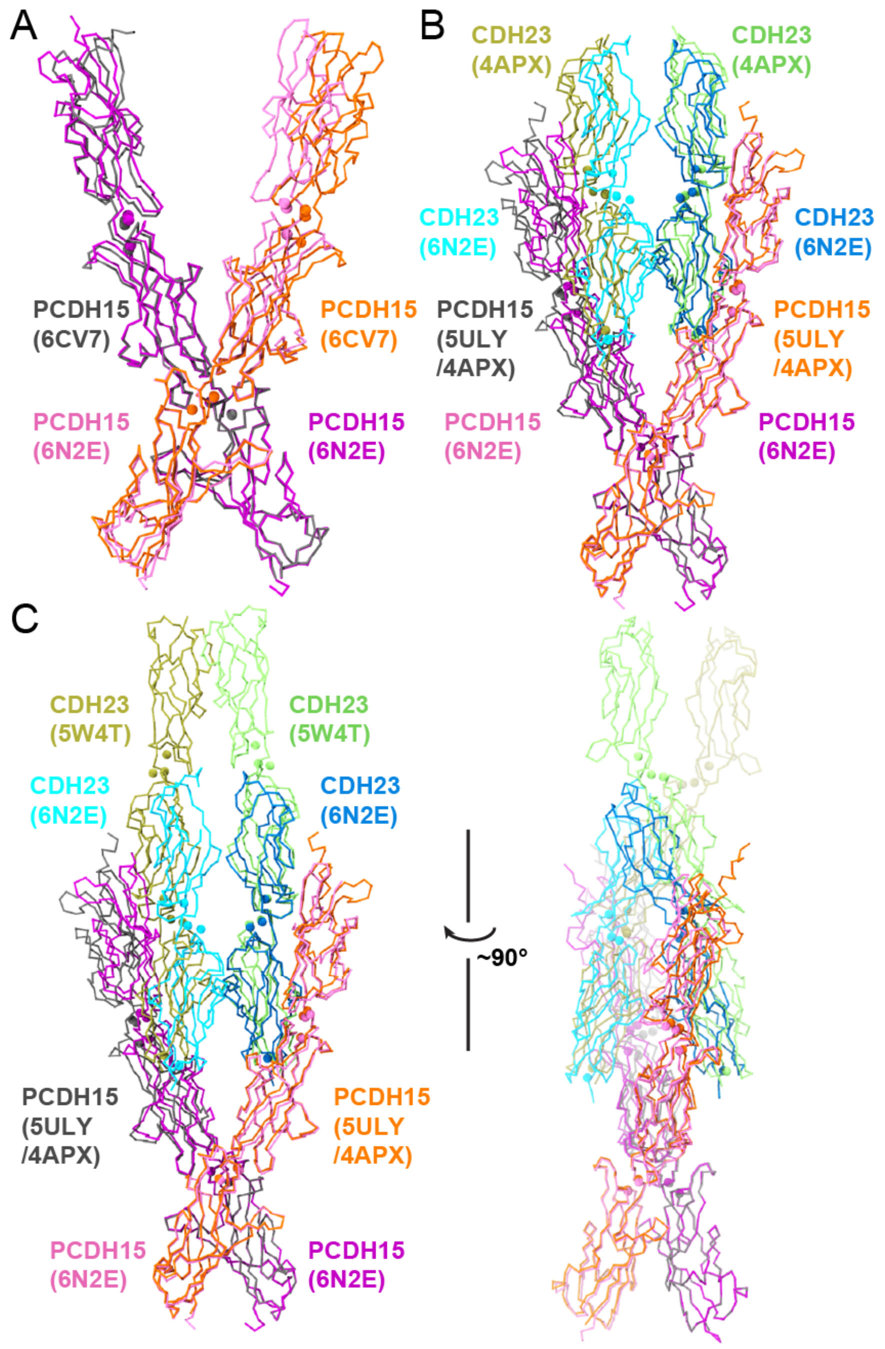


**Figure S8. Structural comparison of PCDH15 X-dimer based tetrameric assemblies.** (**A**) Superposition of *mm* PCDH15 EC1-3 (PDB: 6CV7; gray and orange)[38] and *hs* PCDH15 EC1-3 G16D/N369D/Q370N (PDB: 6N2E; mauve and purple; *mm* CDH23 EC1-2 T15E is not shown). Repeats EC3 from one protomer (gray and purple) were structurally aligned. The "scissor" is less open in the structure where PCDH15 is bound to CDH23 EC1-2 (PDB: 6N2E), with a smaller separation between repeats EC1-2 of the two PCDH15 protomers. (**B**) Superposition of the *hs* PCDH15 EC1-3 G16D/N369D/Q370N (mauve and purple) + *mm* CDH23 EC1-2 T15E (blue and cyan) tetramer complex structure (PDB: 6N2E) with a tetramer model (orange, gray, green, and tan) created by using the X-dimer conformation observed in *hs* PCDH15 EC2-3 (PDB: 5ULY) and the *mm* PCDH15 EC1-2 + *mm* CDH23 EC1-2 handshake interaction (PDB: 4APX)[33]. A similar model was also reported by Dionne et al.[38] In the tetrameric crystal structure, repeat EC1 of PCDH15 from one protomer moves towards the other, thus bringing the two CDH23 EC1-2 molecules closer to each other. (**C**) Superposition of the *hs* PCDH15 EC1-3 G16D/N369D/Q370N (mauve and purple) + *mm* CDH23 EC1-2 T15E (blue and cyan) tetramer complex structure (PDB: 6N2E) with a tetramer model (orange, gray, green, and tan) created by incorporating the *dr* CDH23 EC1-3 structure (PDB: 5W4T)[45] to the tetramer model generated in panel B through alignment of CDH23 EC1-2 protomers.

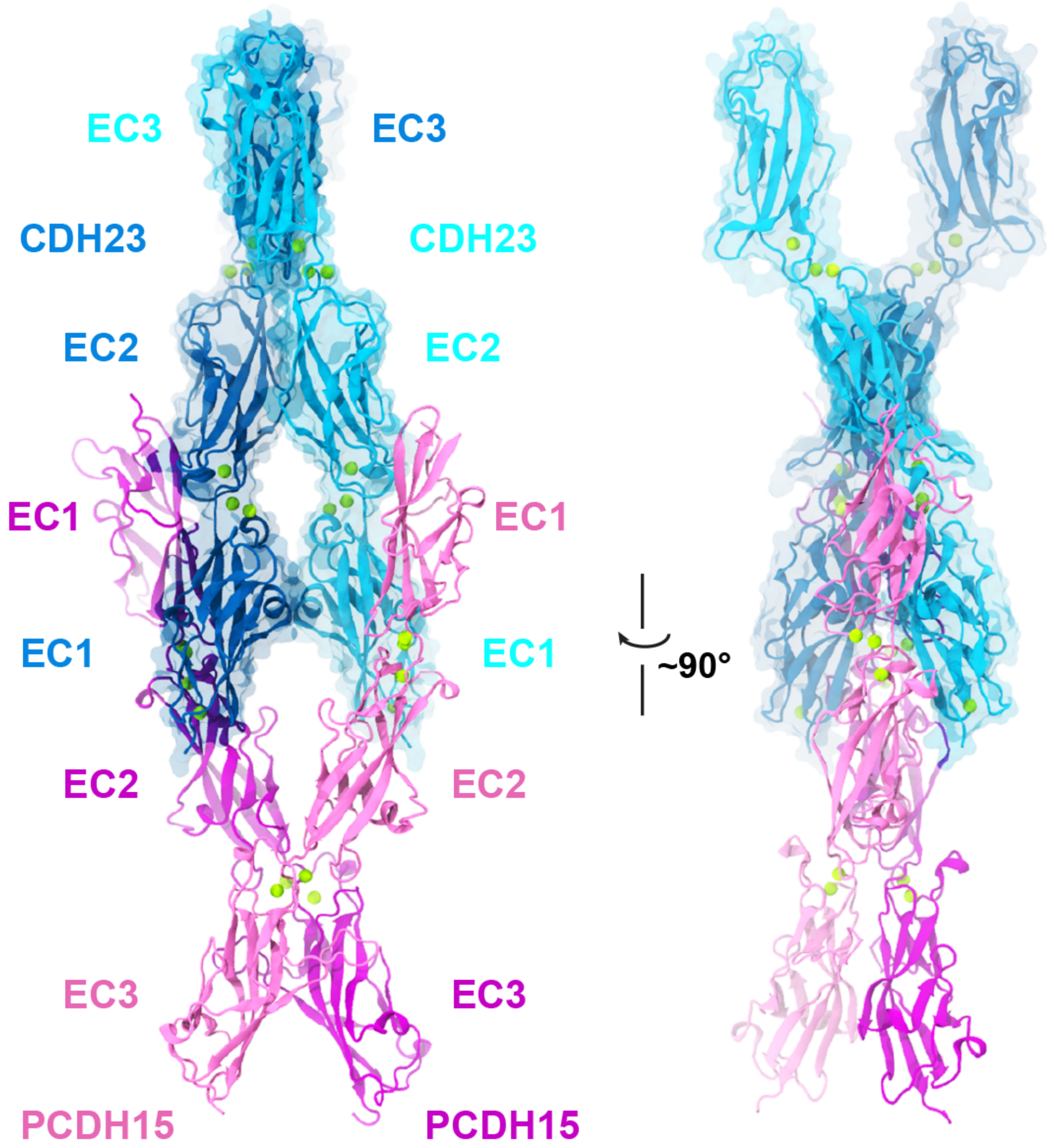


**Figure S9. Structural model of PCDH15 EC1-3 in complex with CDH23 EC1-3.** A model of CDH23 EC1-3 (blue and cyan) in complex with the PCDH15 EC1-3 X-dimer (mauve and purple). $Ca^{2+}$ ions are shown in green. Model was generated by aligning CDH23 EC1-2 repeats from the *dr* CDH23 EC1-3 structure (PDB: 5W4T)[45] and from the *hs* PCDH15 EC1-3 G16D/N369D/Q370N (mauve and purple) + *mm* CDH23 EC1-2 T15E (blue and cyan) tetramer complex structure (PDB: 6N2E). This tetrameric arrangement suggests that CDH23 dimerization involves repeats beyond EC3.

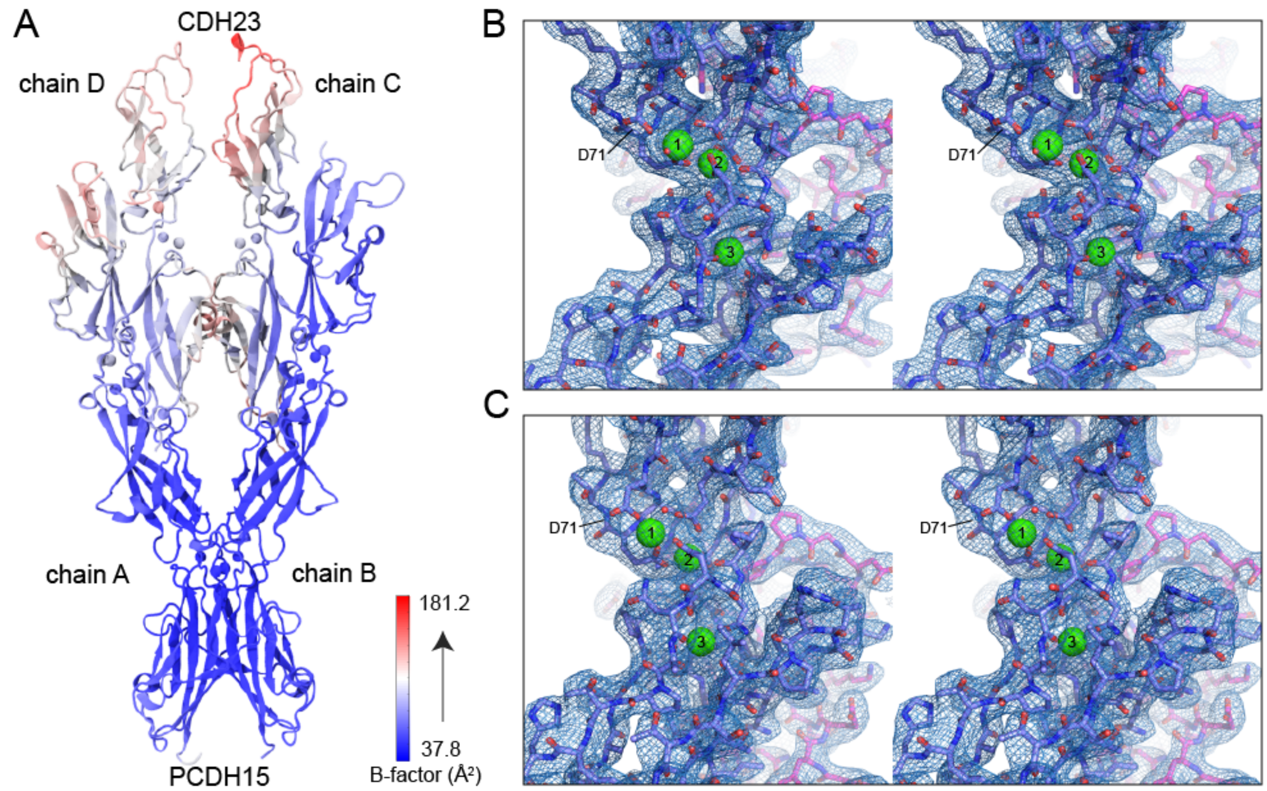


**Figure S10. Flexibility and presence of CDH23 protomers in tetramer structure.** (**A**) Ribbon diagram of the *hs* PCDH15 EC1-3 G16D/N369D/Q370N + *mm* CDH23 EC1-2 T15E structure (PDB: 6N2E) colored by B-factor varying from blue (low) to red (high). The EC2 repeats of CDH23 (chain C and D) and the EC1 repeat of PCDH15 (chain A) have large B factor values (red), indicating inherent flexibility. (**B-C**) Stereo representations of the composite omit electron density map for CDH23 EC1-2 linkers contoured at 0.75σ (blue mesh) for chains C and D, respectively. CDH23 is depicted as blue sticks, PCDH15 is depicted as mauve sticks, $Ca^{2+}$ ions are shown as green spheres. $Ca^{2+}$ ions are clearly seen in the map, corroborating the presence of CDH23 in the complex.

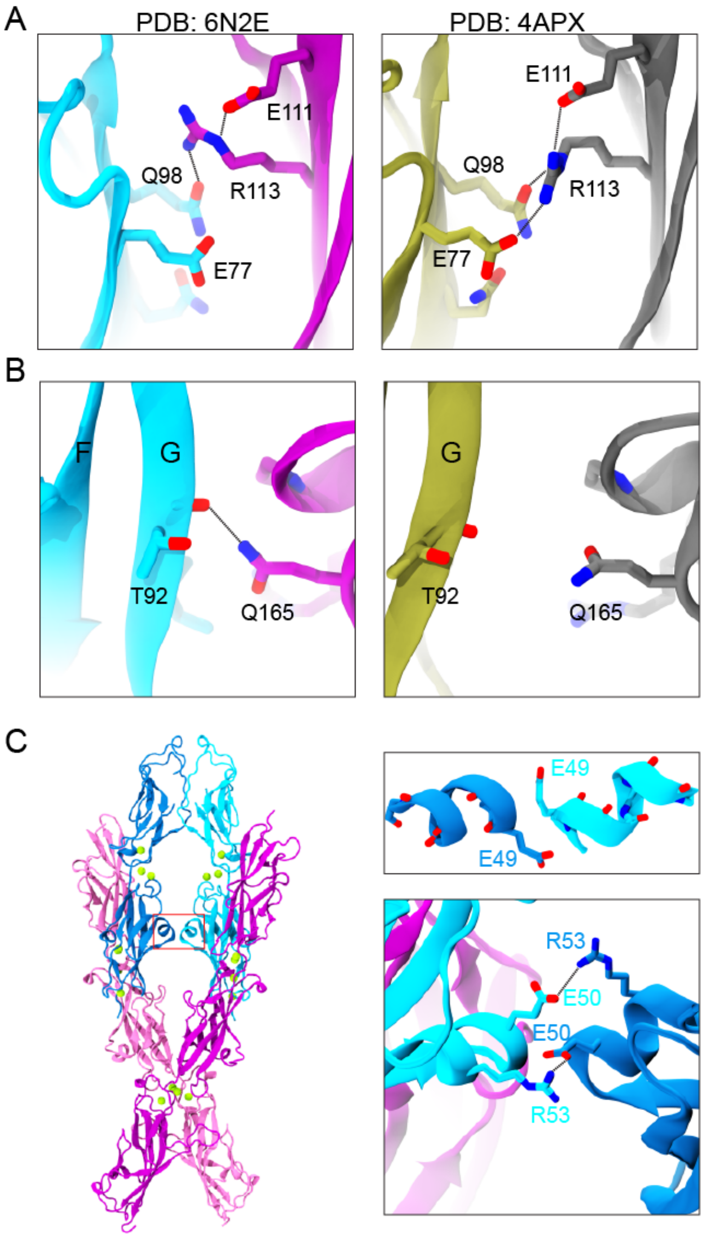


**Figure S11. Structural details of the CDH23-PCDH15 interface in the tetramer complex.** (**A**) Left panel shows details of the interface between CDH23 (chain C, cyan) and PCDH15 (chain D, purple) in the *hs* PCDH15 EC1-3 G16D/N369D/Q370N + *mm* CDH23 EC1-2 T15E structure (PDB: 6N2E). Residue p.R113 in PCDH15 does not interact with p.E77 in CDH23, as observed in a structure of the complex involving isoform N2 of PCDH15 (PDBs: 4XXW)[37]. Right panel shows the same detail for the *mm* PCDH15 EC1-2 (gray) + *mm* CDH23 EC1-2 (tan) complex (PDB: 4APX)[33]. Residue p.R113 is oriented towards the interface forming more favorable interactions with p.E77. (**B**) Left panel shows residue p.Q165 of PCDH15 making a favorable hydrogen bond with the backbone carbonyl of CDH23's p.T92 (β-strand G of EC1) in the *hs* PCDH15 EC1-3 G16D/N369D/Q370N + *mm* CDH23 EC1-2 T15E structure (PDB: 6N2E). The same interaction is not observed in the structure of the *mm* PCDH15 EC1-2 + *mm* CDH23 EC1-2 complex (PDB: 4APX)[33] due to an increased separation between the CDH23 N-terminus and the PCDH15 EC2 repeat. Colors as in A. (**C**) Ribbon representation of the *hs* PCDH15 EC1-3 G16D/N369D/Q370N (purple and mauve) + *mm* CDH23 EC1-2 T15E (cyan and blue) structure (PDB: 6N2E). Red box highlights contacts between CDH23 protomers at their $3_{10}$ helices in EC1 (between β-strands C and D). Insets show details of residue p.E49 from each helix capping the dipole of the opposite helix (top panel) and p.E50:p.R53 salt bridges "interlocking" the helices of the two CDH23 protomers (bottom panel).

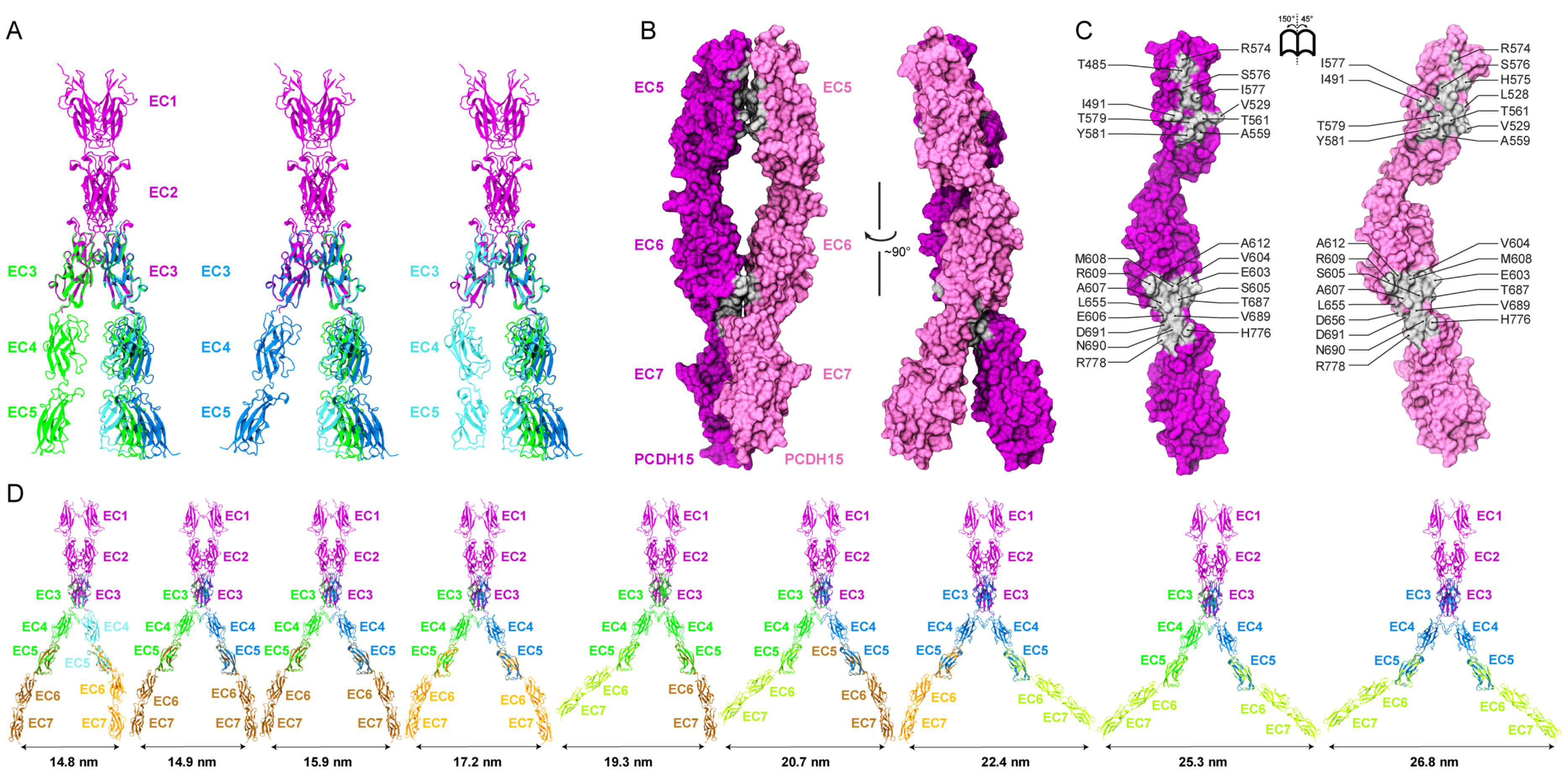


**Figure S12. Conformational diversity of PCDH15's ectodomain and the crystallographic interface in PCDH15 EC5-7.** (**A**) Overlap of different conformations of *hs* PCDH15 EC3-5 CD2-1 (green, blue, cyan for different protomers in the asymmetric unit) and the PCDH15 protomers from the heterotetrameric *hs* PCDH15 EC1-3 G16D/N369D/Q370N + *mm* CDH23 EC1-2 T15E structure. All possible conformations are compatible with the X-dimer arrangement. (**B**) Molecular surface representation of two *mm* PCDH15 EC5-7 I582T protomers (mauve and purple) in the asymmetric unit. Two perpendicular views are shown. (**C**) Interaction surface exposed with interfacing residues listed and shown in silver. (**D**) Using the crystallographic conformations of the PCDH15 EC3-5 and EC5-7 linkers (from our structures *hs* PCDH15 EC3-5 CD2-1 colored as in A, *mm* PCDH15 EC4-7 [brown], and *mm* PCDH15 EC5-7 I582T [orange and lime]), we generated all possible structural configurations (45 total) of the EC2-3 mediated PCDH15 EC1-7 homodimer. Here, we show nine representative configurations. The distance between the EC7 C-termini was measured and is indicated at the bottom of each configuration, some of which are compatible with *cis* (left) or *trans* (right) dimers (Fig. S6A-B).

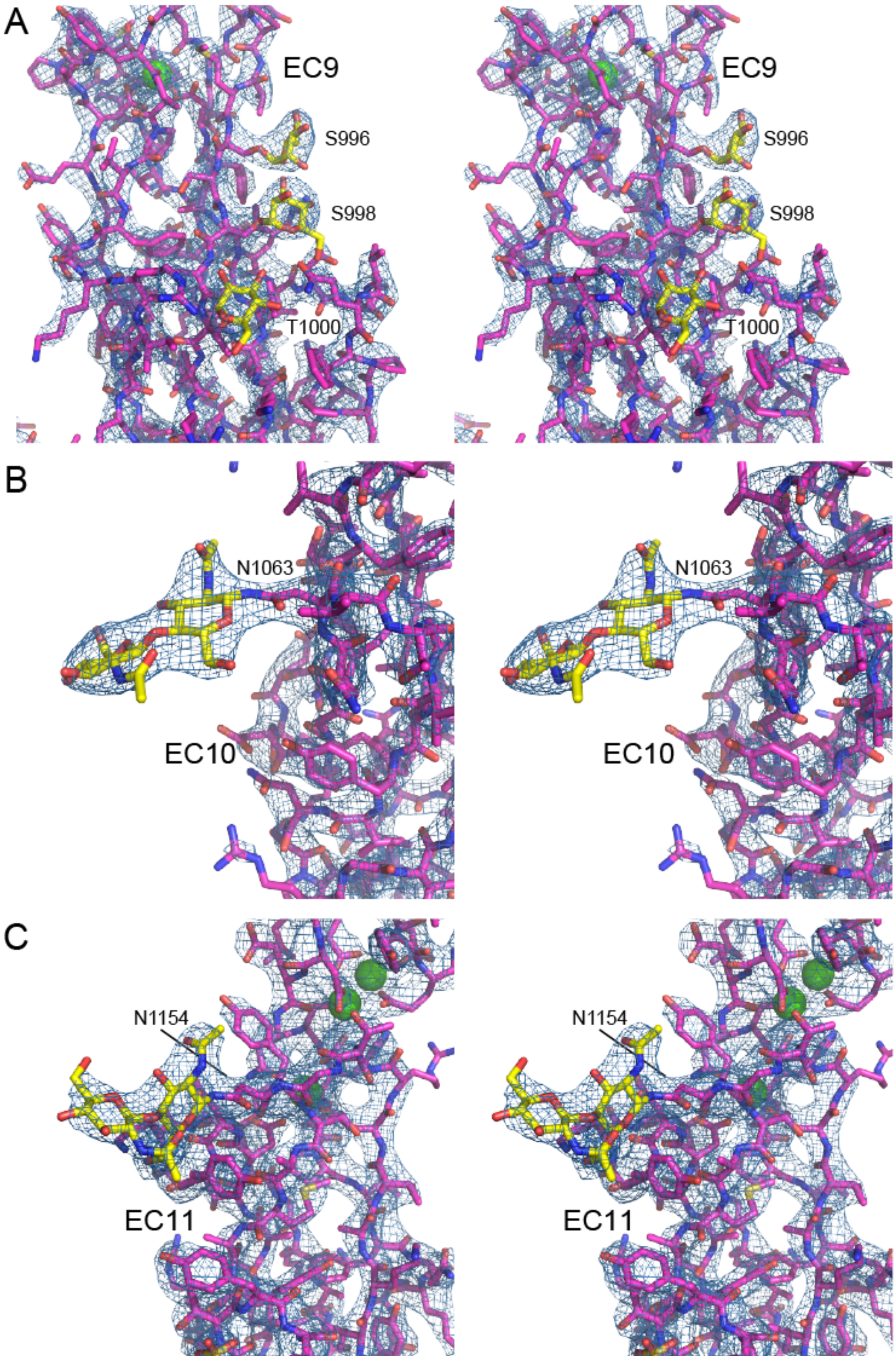


**Figure S13. PCDH15 glycosylation sites in EC9, EC10, and EC11.** Stereo view of the 2Fo-Fc electron density map (blue mesh; contoured at 2.0 σ) for the *mm* PCDH15 EC9-MAD12 structure (PDB: 6EET). (**A**) Detail of glycosylation sites on EC9. (**B**) Detail of glycosylation site on EC10. (**C**) Detail of glycosylation site on EC11.

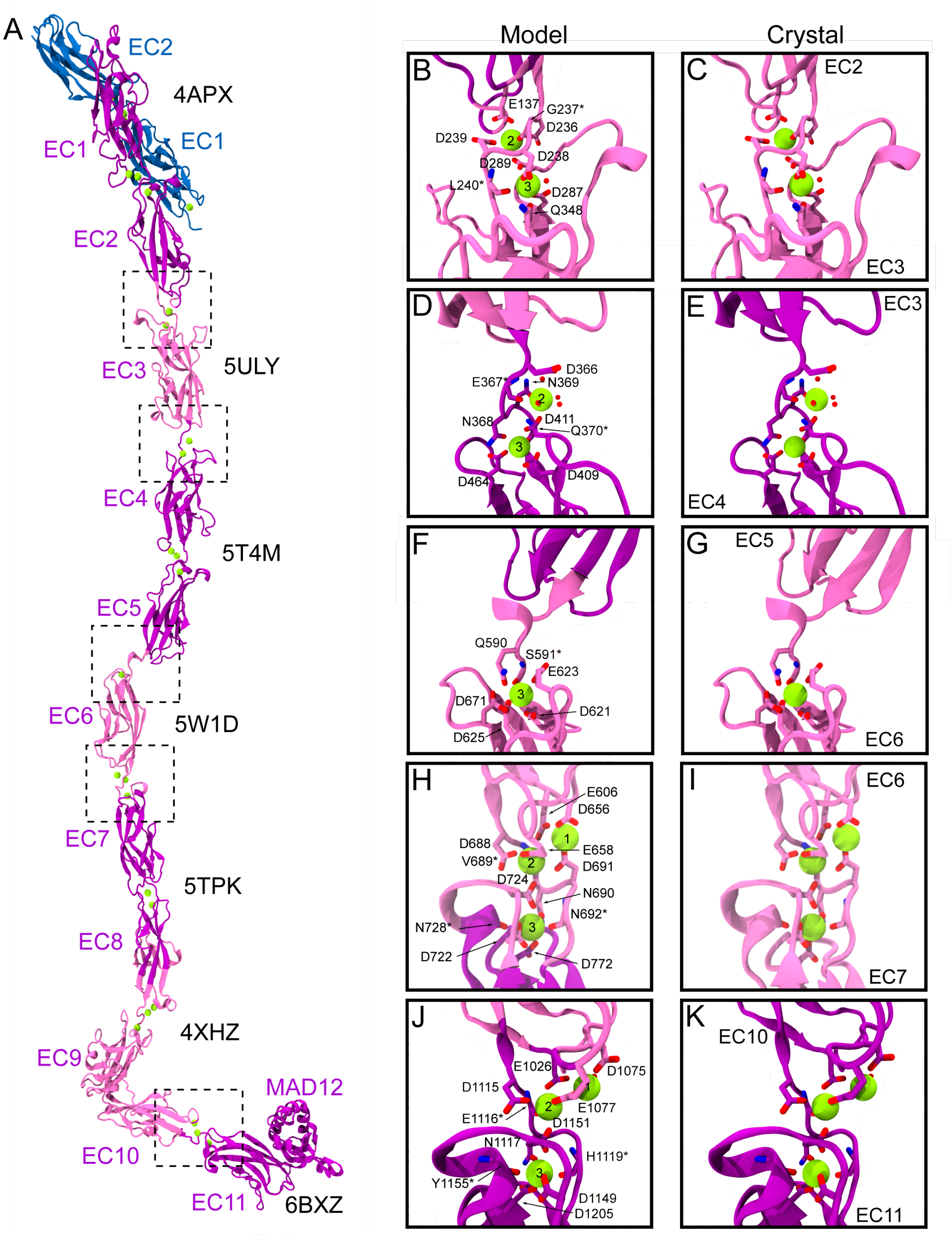


**Figure S14. Overall view and details of the complete *hs* PCDH15 ectodomain model.** (**A**) Ribbon representation of the entire *hs* PCDH15 EC1-MAD12 CD1-1 ectodomain (mauve and purple) bound to *hs* CDH23 EC1-2 (blue). The alternating colors (mauve and purple) highlight different crystal structures used to assemble the model (PDB codes indicated for each in black). Regions inside the dotted boxes highlight select fusion points between crystal structures. Crystallographic water molecules (red spheres), $Ca^{2+}$ ions (green spheres), and $Ca^{2+}$-coordinating residues at these points are highlighted and labeled in insets. For structures with multiple chains in the asymmetric unit the following chains were used: chain D of 5ULY; chain A of 5T4M; and chain C of 6BXZ. (**B-C**) Detail of the fused PCDH15 EC2-3 linker region in the model (PDBs: 4APX + 5ULY) and of the same linker region in a crystal structure (PDB: 5ULY). (**D-E**) Detail of the fused PCDH15 EC3-4 linker region in the model (PDBs: 5ULY + 5T4M) and of the same linker region in a crystal structure (PDB: 5T4M). (**F-G**) Detail of the fused PCDH15 EC5-6 linker region (PDBs: 5T4M + 5W1D) and of the same linker region in a crystal structure (PDB: 5W1D). (**H-I**) Detail of the fused PCDH15 EC6-7 linker region in the model (PDBs: 5W1D + 5TPK) and of the same linker region in a crystal structure (5W1D). (**J-K**) Detail of the fused PCDH15 EC10-11 linker region in the model (PDBS: 4XHZ + 6BXZ) and of the same linker region in a crystal structure (6BXZ).

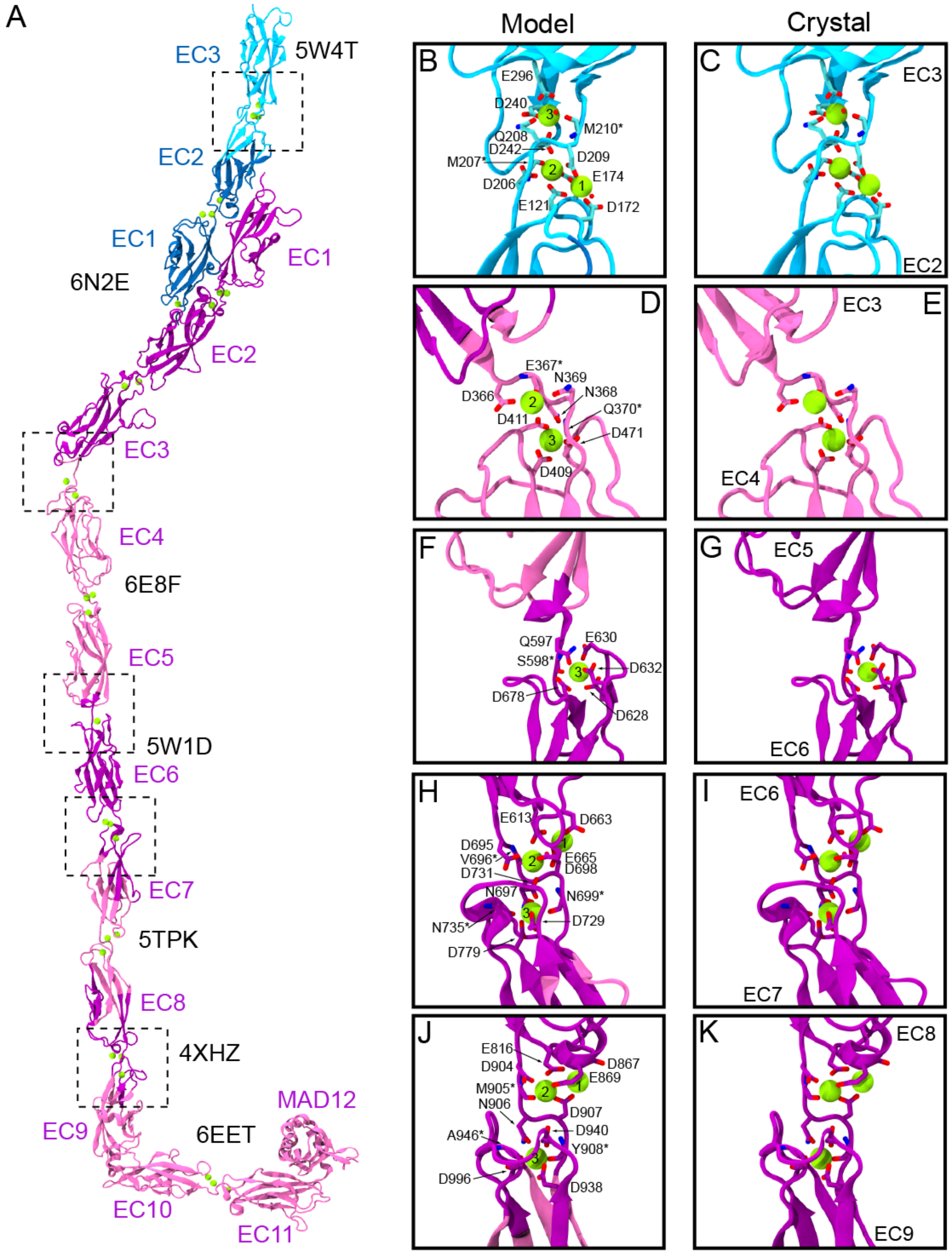


**Figure S15. Overall view and details of the complete *mm* PCDH15 ectodomain model.** (**A**) Ribbon representation of the entire *mm* PCDH15 EC1-MAD12 CD2-1 ectodomain (mauve and purple) bound to *mm* CDH23 EC1-3 (blue and cyan). The alternating colors (mauve and purple; blue and cyan) highlight different crystal structures used to assemble the model (PDB codes indicated for each in black). Regions inside the dotted boxes highlight select fusion points between crystal structures. Crystallographic water molecules (red spheres), $Ca^{2+}$ ions (green spheres), and $Ca^{2+}$-coordinating residues at these points are highlighted and labeled in insets. For structures with multiple chains in the asymmetric unit the following chains were used: chains B & C of 6N2E; chain A of 5W4T; and chain B of 6E8F. (**B**-**C**) Detail of the fused CDH23 EC2-3 linker region in the model (PDBs: 5W4T + 6N2E) and of the same linker region in a crystal structure (PDB: 5W4T), respectively. (**D**-**E**) Detail of the fused PCDH15 EC3-4 linker region in the model (PDBs: 6N2E + 6E8F) and of the same linker region in a crystal structure (PDB: 6E8F). (**F**-**G**) Detail of the fused PCDH15 EC5-6 linker region (PDBs: 6E8F + 5W1D) and of the same linker region in a crystal structure (PDB: 5W1D). (**H**-**I**) Detail of the fused EC6-7 linker region in the model (PDBs: 5W1D + 5TPK) and of the same linker region in a crystal structure (5W1D). (**J**-**K**) Detail of the fused EC8-9 linker region in the model (PDBs: 4XHZ + 6EET) and of the same linker region in a crystal structure (4XHZ).

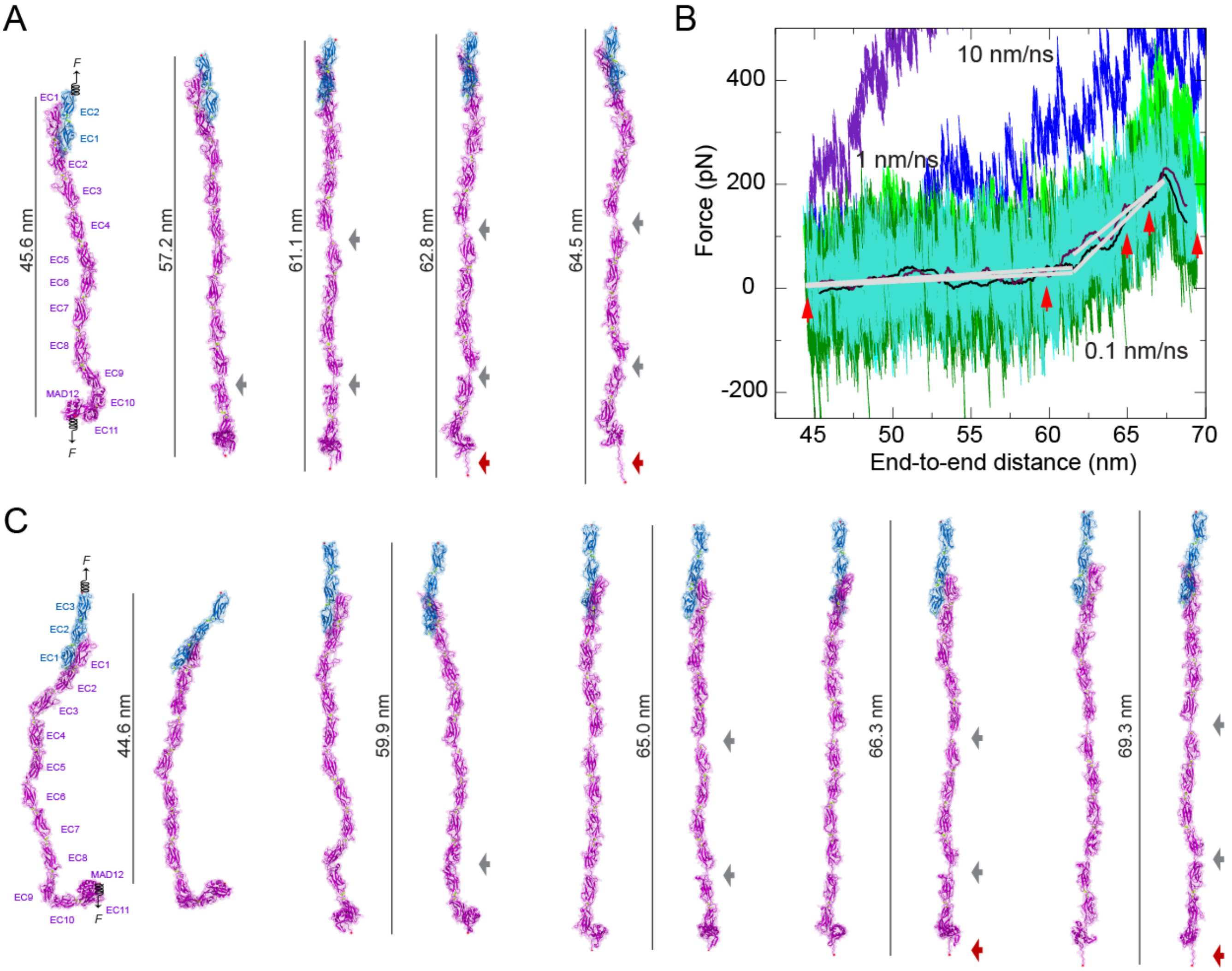


**Figure S16. Elasticity of the monomeric PCDH15 ectodomain.** (**A**) Snapshots of the monomeric *hs* PCDH15 EC1-MAD12 CD1-1 + CDH23 EC1-2 system during stretching simulation S1d (0.1 nm/ns, Table S8). Stretched C-terminal Cα atoms are shown as red spheres. Springs indicate position and direction of applied forces. Gray arrows highlight stretching of PCDH15 EC linkers (EC9-10 followed by EC5-6). Dark red arrow indicates unfolding of PCDH15 MAD12's C-terminal end. Views are rotated versions of snapshots shown in Fig. 5K (~90°). (**B**) Force versus end-to-end distance for constant velocity stretching of *mm* PCDH15 EC1-MAD12 + CDH23 EC1-3 at 10 nm/ns (S2b, purple and blue), 1 nm/ns (S2c, bright green and cyan), and 0.1 nm/ns (S2d, dark green and turquoise; 10-ns running averages shown in black and maroon; gray lines are fits used to determine elasticity of the complex). Red arrowheads indicate time-points for S2d illustrated in C. (**C**) Snapshots of the monomeric *mm* PCDH15 EC1-MAD12 CD1-1 + CDH23 EC1-3 system during stretching simulation S2d (0.1 nm/ns, Table S8). Interestingly, straightening of the bent monomeric PCDH15 conformation, before any unfolding happens, is soft and leads to a ~15.5 nm smooth extension.

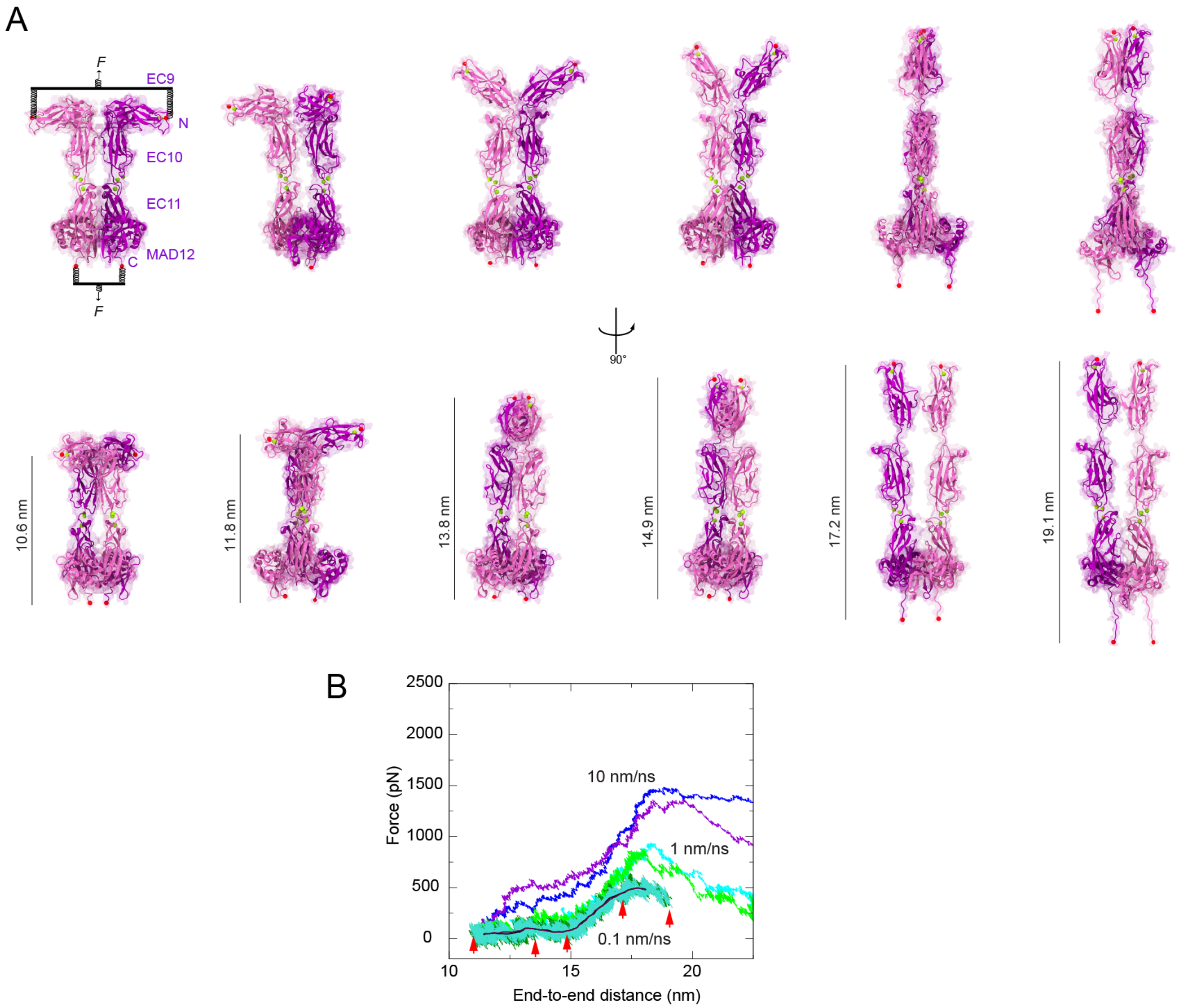


**Figure S17. Elasticity of *mm* PCDH15 EC9-MAD12.** (**A**) Snapshots showing two views (top and bottom rows) of the dimeric *mm* PCDH15 EC9-MAD12 system during stretching simulation S11d (0.1 nm/ns, Table S8). Stretched N- and C-terminal Cα atoms are shown as red spheres. Stretching was carried out by attaching two slabs to springs that were in turn attached to the terminal ends of each protein protomer. Slabs were moved in opposite directions at constant speed through individual springs. (**B**) Force applied to each of the slabs versus protein separation (see Methods) for stretching simulations of *mm* PCDH15 EC9-MAD12 (S11b-d). Traces are for constant velocity simulations at 10 nm/ns (purple and blue), 1 nm/ns (bright green and cyan), and 0.1 nm/ns (dark green and turquoise with 10-ns running averages in black and maroon). Red arrowheads indicate time-points for simulation S11d illustrated in A. The force profile shows a clear deep at 14.9 nm, suggesting an alternate stable state with semi-extended EC9 repeats.

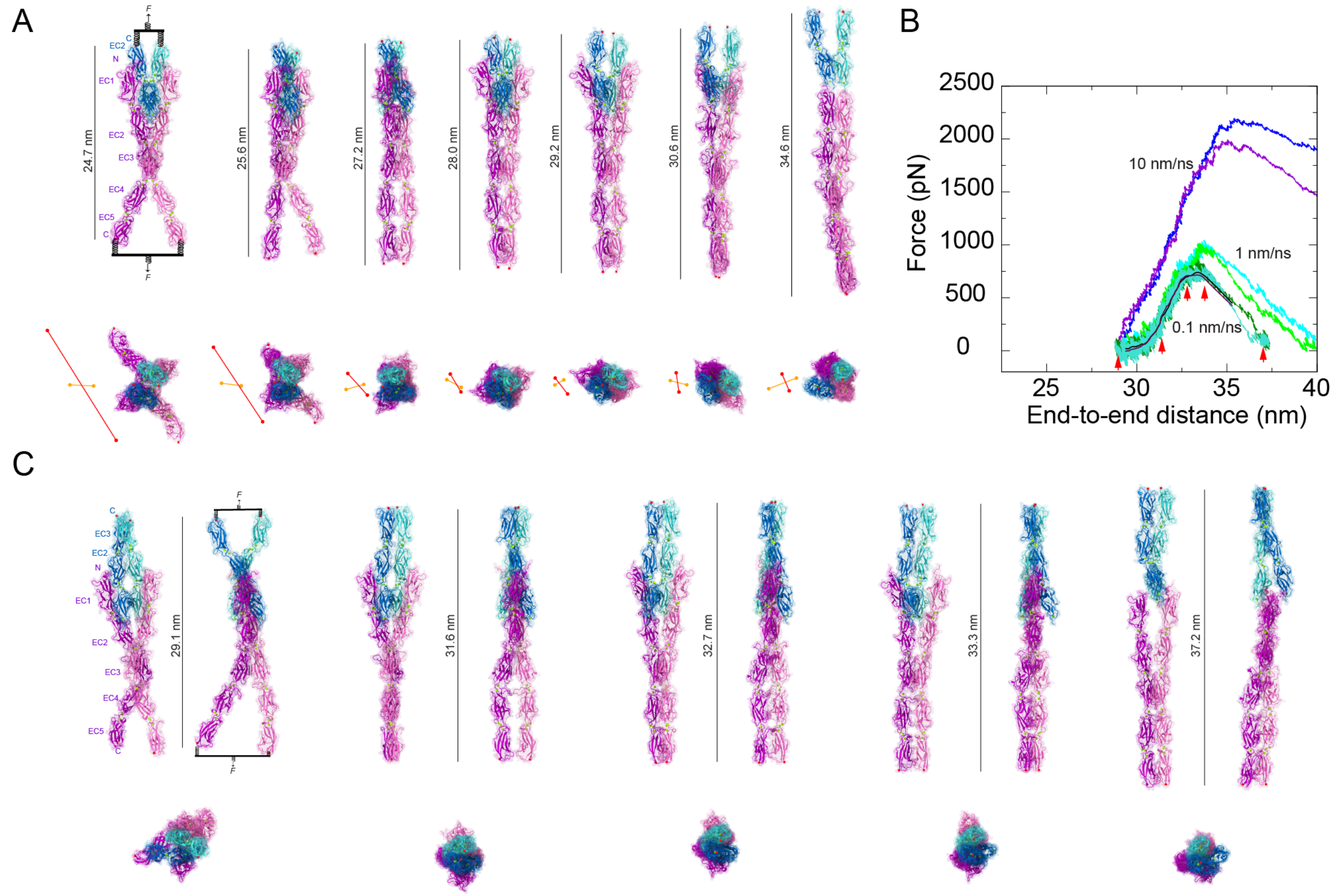


**Figure S18. Constant-velocity SMD simulations of the short PCDH15 + CDH23 heterotetrameric bond.** (**A**) Snapshots of the heterotetrameric *hs* (PCDH15 EC1-5)$_2$ and (CDH23 EC1-2)$_2$ system during stretching simulation S3d (0.1 nm/ns, Table S8). Side and top views are shown as in Fig. 6A. Position of C-terminal Cα atoms for PCDH15 and CDH23 protomers are shown in red and orange, respectively. Lines between these positions illustrate closing of parallel protomers as well as relative angle and rotations of PCDH15 with respect to CDH23 protomers observed throughout the trajectory (**B**) Force applied to each of the slabs versus protein separation (see Methods) for stretching simulations of *mm* (PCDH15 EC1-5)$_2$ and (CDH23 EC1-3)$_2$ (S7b-d). Traces are for constant velocity simulations at 10 nm/ns (purple and blue), 1 nm/ns (bright green and cyan), and 0.1 nm/ns (dark green and turquoise with 10-ns running averages in black and maroon). Red arrowheads indicate time-points for simulation S7d illustrated in C. (**C**) Snapshots of the heterotetrameric *mm* (PCDH15 EC1-5)$_2$ and (CDH23 EC1-3)$_2$ system during stretching simulation in which slabs where connect to stretching springs and moved in opposite directions (S7d 0.1 nm/ns, Table S8). Side and top views are shown as in A.

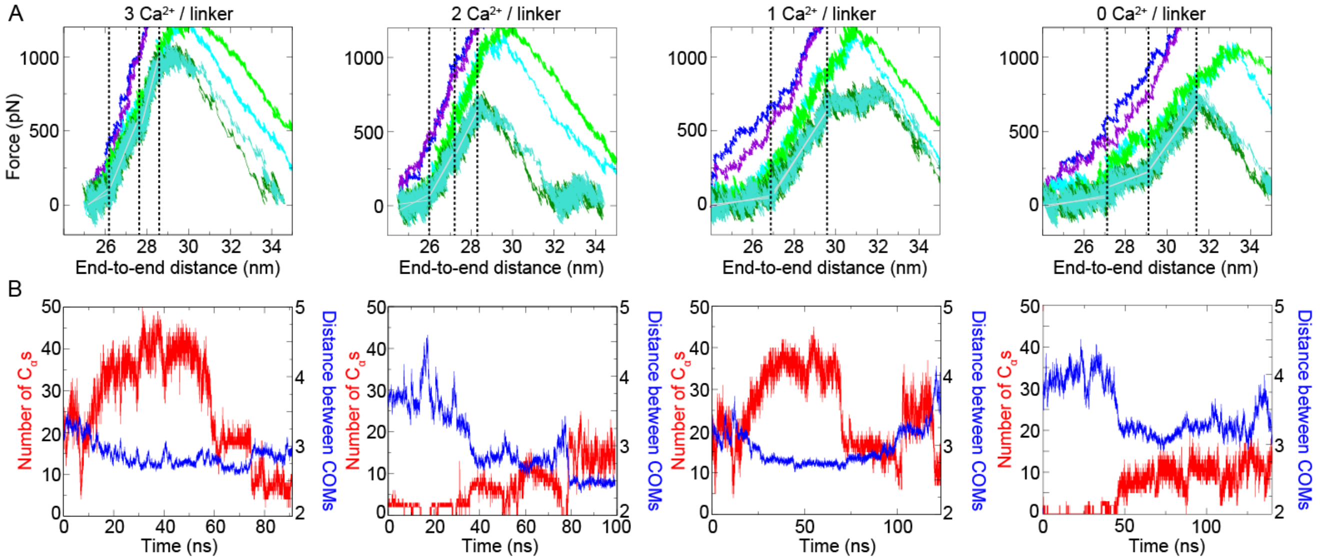


**Figure S19. Ca$^{2+}$-dependent elasticity and dynamics of the short PCDH15 + CDH23 heterotetrameric bond.** (**A**) Force versus end-to-end distance profile in simulations of the *hs* (PCDH15 EC1-5)$_2$ and (CDH23 EC1-2)$_2$ *trans* tetramer with different number of Ca$^{2+}$ ions at linker regions when stretched at 10 nm/ns (purple and blue), 1 nm/ns (bright green and cyan), 0.1 nm/ns (dark green and turquoise). Different phases of the curves were fitted by lines (gray), the slopes of which correspond to stiffness. The soft and stiff phases were fitted separately for 3 Ca$^{2+}$ (92.9 mN/m, 343.5 mN/m, 560.4 mN/m), 2 Ca$^{2+}$ (46.2 mN/m, 250.6 mN/m, 278.2 mN/m), 1 Ca$^{2+}$ (20.5 mN/m, 216.4 mN/m), and 0 Ca$^{2+}$ (21.7 mN/m, 52.3 mN/m, 197.9). Black vertical dashed lines mark boundary of different phases. The complex becomes softer with decreasing number of Ca$^{2+}$ ions at the linker regions. (**B**) Dynamics of CDH23 molecules as a function of time for simulations S3d to S6d. The interaction between CDH23 protomers is defined by residues from the two protomers that are within 4 Å of each other. The total number of such Cα atoms is shown in red. The distance between centers of mass of the two EC1 domains of CDH23 is shown in blue. The two CDH23 protomers come in close contact ("squeezed") during the SMD simulation with Ca$^{2+}$ ions.

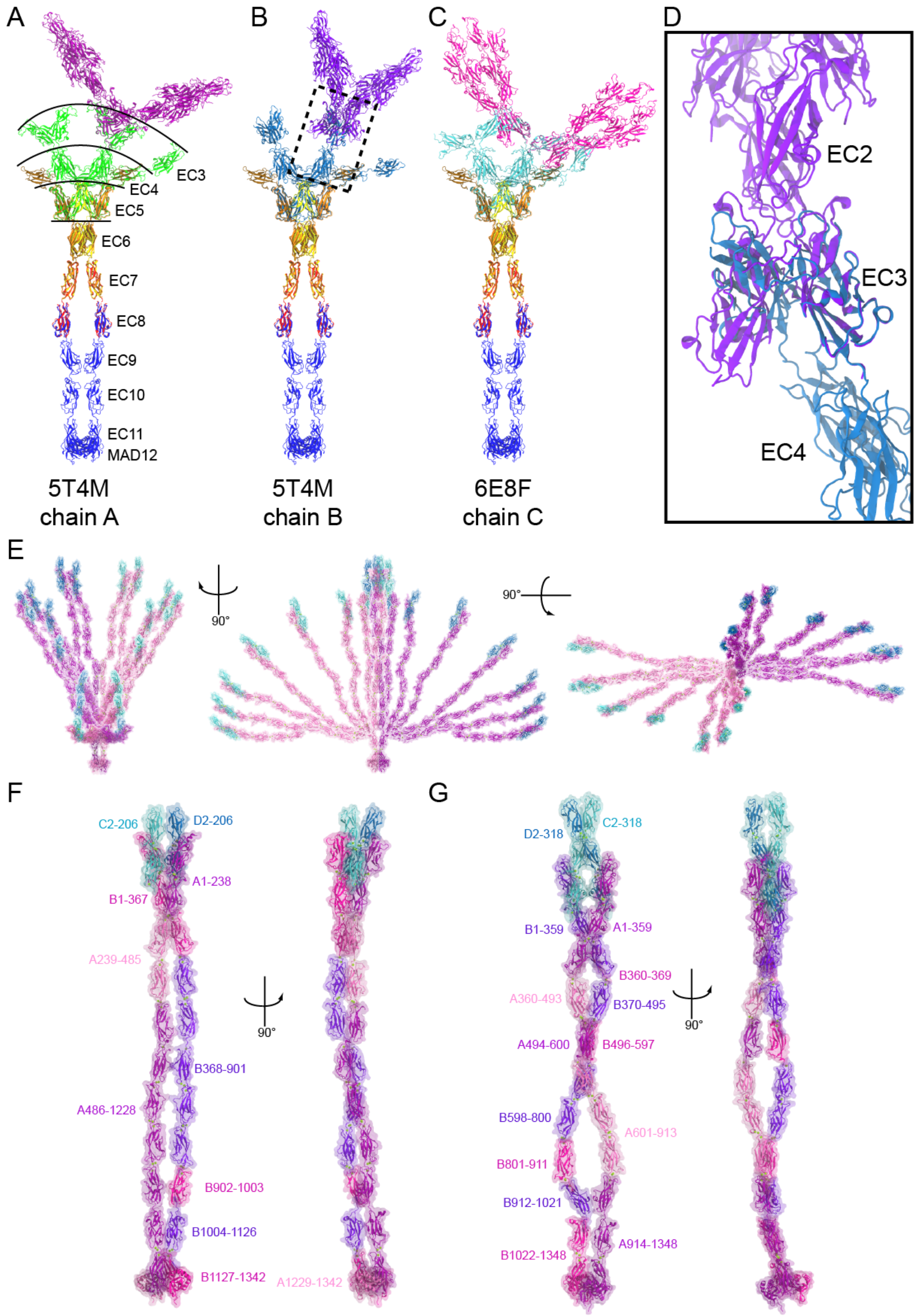


**Figure S20. Assembly of PCDH15 parallel dimers.** (**A**) Use of crystallographic and cryo-EM structures to assemble the PCDH15 parallel dimer. The homodimeric cryo-EM structure of *mm* PCDH15 EC8-MAD12 (blue; PDB: 6C13)[39] and the crystal structure of *mm* PCDH15 EC7-8 V875A (red) were superimposed using EC8. Chain A (orange) and chain B (yellow) of *mm* PCDH15 EC5-7 I582T and *mm* PCDH15 EC4-7 (brown) were superimposed using EC7 of *mm* PCDH15 EC7-8 V875A. Chain A of *hs* PCDH15 EC3-5 (green; PDB: 5T4M)[44] was overlapped with EC4 or EC5. The *hs* PCDH15 EC1-3 G16D/N369D/Q370N + *mm* CDH23 EC1-2 T15E structure (purple) was overlapped using EC3 (only two shown for clarity). The PCDH15 EC1-MAD12 parallel dimer constructed solely from the crystal and cryo-EM structures mentioned above was not compatible with the EC2-3-mediated X-dimer. (**B**) Similar to A but using chain B of *hs* PCDH15 EC3-5 (blue; PDB: 5T4M)[44]. The overlap of EC3 of *hs* PCDH15 EC1-3 G16D/N369D/Q370N + *mm* CDH23 EC1-2 T15E (purple) came close to an X-dimer conformation. (**C**) Similar to panel A but using chain C of *hs* PCDH15 EC3-5 CD2-1 (cyan). The overlap of EC3 of *hs* PCDH15 EC1-3 G16D/N369D/Q370N + *mm* CDH23 EC1-2 T15E (purple) was not compatible with an X-dimer conformation. (**D**) Detail from panel B showing the closest the structures come to creating the PCDH15 X-dimer mediated by EC2-3. The overlap of EC3 of *hs* PCDH15 EC3-5 chain B (blue: 5T4M) and *hs* PCDH15 EC1-3 G16D/N369D/Q370N + *mm* CDH23 EC1-2 T15E (purple) is on the right side. The other EC3 (left side) of *hs* PCDH15 EC1-3 G16D/N369D/Q370N + *mm* CDH23 EC1-2 T15E did not align well to the closest available EC3 from a PCDH15 EC3-5 structure. (**E**) Multiple views of the *hs* PCDH15 EC1-MAD12 CD1-1 + CDH23 EC1-2 system during stretching simulation S1d (0.1 nm/ns, Table S8, 0 – 70 ns) overlapped on the crystal structure of *mm* PCDH15 EC9-MAD12. Snapshots were taken every 10 ns (one at 35 ns). None of the depicted states is compatible with the EC2-3 X-dimer. (**F**) Details of the structures and simulation trajectory coordinates used to assemble the human $(\text{PCDH15 EC1-11+MAD12})_2$ and $(\text{CDH23 EC1-2})_2$ heterotetramer. Residue ranges for pieces of PCDH15 (A,B) and CDH23 (C,D) chains from different structures and simulation snapshots are labeled (see Methods). (**G**) Details of the structures and simulation trajectory coordinates used to assemble the mouse $(\text{PCDH15 EC1-11+MAD12})_2$ and $(\text{CDH23 EC1-3})_2$ heterotetramer. Residue ranges for pieces of PCDH15 (A,B) and CDH23 (C,D) chains from different structures and simulation snapshots are labeled (see Methods).

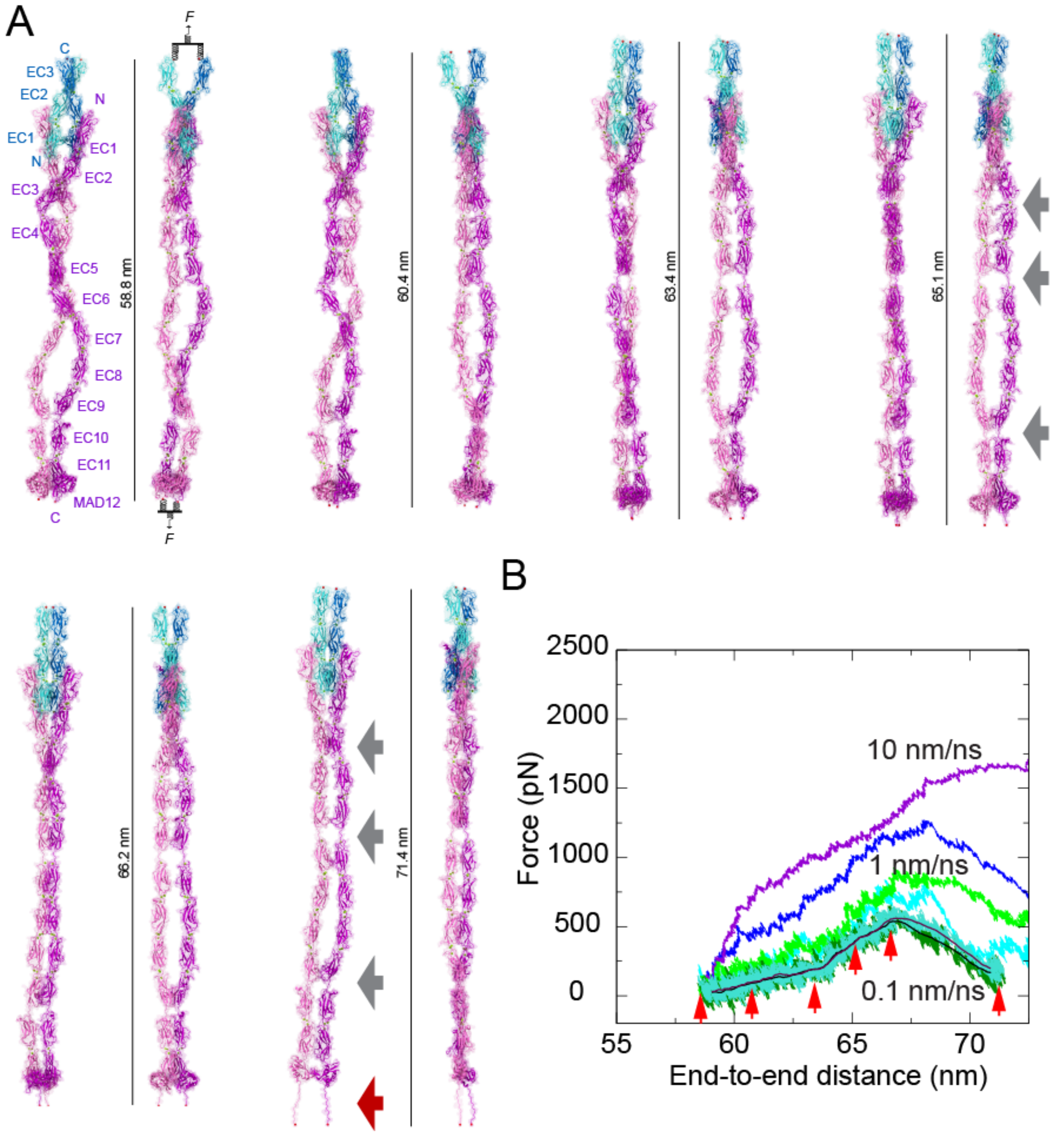


**Figure S21. Mechanics of the mouse PCDH15 ectodomain as part of an heterotetrameric complex with CDH23.** (**A**) Snapshots of the heterotetrameric *mm* (PCDH15 EC1-MAD12)$_2$ + (CDH23 EC1-3)$_2$ system during stretching simulation S9d (0.1 nm/ns). Stretched C-terminal Cα atoms are shown as red spheres. Stretching was carried out by attaching two slabs to springs that were in turn attached to the terminal ends of each protein protomer. Slabs were moved in opposite directions at constant speed through individual springs. Gray arrows highlight location of flexible PCDH15 EC linkers (EC9-10, EC5-6, and EC3-4). Dark red arrow indicates unfolding of PCDH15 MAD12's C-terminal end. (**B**) Force applied to slabs versus protein separation (see Methods and Table S8) for constant velocity stretching simulations of the *mm* (PCDH15 EC1-MAD12)$_2$ + (CDH23 EC1-3)$_2$ system at 10 nm/ns (S9b, purple and blue), 1 nm/ns (S9c, bright green and cyan), and 0.1 nm/ns (S9d, dark green and turquoise with 10-ns running averages in black and maroon). Red arrowheads indicate time-points for simulation S9d illustrated in A.

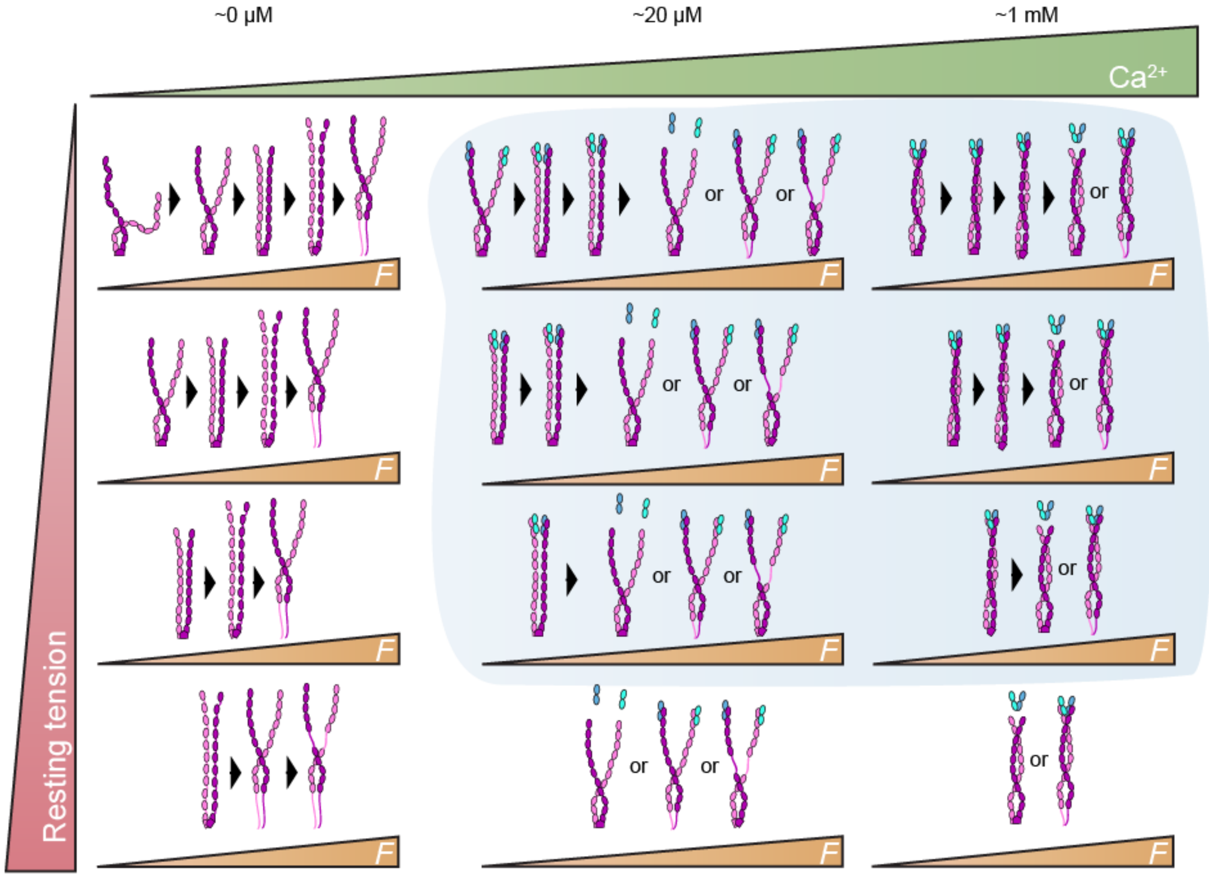


**Figure S22. Elastic response of PCDH15 in the presence of resting tension.** Tip links experience a constant resting tension that varies along the cochlea from ~5 to 50 pN[79]. This resting tension will pre-stretch the PCDH15 ectodomain, thus conditioning its elastic response to an external stimulus (*F* in the figure).

**Table S1.** List of species and PCDH15 accession numbers used in multiple sequence alignment analysis

| Species | Abbreviation | NCBI Accession number |
|---|---|---|
| *Homo sapiens* CD1-1 | *Hs1* | NP_001136235.1 |
| *Homo sapiens* CD2-1 | *Hs2* | NP_001136241.1 |
| *Tursiops truncatus* | *Tt* | XP_019792025.1 |
| *Sus scrofa* | *Ss* | XP_020929194.1 |
| *Canis lupus familiaris* | *Clf* | XP_022266127.1 |
| *Mus musculus* | *Mm* | NP_075604.2 |
| *Desmodus rotundus* | *Dr2* | XP_024416857.1 |
| *Gallus gallus* | *Gg* | XP_015143564.1 |
| *Columba livia* | *Cl* | XP_021147002.1 |
| *Lonchura striata domestica* | *Lsd* | XP_021391314.1 |
| *Corvus brachyrhynchos* | *Cb* | XP_017589248.1 |
| *Pogona vitticeps* | *Pv* | XP_020662251.1 |
| *Anolis carolinensis* | *Ac* | XP_016851436.1 |
| *Crocodylus porosus* | *Cp* | XP_019394135.1 |
| *Danio rerio* | *Dr* | NP_001012500.1 |
| *Clupea harengus* | *Ch* | XP_012675462.1 |
| *Astyanax mexicanus* | *Am* | XP_022525074.1 |
| *Callorhinchus milii* | *Cm* | XP_007897895.1 |

**Table S2.** X-ray crystallography experiments details section (conditions for crystal growth and cryo protection)

| Construct | Crystallization conditions | Cryo | P + B ($\mu$l) |
|---|---|---|---|
| *Mm* PCDH15 EC1-2BAP | 0.02 M $CaCl_2$, 0.1M Sodium Acetate, 30% MPD | | 0.6 + 0.6 |
| *Hs* PCDH15 EC1-3 G16D/N369D/Q370N | 0.1 M HEPES pH 7.7, 66% MPD, 4% Glycerol | | 1.0 + 0.5 |
| *Hs* PCDH15 EC1-3 G16D/N369D/Q370N + *mm* CDH23 EC1-2 T15E | 0.1 M Imidazole pH 6.8, 46% MPD | | 0.6 + 0.6 |
| *Hs* PCDH15 EC2-3 | 0.1 M HEPES pH 7.5, 0.1 M KCl, 15% PEG 6000 | | 0.6 + 0.6 |
| *Hs* PCDH15 EC2-3 V250N | 0.1 M MES pH 5.9, 0.15 M $MgCl_2$, 32% PEG 400 | | 0.6 + 0.6 |
| *Hs* PCDH15 EC3-5 CD2-1 | 40% MPD, 0.2 M Lithium Chloride, 0.01 M ATP Disodium | | 0.6 + 0.6 |
| *Mm* PCDH15 EC4-7 | 0.1 M Tris-HCl pH 8.5, 2.0 M Magnesium Acetate | 23% PEG 400 | 0.6 + 0.6 |
| *Mm* PCDH15 EC5-7 I582T | 0.2 M Sodium Formate, 40% MPD | | 0.6 + 0.6 |
| *Mm* PCDH15 EC6-7 | 30% PEG 1500 | 10% PEG 400 | 0.6 + 0.6 |
| *Mm* PCDH15 EC7-8 V875A | 10% Ammonium Chloride, 10% PEG3350 | 25% Glycerol | 1.0 + 0.5 |
| *Mm* PCDH15 EC9-MAD12 | 0.1 M Sodium Salt, 30% MPD, 5% PEG 4000 | | 0.6 + 0.6 |

**Table S3.** Expression, refolding, and purification conditions for bacterially expressed protein fragments

| Fragment | Cells | Media | T °C | IPTG μM | Refolding | SEC |
|---|---|---|---|---|---|---|
| *Mm* PCDH15 EC1-2BAP | RIPL | TB | 30 | 200 | RS | SB1 pH 7.5 |
| *Hs* PCDH15 EC1-3 WT and G16D/N369D/Q370N | RIPL | TB | 30 | 200 | RS | SB1 pH 8.0 |
| *Hs* PCDH15 EC1-4 WT and L306N/V307N | RIPL | TB | 30 | 200 | RS | SB1 pH 8.0 |
| *Mm* CDH23 EC1-2 T15E | RIPL | LB | 30 | 200 | R1 | SB1 pH 8.0 |
| *Hs* PCDH15 EC2-3 WT & mutants | RIPL | LB | 30 | 200 | R1 | SB1 pH 8.0 |
| *Hs* PCDH15 EC3-5 CD2-1 | RIPL | LB | 30 | 200 | R2 | SB2 |
| *Mm* PCDH15 EC4-7 | BL21 | TB | 30 | 1000 | R3 | SB3 |
| *Mm* PCDH15 EC5-7 I582T | RIPL | TB | 30 | 1000 | R4 | SB4 |
| *Mm* PCDH15 EC6-7 | Rosetta | TB | 30 | 1000 | R5 | SB3 |
| *Mm* PCDH15 EC7-8 V875A | RIPL | LB | 37 | 1000 | R5 | SB4 |
| *Mm* PCDH15 EC9-MAD12 | Rosetta | LB | 30 | 1000 | R7 | SB5 |

RIPL – *E. coli* BL21 CodonPlus(DE3)-RIPL cells.
BL21 – *E. coli* BL21 cells.
Rosetta – *E. coli* BL21 Rosetta(DE3) cells.

RS – Five-step dialysis. Two 24-h dialysis steps against D buffer (20 mM Tris HCl, pH 8.0, 10 mM $CaCl_2$) plus 3 M and 2 M GuHCl, respectively. Last three steps consisted of 12-h dialyses against D buffer with decreasing GuHCl concentration (1, 0.5, and 0 M) plus 400 mM L-Arg and 375 μM GSSG.
R1 – Overnight dialysis in 20 mM Tris HCl, pH 8.0, 150 mM KCl, 50 mM NaCl, 2 mM $CaCl_2$, and 400 mM L-Arg (+10% glycerol for *hs* PCDH15 EC2-3 mutants).
R2 – Overnight dialysis in 20 mM Tris HCl, pH 8.0, 10 mM $CaCl_2$, 400 mM L-Arg, and 1 mM GSSG.
R3 – Overnight dialysis in 20 mM Tris HCl, pH 8.0, 150 mM KCl, 5 mM $CaCl_2$, 400 mM L-Arg, and 2 mM DTT.
R4 – Overnight dialysis in 20 mM Tris HCl, pH 8.0, 150 mM KCl, 50 mM NaCl, 2 mM $CaCl_2$, 400 mM L-Arg, and 2 mM DTT.
R5 – Overnight dialysis in 20 mM Tris HCl, pH 5.2, 150 mM KCl, 50 mM NaCl, 2 mM $CaCl_2$,400 mM L-Arg, and 2 mM DTT.
R6 – Overnight dialysis in 20 mM Tris HCl, pH 8.0, 150 mM KCl, 5 mM $CaCl_2$, , and 10% glycerol.
R7 – Drop-by-drop dilution[40] in 20 mM Tris HCl, pH 8.0, 150 mM KCl, 5 mM $CaCl_2$, 400 mM L-Arg, 1 mM TCEP-HCl and 10% glycerol.

SB1 – 20 mM Tris HCl, 150 mM KCl, 50 mM NaCl, and 2 mM $CaCl_2$.
SB2 – 20 mM Tris HCl pH 8.0, 150 mM KCl, 50 mM NaCl, and 5 mM $CaCl_2$ (double purified).
SB3 – 20 mM Tris HCl pH 8.0, 150 mM NaCl, 50 mM KCl, and 2 mM $CaCl_2$.
SB4 – 20 mM Tris HCl pH 8.0, 150 mM KCl, and 5 mM $CaCl_2$.
SB5 – 20 mM Tris HCl pH 8.0, 150 mM KCl, and 5 mM $CaCl_2$, 1 mM TCEP.

**Table S4.** Interface areas of PCDH15-CDH23 “handshakes” from various structures as computed by PISA.

| PDB | **Chains** | **Area (Å$^2$)** |
|---|---|---|
| 4APX | A/B | 907.0 |
| 4AQ8 | B/D | 1160.2 |
| 4AQ8 | A/C | 1070.2 |
| 4AQA | A/B | 958.4 |
| 4AQE | A/B | 894.6 |
| 4XXW | A/D | 1143.7 |
| 4XXW | B/C | 1061.0 |
| 6N2E | A/D | 1154.8 |
| 6N2E | B/C | 1209.6 |

**Table S5.** SAXS data collection and scattering-derived parameters.

| **Data collection parameters** | ***Mm* PCDH15 EC9-MAD12** |
|---|---|
| Instrument | SIBYLS beamline |
| Beam geometry | Point Focus |
| Wavelength (Å) | 1.03 |
| *q* range ($Å^{-1}$) | 0.009198 – 0.5925 |
| Exposure time (min) | 40 |
| Frame slicing (s) | 3 |
| Concentration (mg/mL) | 4.2 |
| Temperature (K) | 293.15 |
| **Structural parameters** | |
| $R_g$ (Å) [from P(*r*)] | 47.28 ± 0.24 |
| $R_g$ (Å) [from Guinier] | 46.73 ± 2.94 |
| $R_g$ (Å) [from 6EET]* | 42.01 |
| $D_{max}$ (Å) [from GNOM] | 151 ± 10 |
| **Molecular mass determination** | |
| Molecular mass** | 98.12 |
| Calculated mass from sequence (monomer) | 51.9 |
| Discrepancy (%) | 5.62 |
| Oligomeric state | Dimer |
| **Software employed** | |
| Primary data reduction | Beamline software |
| Data processing | PRIMUS, GNOM, SREFLEX |
| Computation of model intensities | FoXS |

* Estimated using VMD[126].

** Molecular mass estimated using the SAXS MoW2 server with the method described in [120].

**Table S6.** Structures used to build *hs* PCDH15 EC1-MAD12 CD1-1 + *hs* CDH23 EC1-2 and *mm* PCDH15 EC1-MAD12 CD2-1 + *mm* CDH23 EC1-3 models.

| | | | |
|---|---|---|---|
| | | ***Hs* PCDH15 EC1-MAD12 CD1-1 model** | |
| **EC** | **PDB ID (chain)** | **Residues** | **Mutations** |
| EC1-2 | 4APX (B) | 1-136, 138-200, 203-233 | L66M, V100I, V107I, E175D, V193I, Y208F |
| EC2-3 | 5ULY (D) | 137, 201-202, 234-365 | - |
| EC3-5 | 5T4M (A) | 366-584 | - |
| EC4-7 | 5W1D (A) | 585-694, 722-727 | E395D, T401S, P435Q, L459S, V473I, R477Q, S502R, S524T, G531A, K532Q, V536I, V544I, S545T, L552M, Q556R, S565A, H575N, V604I, I610V, I613V, P626S, S650T, V667I, R719K, A760V, H763Y, Y785A |
| EC7-8 | 5TPK (A) | 695-721, 728-805, 820-834, 873-894 | R719K, A760V, H763Y, Y785A, V806L, S816T, F817I, L843F, A875V |
| EC8-10 | 4XHZ (A) | 806-819, 835-872, 895-1025, 1027-1076, 1078-1113 | - |
| EC10-MAD12 | 6BXZ (C) | 1026, 1077, 1114-1342 | A1029T, L1036V, K1058T, A1072G, D1118N, A1140T, A1157V, V1300I |
| | | ***Hs* CDH23 EC1-2 model** | |
| EC1-2 | 4APX (A) | | R35Q, P153Q, Q168R, V174T |
| | | ***Mm* PCDH15 EC1-MAD12 CD2-1 model*** | |
| EC1-3 | 6N2E (B) | 5-265, 270-364 | D16G, M66L, I100V, I107V, D175E, I193V, F208Y, I278L, S327T, G359S |
| EC3-5 | 6E8F (B) | 266-269, 365-585(+7) | D395E, S401T, S411G(+7), Q435P(+7), S459L(+7), I473V(+7), Q477R(+7), R502S(+7), T524S(+7), A531G(+7), Q532K(+7), I536V(+7), I544V(+7), T545S(+7), M557L(+7), R556Q(+7), A565S(+7), N575H(+7), I582T(+7) |
| EC4-7 | 5W1D (A) | 585-695(+7), 721-741(+7), 764-783(+7) | - |
| EC7-8 | 5TPK (A) | 703-727(+7), 749-772(+7), 791-809(+7), 823-837(+7), 879-897(+7) | - |
| EC8-10 | 4XHZ (A) | 810-822(+7), 838-878(+7), 898-909(+7), 936-949(+7), 995-1002(+7) | L813V(+7), F850L(+7), L944M(+7) |
| EC9-MAD12 | 6EET (A) | 910-935(+7), 950-994(+7), 1003-1348(+7) | - |
| | | ***Mm* CDH23 EC1-3 model**** | |
| EC1-2 | 6N2E (C) | 3-116, 125-164, 182-199 | E16T, S128P |
| EC1-3 | 5W4T (A) | 114-121(+3), 162-178(+3), 197-315(+3) | I114V(+3), Q115R(+3), A167E(+3), T172V(+3), I174Q(+3), T200I(+3), I204M(+3), T212I(+3), M221Y(+3), D223H(+3), A224S(+3), Y228T(+3), E229T(+3), K232V(+3), R234T(+3), I236V(+3), L240K(+3), I251V(+3), M258I(+3), S267A(+3), V270L(+3), S271N(+3), Q273L(+3), S283H(+3), I287L(+3), A291G(+3), S302D(+3), T310N(+3), L313V(+3) |

* Residue numbering after exon 12a insertion is indicated with (+7).

** Residue numbering of the fish protein is shifted by 3 residues indicated with (+3).

**Table S7.** List of deafness-causing missense mutation sites and in-frame deletions on PCDH15.

| Citation | Mutation in *hs* | EC number | Note | Diagnosis |
|---|---|---|---|---|
| Miyagawa et al., 2013 | G79R | EC1 | o | Non-syndromic |
| Geng et al., 2013 | I108N | EC1 | h | USH1F |
| Ahmed et al., 2003 | R113G | EC1 | h | DFNB23 |
| Aller et al., 2010 | R113Q | EC1 | h | Likely USH1F |
| Ahmed et al., 2008 | D157G | EC2 | **D**xD | USH1F |
| Ahmed et al., 2003 | G241D | EC3 | i | DFNB23 |
| Miyagawa et al., 2013 | R257H | EC3 | o | Non-syndromic |
| Abdi et al., 2016 | E272_Q509del | EC3-5 | | USH1F |
| Miyagawa et al., 2013 | P294L | EC3 | o | Non-syndromic |
| Yang et al., 2013 | L408P | EC4 | o | Non-syndromic |
| Grossman et al., 2010 | D414A | EC4 | o | Non-syndromic |
| Doucette et al., 2009 | V507D | EC5 | i | DFNB23 |
| Zhan et al., 2015 | V767del | EC7 | | DFNB23 |
| Alagramam et al., 2011 | G936_K982del | EC9 | | USH1F |
| Miyagawa et al., 2013 | R941C | EC9 | o | Non-syndromic |
| Chen et al., 2015 | D989G | EC9 | x**D**x | DFNB23 |
| Schrauwen et al., 2018 | R1013H | EC10 | o | DFNB23 |
| Miyagawa et al., 2013 | G1130R | EC11 | i | Non-syndromic |
| Yang et al., 2013 | S1267P | EC12 | o | Non-syndromic |

**Table S8.** Summary of simulations.

| Label | System | $t_{sim}$ (ns) | Type | Start | Speed (nm/ns) | Average Peak Force (pN)[b] | Size (#atoms) | Initial Size ($nm^3$) |
|---|---|---|---|---|---|---|---|---|
| S1a | *Hs* | 11.1 | EQ[a] | – | – | – | 924,024 | 71.2 × 9.2 × 14.9 |
| S1b | heterodimer | 1.2 | SMD[c] | S1a | 10 | 811.6 | | |
| S1c | | 11.2 | SMD[c] | S1a | 1 | 447.2 | | |
| S1d | | 191.9 | SMD[c] | S1a | 0.1 | 331.2 | | |
| S2a | *Mm* | 11.1 | EQ[a] | – | – | – | 1,016,888 | 91.7 × 11.1 × 10.5 |
| S2b | heterodimer | 3.3 | SMD[c] | S2a | 10 | 751.9 | | |
| S2c | | 30.2 | SMD[c] | S2a | 1 | 398.8 | | |
| S2d | | 251.6 | SMD[c] | S2a | 0.1 | 355.9 | | |
| S3a | *Hs* short | 11.1 | EQ[a] | – | – | – | 833,064 | 40.5 × 17.3 × 12.4 |
| S3b | tetramer | 2.1 | SMD[d] | S3a | 10 | 2195.3 | | |
| S3c | 3 $Ca^{2+}$ / | 12.3 | SMD[d] | S3a | 1 | 1288.8 | | |
| S3d | linker | 95.6 | SMD[d] | S3a | 0.1 | 1082.5 | | |
| S4a | *Hs* short | 10.0 | EQ[a] | – | – | – | 833,037 | 40.5 × 17.3 × 12.4 |
| S4b | tetramer | 1.9 | SMD[d] | S4a | 10 | 1940.2 | | |
| S4c | 2 $Ca^{2+}$ / | 13.7 | SMD[d] | S4a | 1 | 1175.8 | | |
| S4d | linker | 99.5 | SMD[d] | S4a | 0.1 | 737.7 | | |
| S5a | *Hs* short | 10.0 | EQ[a] | – | – | – | 832,984 | 40.5 × 17.3 × 12.4 |
| S5b | tetramer | 2.3 | SMD[d] | S5a | 10 | 2084.7 | | |
| S5c | 1 $Ca^{2+}$ / | 15.1 | SMD[d] | S5a | 1 | 1173.3 | | |
| S5d | linker | 129.8 | SMD[d] | S5a | 0.1 | 833.4 | | |
| S6a | *Hs* short | 10.0 | EQ[a] | – | – | – | 832,924 | 40.5 × 17.3 × 12.4 |
| S6b | tetramer | 1.9 | SMD[d] | S6a | 10 | 1846.3 | | |
| S6c | 0 $Ca^{2+}$ / | 14.5 | SMD[d] | S6a | 1 | 1067.2 | | |
| S6d | linker | 142.9 | SMD[d] | S6a | 0.1 | 760.7 | | |
| S7a | *Mm* short | 11.1 | EQ[a] | – | – | – | 743,064 | 51.1 × 11.7 × 12.9 |
| S7b | tetramer | 3.3 | SMD[d] | S7a | 10 | 2064.9 | | |
| S7c | 3 $Ca^{2+}$ / | 11.4 | SMD[d] | S7a | 1 | 1003.0 | | |
| S7d | linker | 88.3 | SMD[d] | S7a | 0.1 | 804.5 | | |
| S8a | *Hs* long | 11.1 | EQ[a] | – | – | – | 1,178,985 | 99.4 × 11.4 × 10.9 |
| S8b | tetramer | 1.7 | SMD[d] | S8a | 10 | 1417.3 | | |
| S8c | 3 $Ca^{2+}$ / | 18.1 | SMD[d] | S8a | 1 | 893.6 | | |
| S8d | linker | 122.3 | SMD[d] | S8a | 0.1 | 510.4 | | |
| S9a | *Mm* long | 11.1 | EQ[a] | – | – | – | 1,292,017 | 91.5 × 11.8 × 12.5 |
| S9b | tetramer | 3.4 | SMD[d] | S9a | 10 | 1484.7 | | |
| S9c | 3 $Ca^{2+}$ / | 20.6 | SMD[d] | S9a | 1 | 852.6 | | |
| S9d | linker | 135.5 | SMD[d] | S9a | 0.1 | 630.3 | | |
| S10a | | 95.3 | EQ[a] | – | – | – | 2,325,962 | 65.5 × 18.9 × 19.5 |
| S11a | *Mm* | 11.1 | EQ[a] | – | – | – | 456,282 | 27.9 × 12.9 × 12.9 |
| S11b | PCDH15 | 3.0 | SMD[d] | S11a | 10 | 1418.5 | | |
| S11c | EC9-12 | 14.9 | SMD[d] | S11a | 1 | 882.8 | | |
| S11d | dimer | 95.8 | SMD[d] | S11a | 0.1 | 604.2 | | |
| **Total** | | 1,742.3 | | | | | | |

[a] EQ indicates simulations that consisted of 1,000 steps of minimization, 100 ps of dynamics with the protein backbone constrained ($k$ = 1 kcal/mol/Å$^2$), 1 ns of free dynamics in the *NpT* ensemble ($\gamma$ = 1 ps$^{-1}$), and 10 ns of free dynamics in the *NpT* ensemble ($\gamma$ = 0.1 ps$^{-1}$).

[b] Average peak force is calculated from the peak force measured on stretched Cα atoms or slabs (using 50-ps running averages).

[c] SMD simulation in which terminal Cα atoms where attached to independent stretching springs.

[d] SMD simulation in which force was applied through slabs to protein ends (see Methods).
